# Ultra-precision deconvolution of spatial transcriptomics decodes immune heterogeneity and fate-defining programs in tissues

**DOI:** 10.1101/2024.07.05.602200

**Authors:** Yin Xu, Zurui Huang, Yawei Zhang, Minghui Gong, Zhenghang Wang, Peijin Guo, Feifan Zhang, Jing Yang, Guanghao Liang, Lihui Dong, Renbao Chang, Yu Xia, Haochen Ni, Wenxuan Gong, Boyuan Mei, Yuan Gao, Zhaoqi Liu, Lin Shen, Jian Li, Meng Michelle Xu, Dali Han

## Abstract

Immune cells infiltrate tissues in response to exogenous pathogens or spontaneous tumors, generating protective immunity to safeguard tissues. Despite advancements in spatial transcriptomics (ST), the precise spatial distribution and functional specialization of immune cells within tissue microenvironments remain elusive. Here, we introduce an ultra-precision ST deconvolution algorithm (UCASpatial) that enhances the mapping of cell subpopulations to spatial locations by leveraging the contribution of genes indicative of cell identity through entropy-based weighting. Using both in silico and real ST datasets, we demonstrate that UCASpatial improves the robustness and accuracy in identifying low-abundant cell subpopulations and distinguishing transcriptionally heterogeneous cell subpopulations. Applying UCASpatial to human colorectal cancer (CRC), we link genomic alterations in individual cancer clones to multi-cellular characteristics of the tumor immune microenvironment (TIME) and reveal the co-evolution of tumor cells and TIME at a clonal resolution. We show that the copy number gain on chromosome 20q (chr20q-gain) in tumor cells orchestrates a T cell-excluded TIME, indicative of resistance to immunotherapy in CRC, and is associated with tumor-intrinsic human endogenous retrovirus subfamily H (*HERV-H*) silencing and impaired type I interferon response. In murine wound healing models, we illuminate the spatiotemporal dynamics of individual cell subsets across various stages of the healing process. We discovered that the scarring-healing mice (C57BL/6) exhibited a replacement of *Prg4*^+^ chondrogenic progenitors to *Igfbp5*^+^ chondrocytes in the wound bed at the regeneration stage, a change not observed in the regenerative strain (MRL/MpJ). The *Igfbp5*^+^ chondrocyte, spatially coordinating with *Cd36^+^ Gpnmb^+^ Il1b^-^* macrophage and *Fmod*^+^ fibroblast, forms a pro-fibrotic community associated with regeneration failure. The cell-cell interactions within this three-cell cluster, mediated by the IL11-IL11RA axis, drive the pro-fibrotic community formation and limit regeneration in C57BL/6 mice. Our findings present UCASpatial as a versatile tool for deciphering fine-grained cellular landscapes in ST and exploring intercellular mechanisms in complex and dynamic microenvironments.

## Introduction

The ability to elucidate the spatial organization of cells within the microenvironment is crucial for understanding cellular functions and exploring intercellular mechanisms that govern development, physiology, and pathogenesis ^1, 2^. Emerging spatial transcriptomics (ST) technologies offer promising avenues for achieving this; however, current spot-based techniques face a significant limitation in lacking single-cell resolution. Methods like Visium ^3^, Tomo-Seq ^4^, Slide-seq ^5^, and HDST ^6^ amalgamate RNA-seq measurements from multiple cells or cell fractions at each spot. As a result, these techniques struggle with the precise identification of individual cell types within the microenvironment. Consequently, annotating cell types at each spot becomes imperative for accurately capturing spatial cell organization ^7, 8^.

To address this challenge, various computational approaches have been proposed to deconvolute spatial transcriptomic profiles into cellular compositions based on gene expression profiles of each cell type identified through single-cell RNA sequencing (scRNA-seq), such as SPOTlight ^9^, RCTD ^10^, cell2location ^11^, CARD ^12^, stereoscope ^13^, and Spotiphy ^14^. These methods effectively delineate significant differences in cell type compositions across diverse anatomical or pathological layers in well-structured tissues such as the brain, heart, and lymphatic structures, even at fine-grained resolutions ^11, 12, 15-17^.

In recent years, a growing body of research has emphasized the importance of characterizing the complex microenvironment, particularly the influx of immune cells that orchestrate dynamic spatial patterns, varying between tissue homeostasis and non-homeostasis conditions. This complexity is evident in scenarios such as tissue damage ^18, 19^ and the tumor microenvironment ^20-24^. However, several immune cells, particularly those infiltrating into tumors, are known for their sparse presence and low transcriptional activity ^25-28^. Their discontinuous spatial distributions in these microenvironments limit the effectiveness of algorithms relying on spot-neighbor information ^28, 29^. Additionally, the low abundance and the context-dependent phenotypic variation of heterogeneous immune cells, particularly T cells, pose challenges for conventional deconvolution algorithms ^25^. Furthermore, their scarcity makes them difficult to capture accurately using histological images, thus constraining the applicability of image-based algorithms ^30, 31^. Consequently, the precise characterization of complex and dynamic microenvironments remains a significant challenge.

To address the aforementioned challenge, we developed UCASpatial, an ultra-precision cellular deconvolution algorithm for spatial transcriptomics based on entropy-weighted regression. A unique advancement of UCASpatial is its formulation of the Shannon entropy-based weights to quantify the ability of each gene for indicating cell identity and incorporate this information in the deconvolution process. This approach improves the robustness and accuracy in identifying low-abundant cell subpopulations and distinguishing transcriptionally heterogeneous cell subpopulations in ST, thus allowing for more precise and unbiased deconvolution of the cellular landscape.

By applying UCASpatial to human colorectal cancer (CRC), UCASpatial has delineated the spatial patterns of immune cells within the tumor immune microenvironment (TIME). By integrating the fine-grained TIME landscape with spatially inferred copy number variations (CNVs), we characterize two functional multicellular communities associated with T cell-rich and T cell-excluded TIMEs at a clonal resolution. Further, we demonstrate that the chr20q-gain in CRC tumor cells is associated with reduced expression of the tumor-intrinsic human endogenous retrovirus subfamily H (*HERV-H*) and impaired type I interferon response, which orchestrates the T cell-excluded TIME phenotype. In the ear wound healing models of both regenerative MRL/MpJ and scarring-healing C57BL/6 mice (B6 mice), we illustrate the spatiotemporal dynamics of immune cell subpopulations involved in the healing process and unveil the spatial segregation of macrophage subpopulations embedded within distinct cellular communities at the wound sites. Moreover, we identify a previously uncharacterized *Igfbp5*^+^ chondrocyte subpopulation that coordinates with *Cd36^+^ Gpnmb^+^ Il1b^-^* macrophage and *Fmod*^+^ fibroblast at the wound bed, establishing a three-cell-type community with pro-fibrotic functions linked to scarring in B6 mice. By blocking the *Fmod*^+^ fibroblast differentiation with the PPARγ agonist rosiglitazone, or by inhibiting the IL11-IL11RA axis within the community in B6 mice, we observed a reduction in the formation of the pro-fibrotic community and an improvement in the regenerative ability of ear wounds in B6 mice.

## Results

### UCASpatial: an entropy-weighted regression model for spatial deconvolution of cellular landscapes

UCASpatial deconvolutes ST data into cellular compositions by leveraging cell identity-specific gene expression profiles (cGEPs) derived from either paired or unpaired scRNA-seq data (Fig. 1). First, UCASpatial extracts cGEPs from the referenced scRNA-seq dataset with a set of user-provided cell-subpopulation labels (fig. S1, A to C). It strategically selects cells meeting a meta-purity threshold to enhance discrimination of referenced cell identities, and then extracts cGEPs through optimized Non-negative Matrix Factorization (NMF). Second, UCASpatial formulates Shannon entropy-based weights for each gene within the extracted cGEPs to enhance genes indicative of cell identities (Methods). Third, UCASpatial employs Weighted Non-Negative Least Squares (WNNLS) to deconvolute the mRNA counts of each spot in the ST data using the weighted cGEPs, thereby estimating the cellular compositions. Additionally, UCASpatial offers a suite of downstream analysis tools. These include identifying tumor clones and their corresponding tumor immune microenvironment (TIME), as well as spatially resolved quantification of transposable elements (TEs). To ensure versatility in applications, UCASpatial is seamlessly integrated with the Seurat framework.

**Fig. 1.**
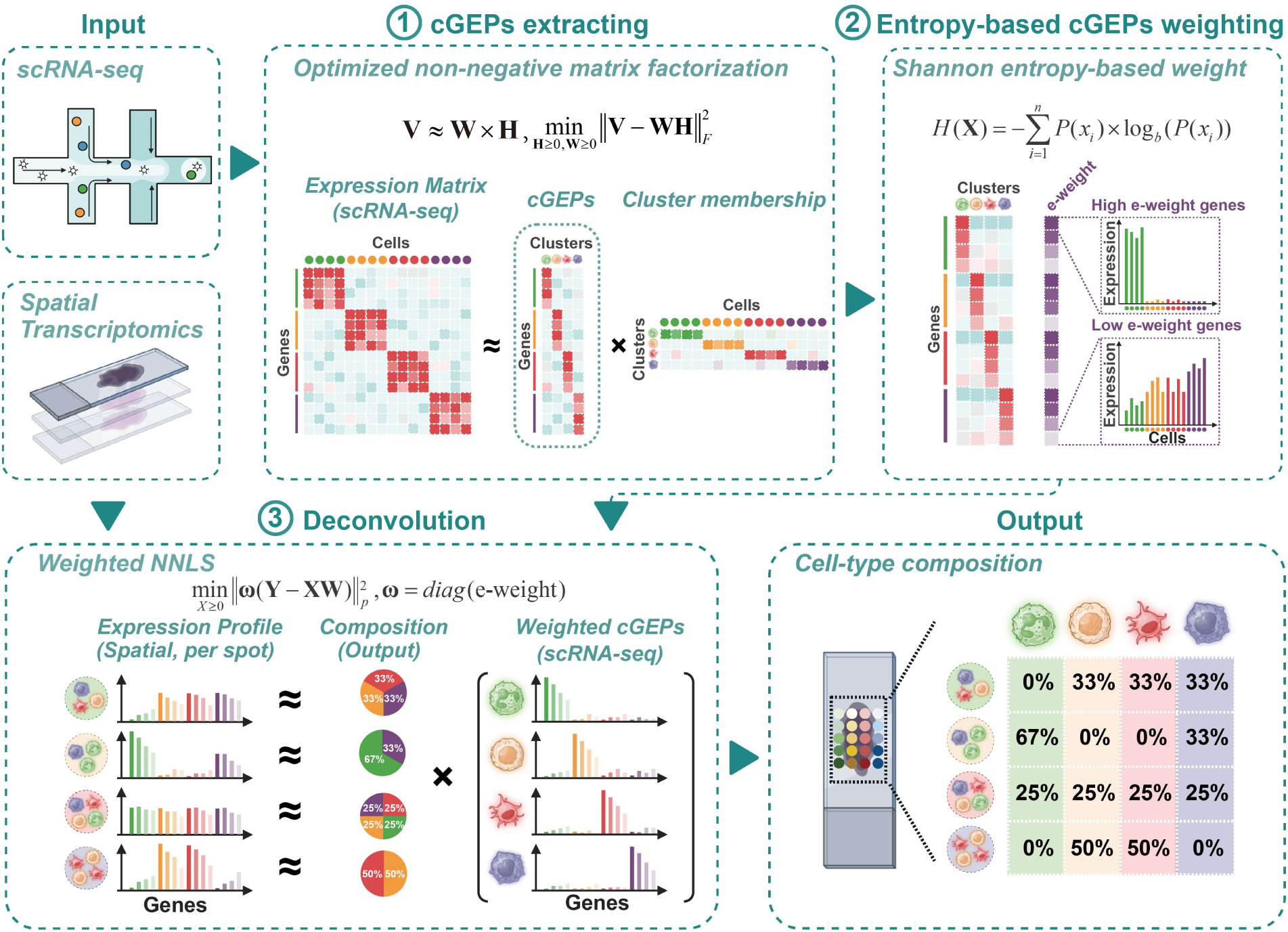
Schematic overview of UCASpatial. UCASpatial takes both the referenced single-cell RNA sequencing (scRNA-seq) data with cluster information and spatial transcriptomic data as input. After three-step processing involving cell-identity-specific GEPs (cGEPs) extraction, entropy-based cGEPs weighting, and deconvolution, UCASpatial decomposes spatially resolved, multi-cell RNA count matrices (i.e., spatial transcriptomic data) into a referenced cell-type composition. GEP, gene expression profile. e-weight, Shannon entropy-based weight. NNLS, non-negative least squares. Figure adapted from images created with BioRender.com.

### Benchmarking the performance of UCASpatial in the simulated ST dataset

To assess UCASpatial’s performance, we generated a simulated ST dataset comprising 750 ST spots designed to mimic the real cellular composition of the tumor microenvironment. Each spot was constituted by the transcripts of 5 to 20 cells sampled from scRNA-seq data of human CRC, encompassing 20 reference cell subpopulations (fig. S1, A and B, and Methods). The cellular composition of simulated spots was designed to recapitulate the spatially correlated cell-type distribution patterns observed in tumor microenvironments, particularly the relationship between epithelial, stromal, and immune cell frequencies across tumor core, margin, and stromal regions (fig. S1D and table S1) ^17, 32^.

To assess the performance of UCASpatial on the simulated ST dataset with varying complexity, we ranked the complexity of these simulated spots based on their cellular composition entropy. Subsequently, we categorized the simulated spots into three groups (low, medium, and high complexity), where higher entropy denotes increased complexity (Fig. 2A and fig. S1D). We compared UCASpatial with several recently established methods, including SPOTlight, RCTD, cell2location, CARD, stereoscope, and Spotiphy, using Root Mean Square Error (RMSE) to first quantify the deviation between estimations and ground truth at the spot level (Methods). UCASpatial exhibited significantly superior performance across all complexity levels (median RMSE = 0.051) compared to the aforementioned methods, with 10%, 21%, 31%, 31%, 36%, and 35% improvement in terms of RMSE compared to RCTD (0.058), SPOTlight (0.066), CARD (0.075), Spotiphy (0.075), cell2location (0.081), and stereoscope (0.080), respectively (Fig. 2B and fig. S1E). Additionally, UCASpatial demonstrated a comparable Pearson Correlation Coefficient (PCC) to cell2location and significantly outperformed other methods, when comparing estimations with the ground truth (fig. S1F). Moreover, we also assessed the ability for identifying the existence of cell subpopulations within spots using the F1 score. Consistently, UCASpatial significantly outperformed other methods across all complexity levels with the highest F1 score (with a 6-25% improvement) (Fig. 2B and fig. S1G).

**Figure 2.**
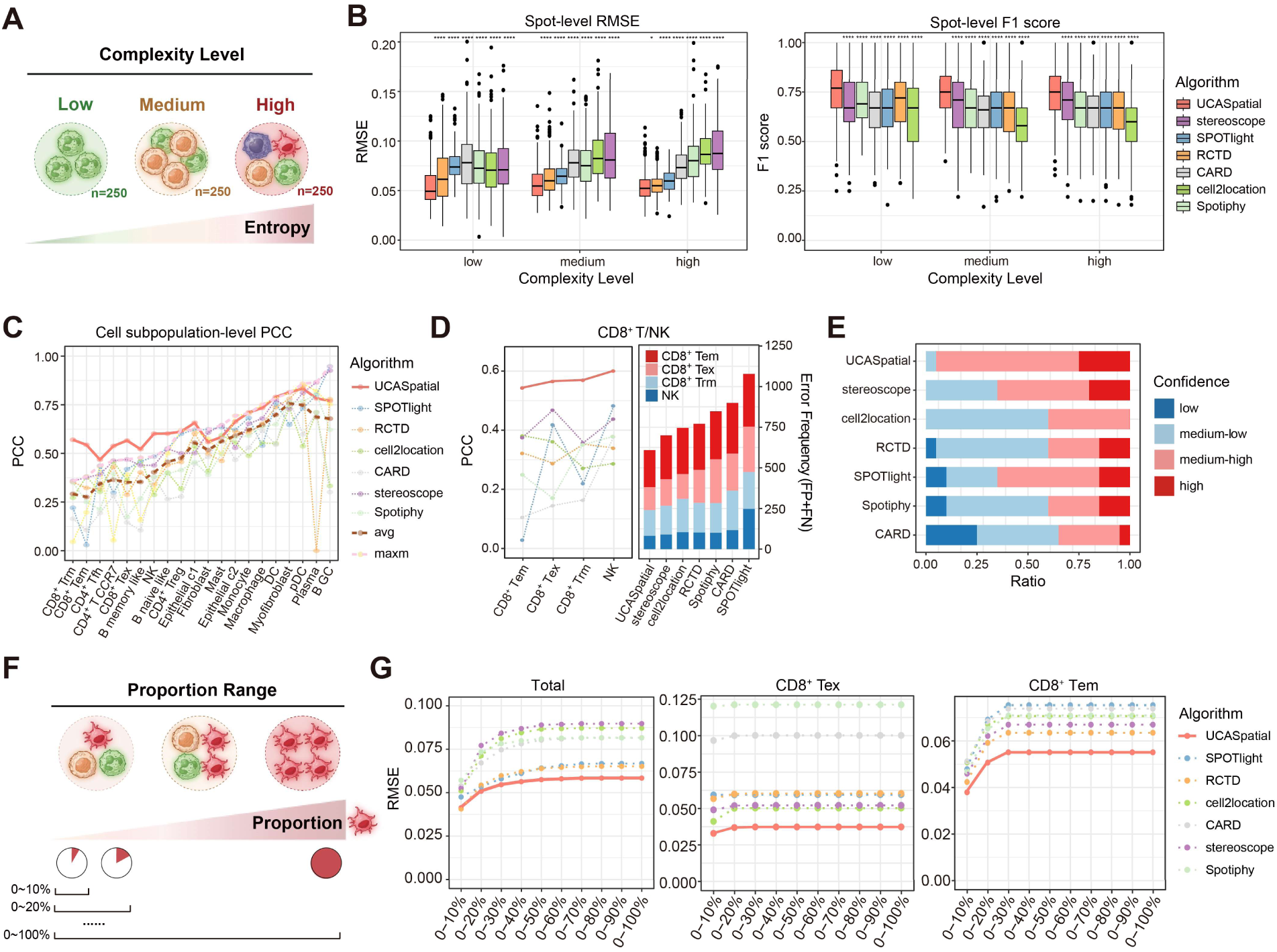
Validation of UCASpatial using simulated data. (**A**) Graphic model showing the classification of 750 simulated spots into three groups, each representing a different complexity level - low, medium, and high, respectively. The classification is based on the entropy of cell-type composition for each spot, with each group containing 250 spots. (**B**) Boxplot showing spot-level root mean square error (RMSE) (left) and F1 score (right) for each method across three complexity levels. Boxplot elements are defined as: center line, median; box limits, upper and lower quartiles; whiskers, 1.5× interquartile range; points, outliers. Statistical analysis was performed using a two-tailed paired t-test. ns, non-significant, p > 0.05; *0.01 ≤ p < 0.05; **p < 0.01; ***p < 0.001; ****p < 0.0001. (**C**) Line charts showing cell-subpopulation-level Pearson correlation coefficient (PCC) for each method across 20 cell subpopulations. ‘avg’ and ‘maxm’ represent the average and maximum PCC, respectively, of the six methods excluding UCASpatial. (**D**) Left, line charts representing cell-subpopulation-level PCC of CD8^+^ effector memory T cells (CD8^+^ Tem), CD8^+^ exhausted T cells (CD8^+^ Tex), CD8^+^ tissue-resident memory T cells (CD8^+^ Trm), and natural killer cells (NK) for each method. Right, stacked bar plot showing the accumulated count of false predictions of these cell subpopulations for each method. Colors represent different cell subpopulations. FP, false positive; FN, false negative; FP + FN, total error frequency. (**E**) The cell-subpopulation-level PCC values for each method were binned into four confidence groups: 0-0.25 (low), 0.25-0.5 (medium-low), 0.5-0.75 (medium-high), and 0.75-1 (high). Stacked bar plot showing the ratio of four confidence groups for each method. (**F**) Graphic model representing the classification based on the proportion range of a specific cell subpopulation within the simulated spots. All simulated spots were classified into ten groups from proportion 0-10% to proportion 0-100% for each cell subpopulation. (**G**) Left, line charts representing the average RMSE of 20 cell subpopulations across various proportions for each method. Middle, line charts showing an example of the RMSE for CD8^+^ Tex across different proportions. Right, line charts show an example of the RMSE for CD8^+^ Tem across different proportions. Each dot in the line charts represents the average RMSE from multiple simulations. Figures (**A** and **F**) adapted from images created with BioRender.com.

Additionally, to assess the robustness of UCASpatial across different cell numbers, we categorized spots into three groups based on the number of cells per spot: 5-9 cells, 10-12 cells, and 13-20 cells (fig. S1H). Our analysis revealed that UCASpatial maintained robust performance across these varying cell numbers, whereas the performance of cell2location and stereoscope dropped as cell number increased (fig. S1I). These results highlight UCASpatial’s robustness in handling diverse cell densities, thereby confirming its adaptability and reliability.

We next evaluated the precision and robustness of UCASpatial in deconvoluting different cell subpopulations. Through in-silico analyses using three distinct reference scRNA-seq labels with varying numbers of subpopulations (7, 11, and 20), we observed that the advantages of UCASpatial became more pronounced as the number of subpopulations increased compared with other methods (fig. S1, J and K). With 20 subpopulations, UCASpatial exhibited superior performance across all metrics, including RMSE, PCC, and F1 score, highlighting its proficiency in deciphering fine-grained spatial landscapes (fig. S1K). Specifically, examination of PCC between estimations and ground truth for each method under 20 subpopulations condition unveiled UCASpatial’s superior performance, achieving the highest PCC in 15 out of 20 cell subpopulations (Fig. 2C), particularly notable for NK cell and CD8^+^ T cell subpopulations (with a 39-187% improvement) (Fig. 2D and fig. S1L). By categorizing the PCC values of each cell subpopulation into four groups, we noticed UCASpatial exhibited PCC values surpassing the medium-high correlation level (PCC > 0.5) for 19 out of 20 cell subpopulations, with the sole exception CD4^+^ Tfh (PCC = 0.47) while remaining the best compared to alternative methods (Fig. 2, C and E). In contrast, other methods yielded PCC values below the medium-high correlation level for at least 30% of cell subpopulations, with some even falling below 0.25 (Fig. 2E and fig. S1M). This comparison underscored UCASpatial’s capability in discriminating cell subpopulations, even with subtle transcriptomic differences (fig. S1N). Furthermore, assessments using the PCC and F1 score reaffirmed UCASpatial’s precise and robust decomposition across each cell subpopulation (fig. S1, O and P).

Considering the challenge of ST analysis in precisely mapping the spatial location of low-abundant cell subpopulations, we next investigated UCASpatial’s sensitivity in annotating cell subpopulations with low frequencies within spots. We categorized the proportion of each cell subpopulation within simulated spots into ten groups, forming a gradient that spanned from 0-10% to 100% (Fig. 2F). Subsequently, we calculated the RMSE across various proportions to evaluate the deviation of cell-subpopulation proportions. We observed that UCASpatial exhibited the lowest overall RMSE (averaged by each cell subpopulation, medium RMSE = 0.051) across different proportions with a 7-31% improvement over the other methods (Fig. 2G). Moreover, we specifically examined the RMSE of different cell subpopulations, revealing that UCASpatial notably outperformed other methods for cell subpopulations with subtle transcriptomic differences such as CD8^+^ Tex and CD8^+^ Tem (with 25-63% and 13-26% improvements, respectively) (Fig. 2G and fig. S1Q). Consistent results were observed in assessments using the PCC and F1 score (fig. S1R), highlighting UCASpatial’s superiority in accurately annotating cell subpopulations with low frequencies within spots, even with subtle transcriptomic differences.

Furthermore, we evaluated the performance enhancement afforded by UCASpatial’s entropy-based weighting and meta-purity filtering designs (fig. S1, S to U). We confirmed that the meta-purity filtering process did not result in the loss of any predefined cell subpopulations, with PCC values before and after filtering consistently exceeding 0.9 (fig. S1, V to X). We found that the entropy-based weighting design showed improvements of 22%, 50%, and 17% in terms of RMSE, PCC, and F1 score, respectively, compared to UCASpatial without weighting. The meta-purity filtering design exhibited improvements across different parameters with 4-11%, 1-8%, and 1-3% in terms of RMSE, PCC, and F1 score. In addition, the same analyses as before showed that UCASpatial, without entropy-weighting and meta-purity filtering, lost the aforementioned improvements (fig. S1, E to R).

We evaluated the runtime and memory usage of UCASpatial and other methods (Methods), analyzing the impact of the number of spots in spatial data and the number of cells in scRNA-seq data on computational resources. UCASpatial showed stable time costs, processing data in under 1 hour as the number of spots increased (fig. S1Y). Cell2location encountered memory errors on our GPU platform with the largest dataset (50,000 spots and ∼12,500 cells). Other methods, except stereoscope, were minimally affected by the scRNA-seq reference size (fig. S1Y). Using a simulated dataset (50,000 spots, ∼12,500 cells, 20 cell subpopulations), UCASpatial demonstrated the lowest memory usage (< 26GB) and highest efficiency (fig. S1Z). Collectively, our in-silico benchmarking demonstrates that UCASpatial enables precise and sensitive spatial deconvolution, excelling in discerning cell subpopulations with subtle transcriptomic differences, even in low proportions.

### UCASpatial demonstrates broad applicability and high precision across multi-resolution platforms

To test whether UCASpatial is applicable across platforms with varying resolutions, we applied UCASpatial to 10x Visium HD data from a colorectal cancer (CRC) sample ^33^, using the recommended bin size of 8 μm × 8 μm (fig. S2A and Methods). We compared UCASpatial with RCTD, a method also suitable for processing 10x Visium HD data using PCC between estimated proportions of cell subpopulations and marker-gene scores in the absence of ground truth ^16^. We observed that UCASpatial achieved the highest PCC, and exhibited over 20 times faster than RCTD on the runtime evaluation (fig. S2, B to D, Methods).

We also analyzed Slide-seq V2 data from the mouse hippocampus to evaluate UCASpatial’s performance at single-cell resolution (fig. S2, E to H). Since strictly defined ground-truth cell type labels are not available experimentally, we adopted the widely accepted evaluation strategy established by the CARD paper^12^. This involves verifying whether the deconvolved cell types accurately reconstruct known anatomical structures (e.g., CA1, CA3, Dentate Gyrus). Specifically, we visualized the predicted top 1 cell type for each location (fig. S2F) and quantitatively evaluated the concordance between method-predicted cell-type proportions and known region-specific marker expression in three anatomical structures^12^ by calculating PCC (fig. S2G). Our results show that UCASpatial is highly concordant with other methods, successfully captures key anatomical structures, and confirms a comparable level of accuracy and spatial resolution on this neurobiological dataset (fig. S2, G and H).

Together, these results demonstrate UCASpatial’s broad applicability and high precision across platforms with varying resolutions, from bin resolution (10x Visium HD) to single-cell resolution (Slide-seq V2).

### Benchmarking the performance of UCASpatial using Visium HD-derived pseudo-spots

To obtain synthetic data that better capture spatial correlation structures between neighboring spots and cell-subpopulation-specific capture efficiency variations, we generated synthetic spots by spatially aggregating segmented single-cell transcriptomes from three 10x Genomics Visium HD colorectal cancer tissue sections^37^ into 49 µm × 49 µm bins (fig. S2, I to K), closely matching standard Visium spot size. This approach preserves local spatial correlations and yields 39,813 pseudo-spots with ground-truth cell-subpopulation compositions reflecting the tumor microenvironment’s complexity.

Benchmarking across this dataset recapitulated prior performance trends. UCASpatial achieved the highest average PCC (0.87) and F1 score (0.82), alongside the lowest RMSE (0.05) among other methods at the spot level (fig. S2L), even under scenarios of varying complexity (fig. S2M). Cell-subpopulation-level evaluation further confirmed UCASpatial’s superior accuracy in deciphering different cell subpopulations (fig. S2, L and N). This data further supports UCASpatial’s accurate and stable performance on biologically realistic synthetic data.

### UCASpatial accurately annotates cell subpopulations in human CRC TIMEs

Spatial omics has provided a powerful tool for dissecting the intricate interplays between tumor alterations and the co-evolving immune microenvironment ^22, 38, 39^. However, the characterization of immune cell compositions, particularly those infiltrating into distinct tumor clone regions, remains a formidable challenge due to their low abundance and limited transcriptional output ^22, 25, 27, 28, 40^. To investigate whether UCASpatial can precisely resolve the tumor immune microenvironment (TIME), we curated an ST dataset from eight tumor slices of seven patients with human CRC using the 10x Genomics Visium platform as the discovery cohort ^41^ (Fig. 3A, fig. S3A, and table S2). We leveraged an integrated scRNA-seq dataset comprising an in-house scRNA-seq with three patients, and a public scRNA-seq dataset with five patients (P1-5) ^41^, encompassing 34 cell subpopulations as a reference (Fig. 3, A and B, fig. S3, B to D, and Methods).

**Fig. 3.**
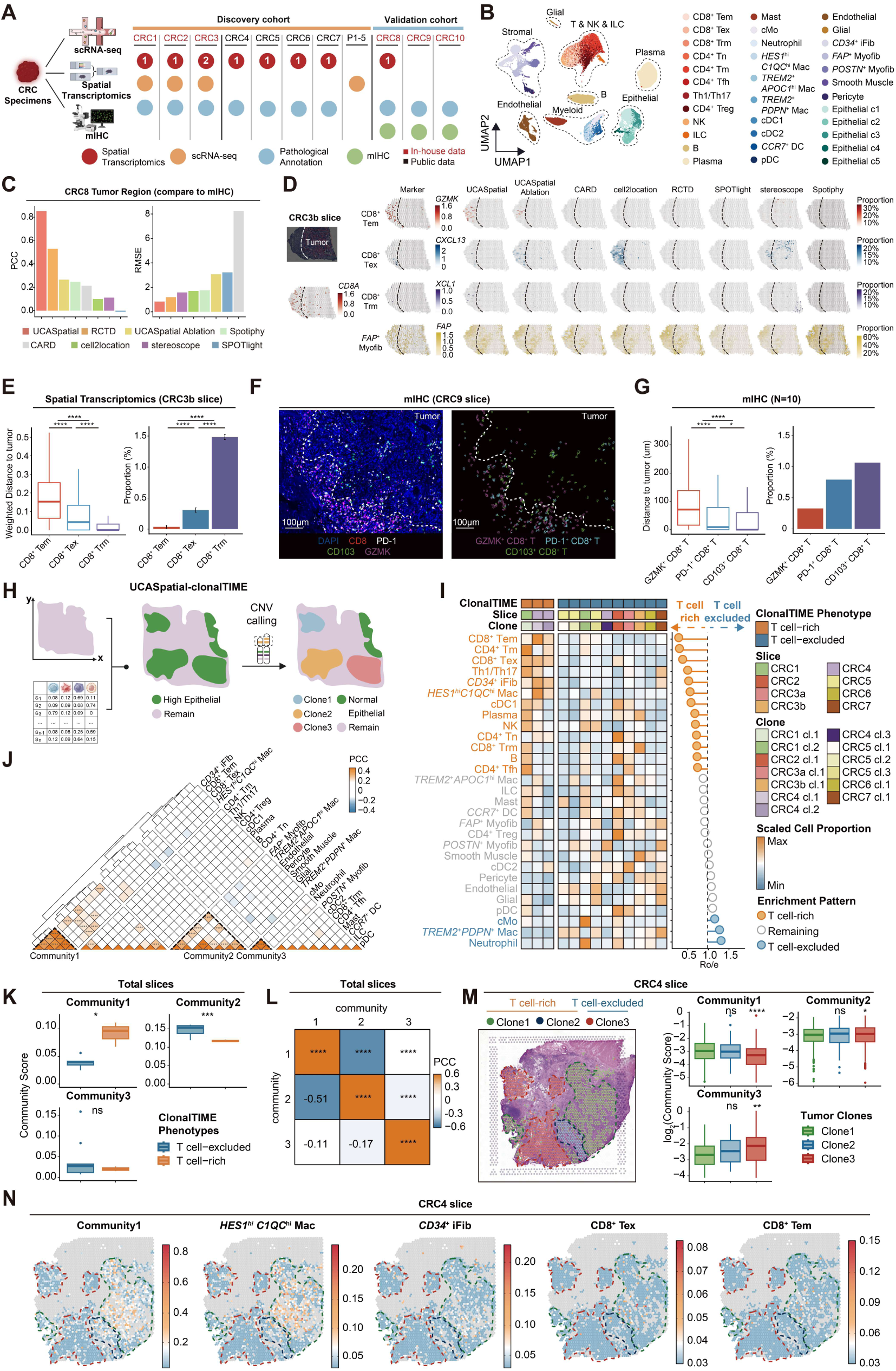
Characteristics of clonal-resolution landscape with two TIME phenotypes in human CRC. (**A**) Left, experimental design of in-house colorectal cancer (CRC) dataset. Right, an overview of the CRC dataset used for UCASpatial analysis, including a discovery cohort and a validation cohort, with samples organized by resource (indicated by text color). The red, orange, blue, and green dots represent the presence of spatial transcriptomics, scRNA-seq, pathological annotation, and mIHC, respectively. The number in red dots indicates the count of spatial transcriptomic slices per patient. (**B**) UMAP plot displaying 34 cell subpopulations identified from scRNA-seq reference. (**C**) Bar plots displaying PCC (left) and RMSE (right) were used to compare each method to the mIHC results in the tumor regions of the CRC8 slice. (**D**) Left, the pathological annotations and indicating markers’ expression (*CD8A*, *GZMK*, *CXCL13*, *XCL1*, and *FAP*) in the CRC3b slice. Colors represent the expression level of each gene. Right, spatial visualization of the distribution of indicated CD8^+^ T cell subpopulations at the CRC3b slice based on UCASpatial, UCASpatial without entropy-based weight and meta-purity filter (UCASpatial Ablation), CARD, cell2location, RCTD, SPOTlight, and stereoscope. Colors represent the proportion of each CD8^+^ T cell subpopulation. The dashed line indicates the tumor-invasive front. (**E**) Left, boxplot showing the distance between CD8^+^ T cell subpopulations and tumor region in the CRC3b slice. Right, bar plot showing the proportion of CD8^+^ T cell subpopulations in the tumor region within the CRC3b slice. (**F**) Left, representative immunofluorescence imaging of CRC tissue (CRC9). The staining images indicate DAPI (blue), CD8 (red), PD-1 (white), CD103 (green), and GZMK (magenta). Right, the distribution of CD8^+^ T cell subpopulations within CRC9. The cyan, green, and magenta colors indicate PD-1^+^ CD8^+^ T cells, CD103^+^ CD8^+^ T cells, and GZMK^+^ CD8^+^ T cells, respectively. The dashed white lines indicate the tumor-invasive front. Scale bar, 100 μm. (**G**) Left, boxplot showing the distance between CD8^+^ T cell subpopulations and tumor regions in 10 ROIs. Right, bar plot showing the proportion of CD8^+^ T cell subpopulations within the tumor region in 10 ROIs. ROI, region of interest. (**H**) Schematic representation of UCASpatial-clonalTIME platform. From left to right: identification of epithelial regions based on UCASpatial; delineation of tumor clone regions and normal epithelial regions using spatialCNV (Methods). clonalTIME, clonal tumor immune microenvironment. (**I**) Heatmap showing the distribution of cell subpopulations across clones grouped by clonalTIME phenotypes. Colors reflect scaled cell subpopulation proportions estimated by UCASpatial. Lollipop plot displaying the Ro/e scores across all clones. Ro/e scores represent the enrichment pattern for each cell subpopulation between 2 clonalTIME phenotypes. Columns in the heatmap were annotated by cloneID, sliceID, and clonalTIME phenotype. Ro/e, the ratio of observed over expected cell subpopulation proportions. (**J**) Heatmap showing the PCC between the proportion of each two cell subpopulations across all spots after neighborhood processing within tumor regions (Methods). Communities were identified using hierarchical clustering. (**K**) Boxplot showing the score of each community between T cell-excluded and T cell-rich clonalTIMEs at clonal-resolution. (**L**) Heatmap displaying the PCC between the community scores of spots within tumor regions. Colors represent the PCC value. (**M**) Left, the spatial distribution of each clone in the CRC4 slice. Spots filled with green, blue, and red represent the tumor regions of clone1, clone2, and clone3, respectively. Dashed lines mark the general contour of each tumor clone region. Right, boxplots showing the score of each community in each tumor clone of CRC4. (**N**) Spatial visualization of the score of community1, and the proportion of cell subpopulations involved in community1 (comprising *HES1*^hi^ *C1QC*^hi^ macrophage, *CD34*^+^ iFib, CD8^+^ Tex, and CD8^+^ Tem) in the tumor regions of the CRC4 slice. Non-tumor regions were depicted in light grey for clarity. Figures (**A** and **H**) adapted from images created with BioRender.com. In Figures (**E** and **G**), statistical analysis was performed using a Wilcoxon rank-sum test. In Figures (**J**, **K**, **L**, and **M**), statistical analysis was performed using a two-tailed unpaired *t*-test. ns, non-significant, p > 0.05; *0.01 ≤ p < 0.05; **p < 0.01; ***p < 0.001; ****p < 0.0001. Boxplot elements in Figures (**E**, **G**, **K**, and **M**) are defined as: center line, median; box limits, upper and lower quartiles; whiskers, 1.5× interquartile range; points, outliers.

We annotated the spatial distribution of 34 cell subpopulations in human CRC ST discovery cohort using UCASpatial. Notably, stromal cell subpopulations showed consistent distribution with anatomical locations as expected from prior literature ^42, 43^, such as smooth muscle cells of the muscular layer and endothelial cells around blood vessels, along with pericytes (fig. S3E). Similarly, epithelial cell subpopulations displayed enrichment in the epithelial region, consistent with corresponding H&E staining and pathological annotations (fig. S3F). Upon further examination of immune subpopulations, we noticed B cell and various T cell and dendritic cell (DC) subpopulations, including CD4^+^ Tn, CD4^+^ Tfh, CD8^+^ Tex, CD8^+^ Tem, cDC2, and *CCR7^+^ DC*, formed a multicellular community resembling a tertiary lymphoid structure (TLS) (fig. S3G, see table S3 for abbreviation information) ^44, 45^. Additionally, we noticed the spatial exclusivity of *HES1*^hi^ *C1QC*^hi^ macrophages and *TREM2*^+^ *PDPN*^+^ macrophages (fig. S3H), indicating their distinct functions in TIMEs.

For comparative analysis, we also annotated this ST dataset using other methods. Remarkably, UCASpatial achieved the highest average PCC with a significant improvement of 17-104% over other methods between the cell-subpopulation proportions and signature expressions, especially in T cell subpopulations (with at least a 26% improvement) (fig. S3, I and J, table S4, and Methods). In alignment with in-silico benchmarking results, all evaluated methods correctly annotated structural cell subpopulations, for example, *FAP*^+^ myofibroblast (fig. S3J), yet UCASpatial showed a higher sensitivity in annotating immune cell subpopulations (fig. S3, I and J). For instance, UCASpatial performed better than other methods in identifying the spatial distribution of T cell subpopulations in almost all five TLS regions (fig. S3K). While UCASpatial depicted 4-6 T cell subpopulations consistent with their marker gene expressions, all compared methods only identified 0-2 T cell subpopulations (fig. S3L).

To assess the accuracy of immune cell annotation in the real dataset, we constructed the validation cohort. We first performed ST on an additional human CRC tumor sample (CRC8) and validated the ST results using mIHC staining on consecutive slices from the same tissue block utilized for ST (Fig. 3A, fig. S4, A to C, and Methods). The ST data were annotated using unpaired scRNA-seq data obtained from independent CRC datasets (Fig. 3B)^41^. Quantitative analysis of cellular proportions via mIHC revealed that among 12 validated immune subpopulations, the majority of them had an average proportion of less than 10% within the pseudo-spots (Methods), with T cell subpopulations exhibiting the lowest proportions (fig. S4D). In alignment with the results from mIHC, UCASpatial correctly annotated the spatial distribution of these cell subpopulations (Fig. 3C and fig. S4, E to H). We further compared the cellular proportions estimated by other computational algorithms with those quantified by mIHC in tumor regions. Our analysis demonstrated that UCASpatial consistently achieved the highest PCC and the lowest RMSE for these immune cell subpopulations, particularly for T cell subpopulations, in tumor regions (Fig. 3C and fig. S4, E to H). These findings underscore the capability of UCASpatial in accurately identifying low-abundant cell subpopulations, even in complex tissue contexts.

We found that the spatial distribution patterns of three CD8^+^ T cell subpopulations at the tumor-invasive front were distinct: CD8^+^ Tex and CD8^+^ Trm were more proximal to the tumor, whereas CD8^+^ Tem was relatively distal (Fig. 3, D and E). Also, there was a higher proportion of CD8^+^ Tex and CD8^+^ Trm infiltrated into the tumor region than CD8^+^ Tem (Fig. 3E). To validate these findings, we performed mIHC on human CRC samples from two additional patients (CRC9 and CRC10) in the validation cohort (Fig. 3A) and quantified the distance of each CD8^+^ T cell subpopulation to the tumor. This analysis revealed that PD-1^+^ CD8^+^ T cells and CD103^+^ CD8^+^ T cells were positioned closer to the tumor compared to GZMK^+^ CD8^+^ T cells (Fig. 3, F and G). Further calculation of the proportion of each CD8^+^ T cell subpopulation within the tumor region supported that PD-1^+^ CD8^+^ T cells and CD103^+^ CD8^+^ T cells exhibited higher tumor infiltration proportions compared to GZMK^+^ CD8^+^ T cells (Fig. 3, F and G). Additionally, we noticed UCASpatial without entropy-weighting and meta-purity filtering was unable to either identify T cell subpopulations in TLS regions or discriminate the spatial distribution of tumor-infiltrating CD8^+^ T cell subpopulations (Fig. 3D, fig. S3, I to L, and fig. S4, E to H). These results demonstrate that UCASpatial can accurately resolve cell subpopulations in the complex TIME.

### Identification of functional multicellular communities associated with T cell-rich and T cell-excluded clonalTIMEs in CRC

Tumor-intrinsic genomic alterations have collateral effects that modify the spatial organization of the TIMEs ^46-48^. To spatially characterize diverse TIMEs at a detailed clonal resolution, we generated an integrated immunogenomics platform termed ‘UCASpatial-clonalTIME’ to spatially link tumor genomics with the TIMEs. This platform enables the identification of tumor clones within tissue slices by inferring spatial copy number variations (CNVs) ^39^ and concurrently estimating the composition of the TIME for each clone (clonalTIME) (Fig. 3H, fig. S4I, and Methods). By assessing the proportions of CD4^+^ and CD8^+^ T cell subpopulations, we categorized the clonalTIMEs into two phenotypes: T cell-rich and T cell-excluded (Fig. 3I, fig. S4J, and Methods). We detected three out of thirteen tumor clones displaying the T cell-rich clonalTIME phenotype (Fig. 3I and fig. S4, J to K). Within these T cell-rich clonalTIMEs, all three subpopulations of CD8^+^ T cells—CD8^+^ Tem, CD8^+^ Trm, and CD8^+^ Tex—were notably enriched (Fig. 3I). Analysis of the CD4^+^ T cell compartment showed significant enrichments of CD4^+^ Tm and Th1/Th17 within these T cell-rich clonalTIMEs (Fig. 3I). Conversely, within T cell-excluded clonalTIMEs, we found a depletion of all T cell subpopulations. Instead, there was an enrichment of classical monocytes (cMo), neutrophils, and *TREM2*^+^ *PDPN*^+^ macrophages (Fig. 3I).

Immune cells within the TIMEs engage in crosstalk with non-immune cells that coordinately form multicellular communities. To identify these communities associated with two clonalTIME phenotypes, we calculated the PCC between the neighborhood-averaged proportions of various cell subpopulations including 22 immune subpopulations and 7 non-immune subpopulations across all spots from the 13 clonalTIMEs (Methods). Through hierarchical clustering followed by functional enrichment analysis (Methods), we observed three multicellular communities discriminated by different cellular compositions and specialized functionalities: 1) Community1 encompassed *HES1*^hi^ *C1QC*^hi^ macrophages, CD8^+^ Tem, CD8^+^ Tex and inflammatory *CD34*^+^ fibroblast (*CD34*^+^ iFib) (Fig. 3J and fig. S4M), with enriched pathways in leukocyte mediated cytotoxicity and homeostasis (fig. S4N); 2) Community2 comprised endothelial cells, pericytes, *TREM2*^+^ *PDPN*^+^ macrophages, *TREM2*^+^ *APOC1*^hi^ macrophages, *FAP*^+^ myofibroblasts, smooth muscle, and glial cells (Fig. 3J), with functional pathways related to angiogenesis, wound healing, and WNT signaling (fig. S4N); 3) Community3, including neutrophils and cMos (Fig. 3J), exhibited enriched pathways in cytokine-mediated signaling pathway and acute inflammatory response (fig. S4N).

To link these communities with T cell-rich versus T cell-excluded clonalTIME phenotype, we examined the score of each community in the respective clonalTIME phenotypes (Methods). We observed a significantly higher score of community1 within T cell-rich clonalTIMEs, while the score of community2 was higher in T cell-excluded clonalTIMEs (Fig. 3K). The score of community3 showed no significant differences between the two clonalTIME phenotypes (Fig. 3K). Moreover, an opposite enrichment trend of community1 and community2 was detected across 13 clonalTIMEs (PCC = -0.51) (Fig. 3L). We next closely examined community distributions in a particular ST slice (CRC4) containing both T cell-rich (clone 1 and 2) and T cell-excluded (clone 3) tumor clones. The result revealed a loss of community1 coincident with an increase of community2 and community3 as the clonalTIME transitioned from a T cell-rich to a T cell-excluded phenotype (Fig. 3, M and N). Together, we spatially characterized the multicellular communities within T cell-rich and T cell-excluded TIME at a clonal resolution and identified two of these functional communities associated with T cell-rich versus excluded clonalTIME phenotypes in CRC.

### Identification of tumor genetic alterations associated with clonalTIME transition in CRC

To investigate whether genetic alterations are associated with immune phenotypic transitions in tumor clones, we conducted a comparison of CNVs across clones (fig. S5A and table S5). Our findings revealed a distinct pattern among T cell-excluded clones, exhibiting unique CNVs that are absent in T cell-rich clones (Fig. 4A). Notably, we observed a frequent copy number gain on chromosomes 20q, 20p, and 7p (chr20q-gain, chr20p-gain, and chr7p-gain) within T cell-excluded clones (Fig. 4A). This observation prompted us to hypothesize that these chromosomal gains may arise during clonal evolution and promote the transition within the tumor microenvironment from a T cell-rich to a T cell-excluded state.

**Fig. 4.**
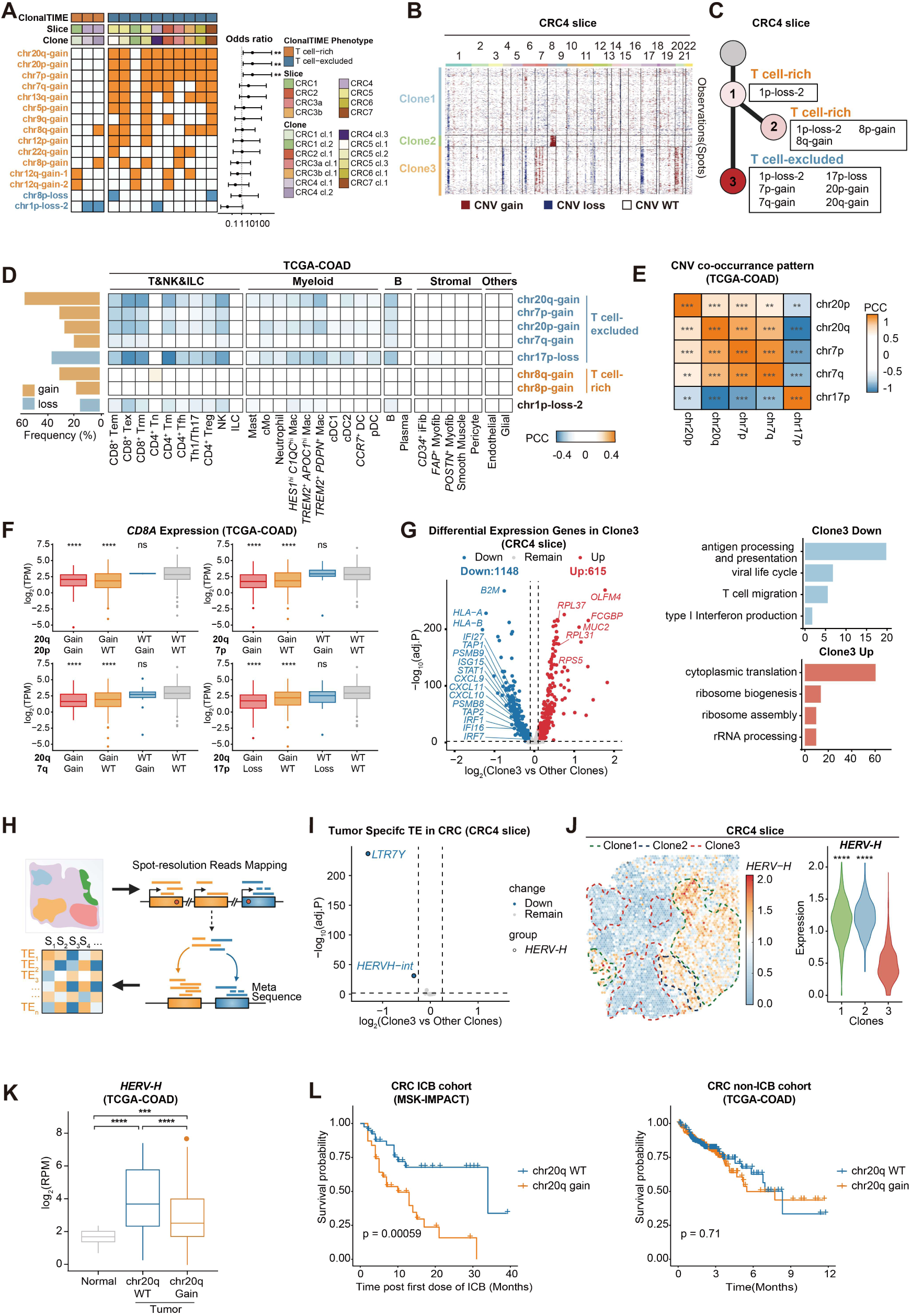
UCASpatial reveals clonalTIME transition orchestrated by chr20q-gain-associated endogenous retrovirus silencing. (**A**) Heatmap showing copy number variation (CNV) landscape of each clone grouped by clonalTIME phenotypes. Colors indicate the CNV status (Orange indicates CNV gain, and blue indicates CNV loss). Columns in the heatmap were annotated by cloneID, sliceID, and clonalTIME phenotype. Forest plots showing the odds ratio associated with clonalTIME phenotypes of each CNV. Dots indicate the calculated odds ratio, and whiskers indicate 95% confidence intervals (CIs). Only CNVs occurring in at least 2 clones were shown in the heatmap. (**B**) Heatmap showing the CNV profile inferred from spatial transcriptomics data from the CRC4 slice. The other slices are shown in Figure S4A. Clonal groupings of spots were determined by hierarchical clustering. (**C**) Phylogenetic tree of three clones in the CRC4 slice. The corresponding clonalTIME phenotype of each clone is listed alongside. The CNV status of each clone is listed (summarized by the chromosome number where the event was located, p/q arm, and gain/loss; full genomic locations of each CNV are provided in table S5). Gray dots represent benign cells, while red dots signify tumor clones. The intensity of red corresponds to the number of CNVs. (**D**) Left, bar plot showing the frequency of each CNV status in the TCGA-COAD dataset. Right, heatmap showing the PCC between the copy number of indicated CNV status and signature scores of each cell subpopulation in the TCGA-COAD dataset. Colors represent the PCC value. Blank squares indicate non-significant associations (PCC < 0.2). The color of CNV names represents associated clonalTIME phenotypes in the CRC4 slice. (**E**) Heatmap displaying the PCC between the copy numbers of different CNV statuses within the TCGA-COAD dataset. Colors represent the PCC value. (**F**) Boxplot showing *CD8A* expression in samples with indicated CNV status in the TCGA-COAD dataset. (**G**) Gene expression difference between clone3 and other clones in the CRC4 slice. Left, volcano plot displaying differentially expressed genes in clone3 versus other clones in the CRC4 slice. Differentially expressed genes were defined by adjusted p-value < 0.01 and |log_2_(fold change)| ≥ 0.1. P-values were calculated using the Wilcoxon rank-sum test and adjusted by the Bonferroni correction method. Right, functional enrichment analysis of upregulated (n=615) and downregulated genes (n = 1148) of clone3. (**H**) Schematic representation of transposable element (TE) quantification in spatial transcriptomics. The resulting assignments are condensed into a matrix of read counts for each cell, versus each TE. (**I**) Volcano plot showing the differentially expressed TEs between clone3 and other clones in the CRC4 slice. Differentially expressed TEs of clone3 were defined by adjusted p-value < 0.01 and a |log_2_(fold change)| ≥ 0.25. P-values were calculated using the Wilcoxon rank-sum test and adjusted by the Bonferroni correction method. (**J**) *HERV-H* expression of each clone in the CRC4 slice. Left, spatial visualization of *HERV-H* expression in the CRC4 slice. Dashed lines mark the general contour of each tumor clone region. Right, violin plot displaying the *HERV-H* expression across 3 clones in the CRC4 slice. (**K**) Boxplot representing the *HERV-H* expression across normal tissue, chr20q-WT, and chr20q-gain tumor samples in the TCGA-COAD dataset. (**L**) Prognostic significance of chr20q-gain in ICB-treated CRC patients. Left, overall survival (Kaplan-Meier curves) of CRC patients post-ICB in the MSK-IMPACT cohort stratified by the levels of chr20q CNV status. Right, overall survival (Kaplan-Meier curves) of CRC patients without ICB treatment in the TCGA-COAD dataset, stratified by the levels of chr20q CNV status. Log-Rank p-values are shown. Figure (**H**) is adapted from images created with BioRender.com. In Figures (**A**, **E**, **F**, **J**, and **K**), statistical analysis was performed using a two-tailed unpaired *t*-test. ns, non-significant, p > 0.05; *0.01 ≤ p < 0.05; **p < 0.01; ***p < 0.001; ****p < 0.0001. Boxplot elements in Figures (**F**, **K**, and **L**) are defined as: center line, median; box limits, upper and lower quartiles; whiskers, 1.5× interquartile range; points, outliers.

To further investigate the relationship between genetic alterations and the immunophenotype of tumor clones, we focused on the aforementioned CRC4 ST slice containing three clones exhibiting sequential clonal evolution. Clone1, characterized by chr1p-loss-2, served as a common ancestor (Fig. 4, B and C). It gave rise to two distinct progenies: clone2 that acquired chr8p-gain and chr8q-gain, and clone3 with chr20q-gain, chr20p-gain, chr7p-gain, and two additional CNVs (Fig. 4, B and C). Notably, while the evolution from clone1 to clone2 was accompanied by maintenance of a T cell-rich TIME (Fig. 4C), the evolution to clone3 was marked by a transition to the T cell-excluded phenotype, evidenced by a significant reduction in CD8^+^ Tex, CD8^+^ Tem, and *HES1*^hi^ *C1QC*^hi^ macrophages infiltration within their clonalTIMEs (Fig. 3, I and N, and Fig. 4C). This transition suggests that the genetic changes accumulated in clone3, particularly the chr20q-gain and associated CNVs, may be critical in shaping a T cell-excluded TIME.

For further validation of the relationship between the genetic changes specific to CRC4-clone3 and the T cell-exclusive immunophenotype, we conducted an integrative analysis utilizing the TCGA-COAD dataset, which encompasses DNA copy-number variation (Affymetrix 6.0 microarrays) and bulk RNA sequencing data with 471 samples. We identified the existence of CRC4-identified CNVs and estimated the infiltration levels of various immune cell subpopulations based on their signature expressions derived from bulk RNA-seq for each patient. We then calculated the PCC between the presence of these CNVs and estimated cell subpopulations (fig. S5B), revealing significant correlations between the presence of clone3-specific CNVs and the reduction in the signature expression of various T cell subpopulations (Fig. 4D).

Notably, all of these five clone3-specific CNVs (chr20q-gain, chr20p-gain, chr7p-gain, chr7q-gain, and chr17p-loss) exhibited a concurrent occurrence pattern within the TCGA-COAD dataset (Fig. 4E). To determine the primary CNV responsible for the diminished T-cell infiltration, we isolated the individual impacts of each CNV in the TCGA-COAD dataset based on their presence (Methods). Our analysis revealed that only the group harboring chr20q-gain, prevalent in 57.9% of CRC patients (Fig. 4D), exhibited a significant reduction in the expression levels of T cell marker genes (*CD3G*, *CD8A*, and *CD4*) when compared to the group without chr20q-gain (Fig. 4F and fig. S5C). Since chromosomal instability (CIN) has been previously reported to be associated with decreased immune cell infiltration ^49^, we stratified the samples by aneuploidy levels to control for its potential effects (Methods). Patients with chr20q-gain showed significantly lower T cell marker expression than chr20q-WT patients in both low and intermediate aneuploidy groups (fig. S5D), demonstrating a CIN-independent role of chr20q-gain in driving T cell exclusion. These findings suggest that the presence of chr20q-gain may play a pivotal role in driving the transition of the clonalTIME phenotype from T cell-rich to T cell-excluded.

### Tumor-intrinsic endogenous retrovirus silencing correlates with chr20q-gain related T cell-excluded clonalTIME formation

To investigate the potential mechanism of the immune phenotypic transitions in tumor clones, we performed differential gene expression analysis in clone3, which had acquired a chr20q gain, compared to clone1 and clone2 of CRC4. Through functional enrichment analysis, we identified enrichment for pathways related to translation machinery among the upregulated genes in clone3 (Fig. 4G). Conversely, pathways related to type I interferon (IFN) production were notably enriched among the downregulated genes, indicating a potential suppression of immune response (Fig. 4G and fig. S5, E to G). Notably, chemokine-coding genes responsible for cytotoxic CD8^+^ T cell recruitment, such as *CXCL9* and *CXCL10*, and genes associated with antigen presentation, including *TAP1*, *TAP2*, *HLA-A*, *HLA-B*, and *B2M* ^50, 51^, exhibited reduced expression (Fig. 4G and fig. S5, E to G). This observation potentially elucidates the T cell-excluded phenotype of clone3.

It is well-established that the activation of the type I IFN response in tumors can be attributed to the innate sensing of cytosolic RNA derived from transposable elements (TEs)^52^. Considering their role as internal adjuvants eliciting T cell responses against tumors, we hypothesized that differences in TE expression levels could directly influence the immune phenotypic transition from T cell-rich to T cell-excluded. To examine this hypothesis, we conducted a spatial analysis of the expression of 40 TEs that specifically over-expressed in CRC tumor cells (Fig. 4H and Methods) ^36, 53^. Our analysis identified two CRC-specific TEs—the internal sequence of human endogenous retrovirus subfamily H (*HERV-H*) and its associated long terminal repeat *LTR7Y* ^54, 55^—were significantly downregulated in clone3 compared to clone1 and clone2 (Fig. 4, I and J, fig. S5H, and Methods). The statistical analysis across all slices verified the observations in CRC4 (fig. S5I). This downregulation suggests that tumor cells might suppress *HERV-H* expression to evade immune recognition in CRC.

To investigate whether the decreased *HERV-H* expression and attenuated antiviral response observed in clone3 were associated with chr20q-gain, we analyzed TCGA-COAD tumor samples. Notably, tumors with chr20q-gain showed significantly reduced *HERV-H* expression and diminished viral transcription and type I IFN signaling compared to chr20q-WT (Fig. 4K, fig. S5, J to K, and Methods). We also confirmed the attenuated type I IFN response in tumor cells with chr20q-gain using scRNA-seq data from our cohort (fig. S5, L to O). These results collectively suggest a compelling correlation between the chr20q-gain in tumor cells and the downregulation of *HERV-H*, which may be implicated in the immune evasion strategies of CRC.

Given our discovery of a strong correlation between chr20q-gain and the consequent downregulation of *HERV-H*, which results in a T cell-excluded immune phenotype, coupled with the recognized influence of T cells on the clinical response to immune checkpoint blockade (ICB) treatment ^56^, we hypothesized that chr20q-gain might influence the response to ICB treatment in CRC patients. To examine this, we evaluated survival post-ICB treatment by categorizing patients from Memorial Sloan Kettering Cancer Center (MSKCC) IMPACT (MSK-IMPACT) cohorts (comprising 71 patients) ^57, 58^ into two groups based on their chr20q status. Our evaluation revealed that patients with chr20q-gain exhibited significantly poorer survival outcomes following ICB treatment (Fig. 4L and fig. S5P). In contrast, this adverse survival trend was not observed in CRC patients who did not receive ICB treatment (TCGA-COAD dataset) (Fig. 4L and fig. S5Q). To rule out the potential impact of tumor mutation burden (TMB), we categorized patients based on both CNV status and TMB. Our analysis indicated that the correlation between chr20q-gain and diminished survival in the CRC ICB cohort remained irrespective of TMB levels (fig. S5R). Taken together, we identified chr20q-gain, a prevalent genomic defect in CRC, as a genomic biomarker for T-cell exclusion, suggesting its utility as an immunogenomic parameter for predicting responsiveness to ICB. Furthermore, our findings lay the groundwork for a novel model in which tumor cells with chr20q-gain orchestrate a T cell-excluded clonalTIME by suppressing *HERV-H* expression during clonal evolution, thereby potentially evading immune surveillance.

### UCASpatial accurately annotates cell subpopulations with spatiotemporal dynamics in the murine wound healing ST dataset

To demonstrate UCASpatial’s utility, we next applied it to another real ST dataset, focusing on murine wound healing - a complex, highly organized multicellular process with distinct spatiotemporal dynamics ^59-61^. We employed a 2 mm ear punch wound model in MRL/MpJ mice (hereafter denoted as MRL), a strain with super-healing capacity, which fully regenerates the ear punch wound without fibrosis ^62-64^. As a control, we used non-regenerating strain C57BL/6 mice (hereafter denoted as B6), known to form scar tissue over ear wound edges (Fig. 5A) ^62-64^. To interrogate the tissue-wide spatiotemporal dynamics of various cell subpopulations during wound healing, we collected ear tissue samples from MRL and B6 mice at the following stages: unwounded (day 0), early inflammatory (day 3), return-to-unwounded (day 7), and regeneration stage (day 15) ^60^. We performed ST sequencing using the 10x Genomics Visium platform (Fig. 5, B and C). As a representative reference for capturing the cellular diversity present throughout the wound healing process, we utilized scRNA-seq data from day 7 ear tissues from both MRL and B6 mice, classifying them into 23 distinct cell subpopulations (Fig. 5D and fig. S6, A and B).

**Fig. 5.**
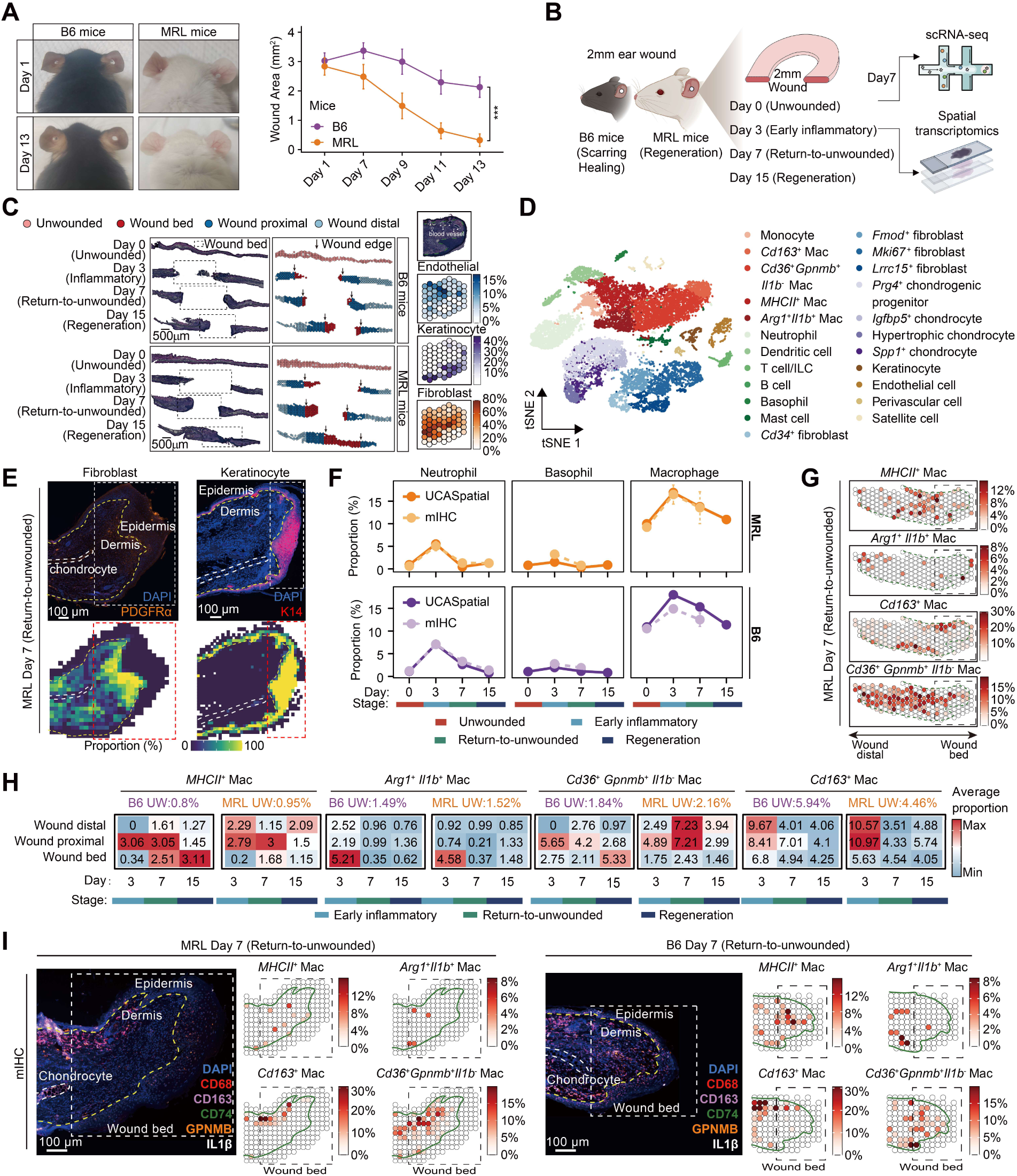
UCASpatial reconstructs the spatiotemporal dynamics of diverse macrophage subpopulations during murine wound healing. (**A**) Left, Gross photographs at day 1 and day 13 post-wounding of B6 and MRL mice. Right, wound curves for B6 (n = 4) and MRL (n = 4) ear wounds show closure over time. Wound area was recorded and calculated as π×*r*1×*r*2, *r*1 and *r*2 represent radius aligned with or orthogonal to the body axis, respectively. Statistical analysis was performed using a two-tailed unpaired *t*-test. ***p < 0.001. (**B**) Graphical overview of experimental design. scRNA-seq was performed using total cells from MRL mice ear tissue on day 7 post-wounding. Spatial transcriptomics data were generated from MRL mice ear tissue on day 0 (unwounded), day 3, day 7, and day 15 post-wounding. Figure adapted from images created with BioRender.com. (**C**) Left, hematoxylin and eosin (H&E) staining of MRL and B6 ear tissues and spatial plots of the mouse ear displaying four groups of regions annotated based on H&E staining: unwounded (day 0), wound bed (1 mm radius), wound proximal (0-1 mm from the wound bed), and wound distal (> 1 mm from the wound bed). The rectangles represent the wound bed. Scale bar, 500 μm. Right, micro-anatomic annotation of wound bed on day 7 in MRL mice and spatial visualization of endothelial cells, keratinocytes, and fibroblasts at the wound bed on day 7. The green dashed line represents the edge of the epidermis-dermis, and the white dashed line represents the blood vessels. Colors represent the proportion of these three cell types. The proportion of fibroblasts was calculated by the sum of the 4 fibroblast subpopulations’ proportions. (**D**) The t-distributed Stochastic Neighbor Embedding (tSNE) plot showing 23 cell subpopulations of all cells from scRNA-seq. (**E**) Upper, representative immunofluorescence imaging of fibroblasts (left) and keratinocytes (right) using ear tissues in MRL and B6 mice at the return-to-unwounded stage (day 7 post-injury). The staining images indicate DAPI (blue), PDGFRα (orange), and K14 (red). Scale bar, 100 μm. Bottom, a quantitative evaluation of the distributions of fibroblasts (left) and keratinocytes (right) based on the mIHC, with a 50 μm-bin resolution. Cells were segmented based on DAPI using Qupath software. The dashed rectangles indicate the wound bed. The dashed yellow lines indicate the interface of the epidermis-dermis. The dashed white lines indicate chondrocytes. (**F**) Line plots showing the temporal dynamics of the proportion of neutrophil, basophil, and macrophage between MRL and B6 mice in both UCASpatial (solid lines) and mIHC (dashed lines). (**G**) Spatial visualization of the distribution of indicated macrophage subpopulations at the wound bed on day 7 based on UCASpatial. Colors represent the proportion of each macrophage subpopulation. The dashed rectangles represent the wound bed. The green dashed line indicates the dermal-epidermal junction. (H) Heatmap showing the spatiotemporal dynamics of macrophage subpopulations of three anatomic regions. Colors and the numbers in the heatmap represent the average proportion of macrophage subpopulations at specific regions and stages. The number of UW (above the heatmap) represents the average proportion of each macrophage subpopulation on Day 0, respectively. UW, unwounded. (**I**) Representative immunofluorescence imaging and the quantitative evaluation (50μm-bin resolution) of the distributions of indicated macrophage subpopulations at the return-to-unwounded stage (day 7 post-injury) in MRL mice (left) and B6 mice (right). For the mIHC, the dashed yellow line indicates the interface of the epidermis-dermis; the dashed white line indicates the chondrocyte; the dashed rectangle indicates the wound bed. The staining images indicate DAPI (blue), CD68 (red), CD163 (purple), CD74 (green), GPNMB (orange), and IL1β (white). Scale bar, 100 μm. For the quantitative evaluations, the green line indicates the interface of the epidermis-dermis; the dashed black rectangle indicates the wound bed. Colors represent the proportion of each macrophage subpopulation.

UCASpatial successfully reconstructed the anatomical distributions of structural cell types, including keratinocytes, endothelial cells, and fibroblasts (Fig. 5, C and E, and fig. S6C). Moreover, UCASpatial resolved the spatiotemporal dynamics of wound-associated immune cells. Specifically, an increase in myeloid cells–neutrophils, basophils, and macrophages–was observed at the early inflammatory stage in both MRL and B6 mice (Fig. 5F). This observation is consistent with the established notion that innate immune cells are the first responders to injury ^65-67^. At the return-to-unwounded stage, neutrophils and basophils exhibited a reduction in frequency, whereas macrophages maintained a high abundance (Fig. 5F), underscoring their roles in both the early injury response and the subsequent tissue repair and regeneration processes ^60, 65, 66, 68, 69^.

We experimentally validated the spatial distribution and proportion of six major cell types including keratinocytes, fibroblasts, endothelial cells, neutrophils, macrophages, and basophils, using multiplexed immunohistochemistry (mIHC) (fig. S6D). The anatomical distributions of major cell types detected by mIHC were consistent with UCASpatial’s deconvolution analysis (Fig. 5, C and E, and fig. S6, E to G). Notably, the spatiotemporal dynamics of cell subpopulations quantified by UCASpatial were more closely aligned with the mIHC results than those obtained by other methods (Fig. 5F and fig. S6G). This consistency between the mIHC and UCASpatial’s deconvolution results validates the accuracy of UCASpatial.

To compare the performance of UCASpatial with existing methods, we calculated the PCC between the spatial distribution of cell-subpopulation proportions and marker-gene scores (table S6 and Methods), which allowed us to quantify the performance of deconvolution in the absence of ground truth ^16^. Among all methods compared, UCASpatial exhibited the highest PCC across cell subpopulations (fig. S6, H and I). Specifically, by comparing UCASpatial with “UCASpatial ablation”, which shares the same framework but lacks the core functions of entropy-based weighting and meta-purity filtering process, we noticed that the improved accuracy of UCASpatial is attributed to the incorporation of entropy weighting and meta-purity filtering (fig. S6, H and I). Notably, existing methods lacked precision in identifying immune cell subpopulations during wound healing. For example, RCTD, stereoscope, and Spotiphy failed to accurately capture the spatiotemporal dynamics of neutrophils, which peaked at the early inflammatory stage in both strains (fig. S6G). Additionally, RCTD, cell2location, CARD, and Spotiphy showed limited ability to detect macrophages at the unwounded stage (fig. S6G).

Further, UCASpatial precisely captured the spatial segregation patterns of macrophage subpopulations. Specifically, *MHCII^+^* macrophages, present at all measured time points, were predominantly located in the wound’s proximal and distal regions in MRL mice, while predominantly located in the wound’s bed and distal regions in B6 mice (Fig. 5, G and H, and fig. S6, J to L). *Arg1^+^ Il1b^+^* inflammatory macrophages, which appeared transiently in the wound bed at the early inflammatory stage (Fig. 5I), preferentially clustered at the migrating epithelial tongue (Fig. 5G and fig. S6M). In contrast to *Arg1^+^ Il1b^+^* macrophages, *Cd163*^+^ macrophages with enriched functions in endocytosis and lysosome (fig. S6N) predominantly resided along the epidermis layer outside the migrating epithelial tongue across all measured time points post-injury (Fig. 5G and fig. S6M). *Cd36^+^ Gpnmb^+^ Il1b^-^* macrophages, functionally resembling lipid-associated macrophages implicated in fibrosis ^70-72^ (fig. S6N), are located in the wound proximal and distal regions at the return-to-unwounded stage (Fig. 5, G and H). Such spatial distribution patterns of four macrophage subpopulations at the return-to-unwounded stage were confirmed by mIHC (Fig. 5I).

We also observed a differential distribution pattern of *Cd36^+^ Gpnmb^+^ Il1b^-^* macrophages in B6 and MRL mice. This subpopulation was initially located in the wound proximal and distal regions of both strains at the return-to-unwounded stage (Fig. 5, G and H, and fig. S6L). However, at the regeneration stage, *Cd36^+^ Gpnmb^+^ Il1b^-^* macrophages in B6 mice accumulated within the wound bed, while in MRL mice, they remained distal to the wound bed (Fig. 5H and fig. S6O). This unique migration pattern observed in B6 mice might explain their scarring-healing phenotype. Collectively, UCASpatial reconstructs a fine-grained spatiotemporal cellular landscape in murine wound healing, revealing the unique spatial segregation and temporal dynamics of macrophage subpopulations that serve in different stages of wound healing.

### Spatial characterization of molecularly distinct chondrocyte subpopulations linked to regenerative versus scarring-healing phenotypes

After demonstrating our capability to spatially map various cell subpopulations in wounded tissues, we proceeded to investigate the cellular events orchestrating regenerative versus scarring-healing phenotypes in MRL and B6 mice between 7 to 15 days post-injury. We identified a scarring-associated myofibroblast subpopulation *Lrrc15^+^* fibroblasts located within the wound bed at the return-to-unwounded stage in both MRL and B6 mice^73-75^ (Fig. S7, A and B). While a reduction in the frequency of *Lrrc15^+^*fibroblasts was observed in both strains at the regeneration stage, these cells were significantly more enriched in B6 mice compared to MRL (Fig. S7, A and B). This differential enrichment may account for the scarring-healing phenotype observed in B6 mice.

We also delineated the spatiotemporal dynamics of keratinocytes and chondrocytes, two major cell populations contributing to regenerative healing ^76, 77^. We observed a comparable distribution of keratinocytes in MRL and B6 mice within regions of the migrating epithelial tongue at the return-to-unwounded stage (Fig. S7, C and D, and fig. S7E), indicating complete re-epithelialization occurred in both strains. However, at the regeneration stage, while the proportion of keratinocytes remained high within the wound bed of MRL mice, a substantial decrease was observed in B6 mice, supporting the notion that keratinocytes play a crucial role in the regeneration process (Fig. S7, C and D) ^78-80^.

Spatial mapping of four chondrocyte subpopulations (fig. S7, F and G) during post-injury chondrogenesis revealed distinct regional and temporal patterns. Injury-induced, de-differentiated *Spp1*^+^ chondrocytes ^81^ were acutely present at the wounded bed, with comparable levels in MRL and B6 mice at the early inflammatory stage (Fig. 6A). *Prg4*^+^ chondrogenic progenitors, which give rise to hypertrophic chondrocytes ^82-84^, were transiently observed within the wound bed of B6 mice but persisted in MRL mice from the return-to-unwounded stage till the regeneration stage (Fig. 6, A and B). Consistently, a higher proportion of mature hypertrophic chondrocytes was observed in MRL mice at the regeneration stage (Fig. 6A), suggesting that this progenitor is linked to regenerative healing in MRL mice.

**Fig. 6.**
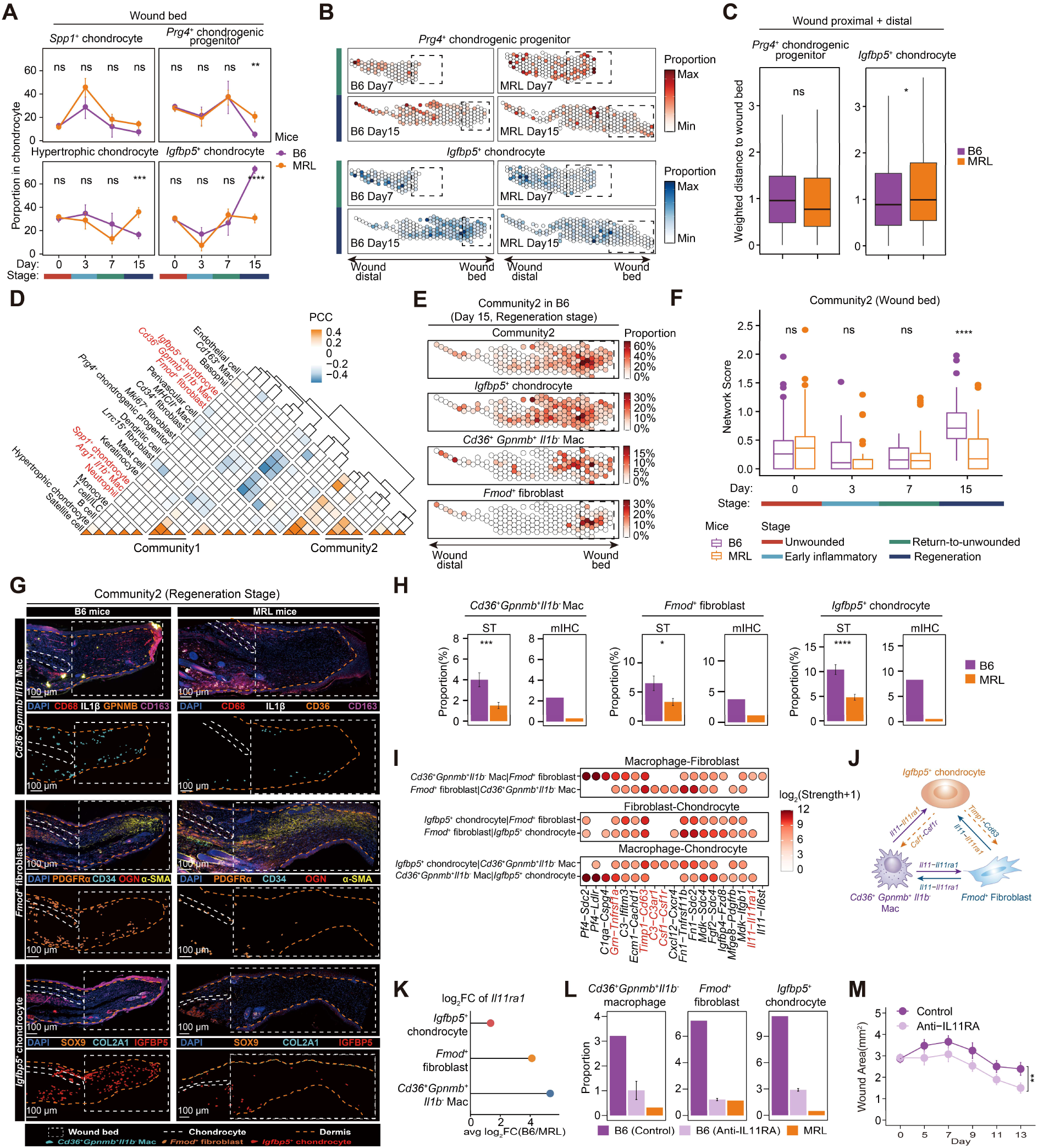
Identification of molecularly distinct chondrocyte subpopulations associated with regenerative versus scarring-healing phenotypes. (**A**) Line plots showing the temporal dynamics of the proportion of four chondrocyte subpopulations at the wound bed between MRL and B6 mice. (**B**) Spatial visualization of the distribution of indicated *Igfbp5*^+^ chondrocyte and *Prg4*^+^ chondrogenic progenitor on day 7 and day 15 in MRL and B6 mice. Colors represent the proportion of each chondrocyte subpopulation. Dashed boxes represent the wound edge. (**C**) Boxplots showing the weighted distance of chondrocyte cell subpopulations in the wound proximal and distal regions to the wound bed on day 15 (regeneration stage) in MRL and B6 mice. Boxplot elements are defined as: center line, median; box limits, upper and lower quartiles; whiskers, 1.5× interquartile range. Statistical analysis was performed using a one-tailed unpaired t-test. *0.01 ≤ p < 0.05. (**D**) Heatmap showing the PCC between the proportion of each two cell subpopulations across all spots after neighborhood processing within the left wound bed (Methods). Communities were identified using hierarchical clustering. (**E**) Spatial visualization of the distribution of indicated community2, *Igfbp5*^+^ chondrocyte, *Fmod*^+^ fibroblast, and *Cd36^+^ Gpnmb^+^ Il1b^-^* macrophage subpopulations at the wound bed on day 15 in B6 mice based on UCASpatial. Colors represent the proportion of each cell subpopulation. The dashed rectangles represent the wound bed. (**F**) Boxplots showing the score of community2 at the wound bed in MRL and B6 mice at different stages. Boxplot elements are defined as: center line, median; box limits, upper and lower quartiles; whiskers, 1.5× interquartile range; points, outliers. (**G**) Representative immunofluorescence imaging of cell subpopulations within community2, including *Cd36^+^ Gpnmb^+^ Il1b^-^* macrophage (upper), *Fmod*^+^ fibroblast (middle), and *Igfbp5*^+^ chondrocyte (bottom), using ear tissues in B6 (left) and MRL mice (right) at the regeneration stage (day 14 post-injury). The dashed white lines indicate chondrocytes. The dashed white rectangle indicates the wound bed. The dashed yellow line indicates the interface of the epidermis-dermis. Scale bar, 100 μm. (**H**) Bar plots showing the proportion of *Cd36^+^ Gpnmb^+^ Il1b^-^* macrophages (left), *Fmod*^+^ fibroblasts (middle), and *Igfbp5*^+^ chondrocytes (right) in wound bed regions at the regeneration stage in ST data and mIHC. (I) Bubble heatmaps showing the log_2_ strength for selected ligand-receptor pairs between *Igfbp5*^+^ chondrocyte, *Fmod*^+^ fibroblast, and *Cd36^+^ Gpnmb^+^ Il1b^-^* macrophage from the scRNA-seq of B6 mice. Attraction strength was calculated by CellPhoneDB. Colors show the attraction strength. (**J**) Schematic diagram showing the summary of indicated ligand-receptor pairs between *Igfbp5*^+^ chondrocytes, *Fmod*^+^ fibroblasts, and *Cd36*^+^ *Gpnmb*^+^ *Il1b*^-^ macrophages. (**K**) Lollipop plot showing the log_2_ fold change of *Il11ra1* in *Igfbp5*^+^ chondrocytes, *Fmod*^+^ fibroblasts, and *Cd36*^+^ *Gpnmb*^+^ *Il1b*^-^ macrophages across MRL and B6 mice. (**L**) Bar plots showing the proportion of *Cd36^+^ Gpnmb^+^ Il1b^-^* macrophages (left), *Fmod*^+^ fibroblasts (middle), and *Igfbp5*^+^ chondrocytes (right) in wound bed regions at the regeneration stage in mIHC data. The B6 control group was administered with anti-rabbit IgG. B6 control, n = 1; B6 Anti-IL11RA, n = 2; MRL, n = 1. (**M**) Wound curves of B6 ear wounds in the control (n = 5) and rosiglitazone-treated (n = 7) groups show wound healing over time. The control group was administered with anti-rabbit IgG. Wound area was recorded and calculated as π×*r*1×*r*2, *r*1 and *r*2 represent radius aligned with or orthogonal to the body axis, respectively. Statistical analysis in Figures (**A**, **C**, **F**, **H,** and **M**) was performed using a two-tailed unpaired *t*-test. ns, non-significant, p > 0.05; *0.01 ≤ p < 0.05; **p < 0.01; ***p < 0.001; ****p < 0.0001.

In addition to the *Prg4*^+^ chondrogenic progenitor, we noticed an uncharacterized *Igfbp5*^+^ chondrocyte subpopulation that expressed *Cd34* and *Ogn* (fig. S7G). *Igfbp5*^+^ chondrocyte was closely related to *Prg4*^+^ chondrogenic progenitor and was maintained in a less-differentiated state, as evidenced by lower expression levels of *Col2a1* and *Acan* (fig. S7, H to K) ^85, 86^. Functional analysis of *Igfbp5*^+^ chondrocytes showed enriched pathways for extracellular matrix (ECM) organization and ECM-receptor interaction (Fig. S7L), implying potential interaction between *Igfbp5*^+^ chondrocytes and ECM-producing cells like fibroblasts and macrophages ^87^. Intriguingly, the spatiotemporal dynamics of *Igfbp5*^+^ chondrocytes differed in MRL and B6 mice. While these cells initially occupied regions proximal to the wound edge in both strains at the return-to-unwounded stage, they continued spreading into the wound bed, replacing the *Prg4*^+^ chondrogenic progenitor at the regeneration stage in B6 mice (Fig. 6, A and B). In contrast, *Igfbp5*^+^ chondrocytes were spatially restricted in MRL mice, remaining distal to the wound bed at the regeneration stage (Fig. 6, B and C). Therefore, the replacement of *Prg4*^+^ chondrogenic progenitors with *Igfbp5*^+^ chondrocytes is associated with the scarring healing phenotype in B6 mice.

### *Igfbp5*^+^ chondrocyte, *Cd36^+^ Gpnmb^+^ Il1b^-^* macrophage and *Fmod*^+^ fibroblast formed a three-cell-type cluster underlying scarring wound healing

Next, we asked whether *Igfbp5*^+^ chondrocytes positioned within the wound bed could locally crosstalk with surrounding cells and form a niche that favors scarring healing in B6 mice. To test this, we calculated the PCC between 23 cell subpopulations and identified two chondrocyte-related communities by hierarchical clustering (Methods). The community1, comprising *Spp1*^+^ chondrocytes, neutrophils, and *Arg1^+^ Il1b^+^* macrophages, formed a transient “inflammatory community” within the wound bed at the early inflammatory stage in both MRL and B6 mice (Fig. 6D and fig. S7M). The community2, which we termed the “pro-fibrotic community”, featured a stable three-cell-type cluster involving *Igfbp5*^+^ chondrocytes, ECM organization-related *Fmod*^+^ fibroblasts (fig. S7N), and *Cd36^+^ Gpnmb^+^ Il1b^-^* macrophages (Fig. 6, D to F, and fig. S7J). This cluster was particularly prominent in the wound bed at the regeneration stage in B6 mice, which has been validated by mIHC (Fig. 6, G and H, and fig. S7O).

Analysis of functional-enriched pathways across three cell subpopulations within community2 suggested their overlapping functions were responsible for fibrosis, including extracellular matrix organization, collagen fibril organization, and supramolecular fiber organization (fig. S7P and Methods). To determine whether the scarring-healing phenotype observed in B6 mice could be attributed to the formation of this pro-fibrotic community, we conducted cellular-level intervention of the pro-fibrotic community by blocking *Fmod*^+^ fibroblast differentiation. Trajectory analysis using the diffusion map algorithm revealed that *Fmod*^+^ fibroblasts might originate from *Cd34*^+^ fibroblasts (fig. S7Q). During this transition, PPARγ signaling-pathway-related genes were down-regulated, which are known to be associated with fibrosis and regeneration failure (fig. S7R) ^88-90^. As expected, treating B6 mice with the PPARγ agonist rosiglitazone inhibited the differentiation of *Cd34*^+^ fibroblasts into *Fmod*^+^ fibroblasts (fig. S7, S and T, Methods). We observed that the three-cell-type cluster in community2 was prominent in the wound bed of control B6 mice at the regeneration stage, whereas this cluster was absent in rosiglitazone-treated B6 mice (fig. S7, U and V). Additionally, rosiglitazone treatment led to a significant reduction in the remaining ear hole area, indicating enhanced wound healing (fig. S7W). Together, these findings underscore the spatial coordination among *Igfbp5*^+^ chondrocyte, *Cd36^+^ Gpnmb^+^ Il1b^-^* macrophage, and *Fmod*^+^ fibroblast, forming a three-cell-type cluster that contributes to the scarring-healing phenotype observed in B6 mice.

### Disruption of the IL11-IL11RA axis within the pro-fibrotic community enhances wound healing in B6 mice

Next, we examined the potential cross-talks within the pro-fibrotic community. Analysis of specific ligand-receptor interactions revealed a stable interaction between *Fmod*^+^ fibroblasts and *Cd36^+^ Gpnmb^+^ Il1b^-^* macrophages (Fig. 6, I and J), supporting their known partnership ^91^. Additionally, we observed reciprocal interactions between *Igfbp5*^+^ chondrocytes and both *Fmod*^+^ fibroblasts and *Cd36^+^ Gpnmb^+^ Il1b^-^* macrophages. Specifically, *Igfbp5*^+^ chondrocytes can crosstalk with *Fmod*^+^ fibroblasts via the ligand-receptor pair *Timp1-Cd63* (Fig. 6I), the ligand-receptor pair for the induction of collagen synthesis during fibrosis ^92, 93^. Further, *Igfbp5*^+^ chondrocytes can potentially interplay with *Cd36^+^ Gpnmb^+^ Il1b^-^* macrophages through the ligand-receptor pair of *C4*-*C3ar1* and *Csf1-Csf1r* (Fig. 6I), signals required for macrophage survival and polarization ^94, 95^. *Cd36^+^ Gpnmb^+^ Il1b^-^* macrophages and *Fmod*^+^ fibroblasts exhibited interaction with *Igfbp5*^+^ chondrocytes via *Il11-Il11ra1*, the ligand-receptor pair that could promote fibrosis (Fig. 6I) ^96^. Besides, *Cd36^+^ Gpnmb^+^ Il1b^-^* macrophages might also promote the proliferation of *Igfbp5*^+^ chondrocytes and *Fmod*^+^ fibroblasts via the interaction of *Grn-Tnfrsf1a* (Fig. 6I) ^97^.

Among these ligand-receptor interactions, we identified an unexplored axis involving the *Il11-Il11ra1* signaling pathway in the pro-fibrotic community (Fig. 6I). IL-11 is a member of the IL-6 cytokine family and plays a pivotal role in driving organ fibrosis and dysfunction ^98-100^. Given that IL11RA is expressed in a wide array of cell types, including stromal cells, pericytes, and epithelial cells, IL-11 signaling can trigger diverse cellular responses, leading to varied outcomes depending on the affected cell type ^98^. It remains unknown which specific cell types receive IL-11 signals during wound healing and whether this subsequently affects wound healing.

We noticed that *Il11ra1* expression was dramatically up-regulated in *Cd36^+^ Gpnmb^+^ Il1b^-^*macrophages, *Igfbp5*^+^ chondrocytes, and *Fmod^+^* fibroblasts of B6 mice compared to MRL mice (Fig. 6K and fig. S7X). This strain-specific difference suggests that the IL11-IL11RA axis might be responsible for the formation of the pro-fibrotic community in B6 mice. To test this, we blocked IL11-IL11RA signaling using an anti-IL11RA neutralizing antibody (anti-IL11RA Ab) in B6 mice. We found that the formation of the pro-fibrotic community was prevented in the wound bed in mice receiving anti-IL11RA Ab compared to the control group, indicating that IL11-IL11RA-mediated interactions contribute to the formation of the pro-fibrotic community in B6 mice (Fig. 6L and fig. S7V). Furthermore, neutralization of IL11RA significantly reduced the remaining ear hole area, suggesting that IL-11 signaling limits regeneration in B6 mice (Fig. 6M). Together, these results reveal that the strain-specific difference in wound healing between B6 and MRL mice is driven by the IL11-IL11RA1 axis, underscoring the power of UCASpatial in dissecting the molecular mechanisms underlying complex phenotypes.

## Discussion

Spatial deconvolution methodologies have made significant strides in deciphering the cell composition of well-structured tissues like the mouse brain and heart ^9-13, 17^. However, these methodologies encounter challenges when applied to complex microenvironments, where cell subpopulations typically exhibit subtle differences, low abundance, and high heterogeneity. To address these challenges, we introduce UCASpatial, a robust, sensitive, and versatile algorithm designed to deconvolute fine-grained spatiotemporal cellular composition.

UCASpatial shares conceptual similarities with existing methodologies such as SPOTlight and CARD, which apply NMF-based models to estimate cell-type compositions ^9, 12^. Yet, UCASpatial distinguishes itself through its meta-purity filtering and entropy-based weighting mechanisms, providing distinct advantages for discerning cell subpopulations with subtle transcriptomic differences, as evidenced by our ablation tests, in-silico benchmarking results, and analysis of real ST datasets. Moreover, UCASpatial does not incorporate neighboring or image information, a strategy commonly employed previously to enhance performance ^12, 14, 30, 31^. Integrating such information might not be suitable for the annotation of infiltrated immune cell subpopulations with low proportions in complex microenvironments, given their discrete distribution patterns due to recruitment dynamics. Furthermore, the integration of the UCASpatial algorithm and its downstream functions into the Seurat framework significantly facilitates a wide range of applications, streamlining the process for researchers.

Beyond UCASpatial’s strengths in resolving heterogeneous immune microenvironments and rare cell subsets, other deconvolution tools retain niche advantages that justify their continued use. Spotiphy uniquely fuses histological images with expression to deliver single-cell-resolution whole-transcriptome images. RCTD and cell2location perform comparably in benchmarks, yet RCTD’s lightweight Bayesian model demands few dependencies and minimal memory, making it attractive for rapid, resource-limited projects. Conversely, cell2location requires GPUs and large memory, but its seamless integration with Scanpy offers Python-native users a familiar, reproducible pipeline. CARD, by embedding neighborhood or morphological priors, is preferable for highly ordered tissues such as cortex or cardiac muscle, where spatial continuity dominates. Thus, while UCASpatial maximizes performance in heterogeneous, immune-rich microenvironments, method selection should balance compute budget, programming ecosystem, and the value of image information to match each biological question.

Despite its strengths, UCASpatial has some limitations that should be considered. The performance of UCASpatial is partially dependent on the quality of the reference scRNA-seq data provided by the user. While UCASpatial’s ability to leverage unpaired scRNA-seq data enhances its accessibility and utility, high-fidelity scRNA-seq data are crucial to ensure the accuracy and reliability of the deconvolution analysis. Additionally, we noticed that UCASpatial may not always outperform other methods for certain cell types, such as myofibroblasts and epithelial cells. This variability is likely due to the distinct features of these cell types, which allow most methods to annotate them with comparable accuracy. Despite these challenges, by employing entropy-based weighting in UCASpatial, we have noticed significant improvements in UCASpatial’s ability to resolve immune subpopulations compared to other methods, particularly in real datasets.

Our study employs UCASpatial to decipher the fine-grained landscape of TIMEs, offering an immunogenetic platform for linking tumor genomics with the complexity of immune landscapes at clonal resolution. In our analysis of human CRC, we identified a prevalent CNV, chr20q-gain, which emerges during clonal evolution and correlates with a T cell-excluded immune phenotype, consistent with prior observations linking chr20q-gain to low immune infiltration in tumors ^101^. By linking clonal-level cellular community transitions to genetic alterations in chr20q, we uncovered a potential molecular mechanism for T cell exclusion: the suppression of the endogenous retrovirus *HERV-H*, associated with impaired type I interferon response, during clonal evolution from chr20q-WT to chr20q-gain. Furthermore, we found that chr20q-gain serves as a novel biomarker for predicting poor outcomes in ICB therapy, a finding not previously reported. Although our analysis does not provide direct experimental evidence of causation, the consistency of these findings across multiple datasets suggests a biologically significant association. UCASpatial further extends its utility in revealing the spatiotemporal dynamics during the wound healing process, which has empowered us to dissect the previously obscure dynamics of cellular turnover and spatial organization within the regenerative process. This advancement has not only illuminated the complex interplay of cellular communities in murine wound healing, but has also provided a roadmap for future investigations into the spatial orchestration of tissue repair and regeneration. These discoveries highlight the potential of UCASpatial as a tool for uncovering clinically relevant biomarkers and deepening our understanding of the immunogenomic landscape.

As we envision the future development of UCASpatial, its integration with additional omics dimensions, such as whole-exome sequencing and spatial metabolomics, presents a promising avenue. This multi-omics approach could provide a more comprehensive elucidation of the spatiotemporal dynamics across a spectrum of biological processes. Expanding UCASpatial’s application to other cancer types may uncover novel mechanisms influencing TIMEs and offer predictive insights into the efficacy of specific immunotherapies. Furthermore, the effectiveness of UCASpatial is upon the quality of the referenced scRNA-seq data. To this end, the utilization and establishment of a robust scRNA-seq database, inclusive of a wide array of common tissues and tumor types, will be paramount. This resource would not only enhance the algorithm’s accuracy but also broaden its applicability across diverse research and clinical settings.

In conclusion, we anticipate that UCASpatial will find myriad applications in deciphering complex immune microenvironments across various developmental stages, states of health, and disease conditions within a spatial framework.

### Methods UCASpatial model

In brief, UCASpatial is an entropy-weighted model that annotates the cell-identity (equal to cell-type) compositions on each spatial location by deconvoluting the spatial expression count matrix into a predefined set of reference cell types. The model requires both spatial transcriptomics data and paired or unpaired scRNA-seq data from a given tissue as input. By default, the model takes an untransformed spatial-expression count matrix **P** ∊ℝ*^S^*^×^*^G^* of genes *g* = {1, . ., *G*} at spots *s* = {1, . ., *S*} as input, as, for example, obtained from the 10x Space Ranger software (10x Visium data). The scRNA-seq data **B**∊ℝ*^G^*^×^*^N^* serve as a reference and consist of *N* cells with the same set of *G* genes with **P**. All the *N* cells will be classified into *K* cell types which need to be provided by the users.

We denote **X**∊ℝ*^S^*^×^*^K^* as the cell-type composition matrix, where each row ***x_s_*** represents the proportions of the *K* cell types on spot *s*. By default, this matrix is estimated using an entropy-weighted non-negative least square model, which allows for accurate and robust annotation in complex situations. (see below).

### Meta-purity filtering

The existence of intermediate cells blurs cell-type borders due to their mixed identities from multiple cell types, which will disturb the recognition of cell-identity-specific gene expression profiles (cGEPs). To remove the effect brought by intermediate cells in the reference scRNA-seq data, we designed the meta-purity filtering strategy as an optional preprocessing.

Based on the input scRNA-seq reference matrix **B**, we construct meta cells using the function ‘FindClusters’ from R package ‘Seurat’ with a default resolution of 100. We recommend the resolution parameter by an expectation of an average of 100 cells per meta-cell, which serves as a parameter selection criterion. Using pre-existing cell-type labels, we define the purity of meta-cells by 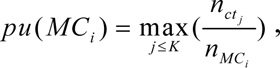 where *MC_i_* denotes the *i*th meta cell, *pu* (*MC_i_*) denotes the purity of *MC_i_*, *n_ct_j__* denotes the cell number of cell type *j* in *MC_i_*, and *n_MC_i__* denotes the cell number of *MC_i_*. The cells contained in meta cells with purity below the given threshold (default 0.95) will be filtered out.

Notably, the meta-purity filtering process does not result in the loss of any predefined cell types (fig. S1, S to U, fig. S3A, and fig. S5B). The meta-purity filtering process has specific designs to prevent the loss of specified cell subsets. We have implemented a stringent criterion for the meta-purity filtering process. If any loss of the specified cell subsets is detected, the process is immediately halted, and displays an error message prompting a review of input parameters. This ensures that there is no unintended loss of any cell types specified by the user.

### Downsampling

Another optional preprocessing for referenced scRNA-seq is down sampling. By default, we will randomly downsample the scRNA-seq reference matrix **B** to 100 cells for each cell type based on a given random seed to improve the speed for downstream analysis. Thus, the processed scRNA-seq expression matrix **B̃** ∊ ℝ*^G^*^×^*^Ñ^* has been established, in which *Ñ* = 100× *K* by default.

### Optimized non-negative matrix factorization (NMF)

In order to efficiently extract the cGEP matrix **E**∊ℝ*^G^*^×^*^K^*, we utilize non-negative matrix factorization (NMF) on the processed scRNA-seq expression matrix **B̃**. Based on the provided cell-type labels, we initialize **H** ∊ℕ*^K^*^×^*^Ñ^* as a one-hot matrix representing the identity of each cell, thus guiding it towards a biologically relevant result. This initialization also aims at reducing variability and improving the consistency between runs. We iterated through NMF to estimate the cGEP matrix **E** :

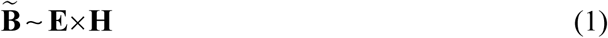

where each element in the matrix **E** and **H** is constrained to be non-negative.

### Entropy-based cGEPs weighting

Considering that the indicative capacity for cell types of each gene is unequal, quantifying the indicative capacity of each gene and taking it into account in the decomposition step would benefit from distinguishing the heterogenous cell states with subtle transcriptome differences. Shannon entropy for the expression percentage of each gene across different cell types can be a great quantification for the indicative capacity of each gene. Thus, we formulate Shannon entropy-based weights for each gene *g* of the extracted cGEP matrix **E** to highlight genes indicative of cell identities.

UCASpatial calculates the expression percentage matrix **PCT**∊ℝ*^K^*^×^*^G^* based on the processed scRNA-seq expression matrix **B̃** :

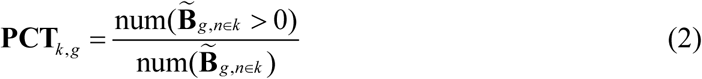

where **PCT***_k_*_, *g*_ denotes the element of **PCT** representing cell type *k* for gene *g,* num(**B̃**_*g*,*n*∊*k*_) represents the number of cells in cell type *k* in **B̃**, num (**B̃** *g*, *n*∊*k* > 0) represents the number of cells in cell type *k* with gene *g* expression over 0. The expression percentage matrix **PCT** will be further normalized into 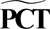 via each element **PCT** being divided by the sum of the corresponding row:

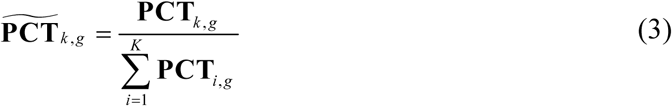

Based on the 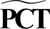 matrix, UCASpatial further calculates the Shannon entropy for each gene:

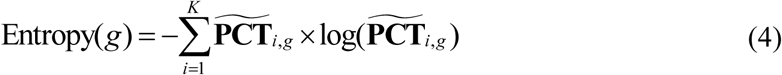

For better understanding, UCASpatial provides an adjusted entropy using the maximum entropy minus the entropy for each gene *g_0_*:

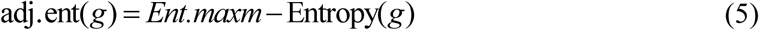

where 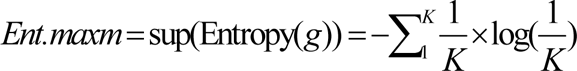 represents the supremum of Entropy(*g*). The maximum entropy *Ent.maxm* is only related to the number of cell types *K*. The adjusted entropy could excellently represent the expression specificity for genes, with a higher adjusted entropy representing a higher indicative capacity for cell types of a gene.

Considering that the adjusted entropy can only evaluate the expression specificity of each gene, we thus took not only the adjusted entropy but also the expression percentage and expression level into account, designing the entropy-based weight of each gene as:

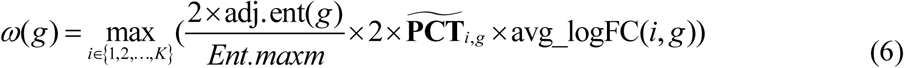

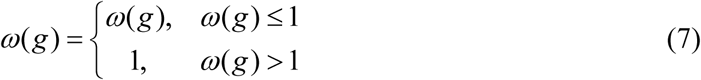

where avg_logFC(*i*, *g*) denotes the average of log_2_ fold-change expression for gene *g* in cell type *i*, *ω*(*g*) denotes the entropy-based weight of gene *g* which is constrained to be non-negative. UCASpatial also combined common marker finding methods including the FindAllMarkers function in Seurat (version 3.1.4) and cosine similarity method ^102^ (version 0.9.0), for optional gene filtering in cGEPs, which can further improve the indicative capacity of genes in cGEPs.

### Entropy weighted non-negative least square model

Taking the untransformed spatial expression count matrix **P** as input, this step aims to decompose the RNA counts of each spot into referenced cell type composition **X**∊ℝ*^S^*^×^*^K^* based on **E**. Unit variance normalization by gene is performed in **P** in order to standardize discretized gene expression levels ^5, 103^. Commonly, this step will utilize the non-negative least square (NNLS) model for each spot *s*:

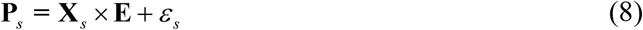

where **P***_s_* denotes the spatial expression count vector of spot *s*, *ε_s_* ℝ *^S^*^×1^ represents the residual error of spot *s* which is restricted to being non-negative. By minimizing the loss function for each spot *s*:

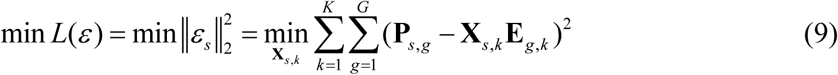

where **X** *_s_* _, *k*_ denotes the cell type composition of spot *s*, with each element restricted to be non-negative; **P***_s_*_,*g*_ denotes the spatial expression count of gene *j* in spot *s*; *ε_s_* denotes the residual error between the deconvoluted expression vector **X** *_s_* _, *k*_ **E** *_g_* _,*k*_ and true spatial expression **P***_s_*_,*g*_ where *j* ∊ {1, 2, …, *G* }. By minimizing the loss function 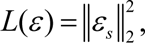 we can obtain the **X** *_s_* _,*i*_ which can best mimic the real cell type composition of spot *s* under the given cell identities.

To further take the entropy-based weight into account, we extended the common NNLS into an entropy-Weighted Non-Negative Least Squares (WNNLS) model. We first extend the weight vector *ω*(*g*) to a diagonal matrix **Ω**∊ℝ*^G^*^×^*^G^* :

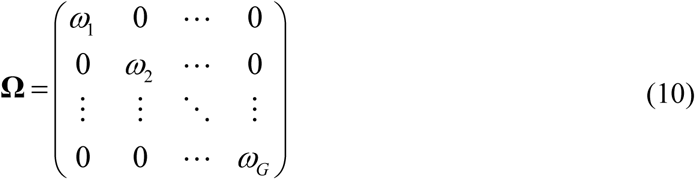

the corresponding WNNLS model has been extended into:

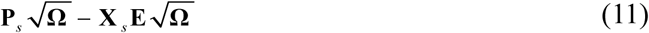

The loss function of the WNNLS model weighted by the diagonal matrix **Ω** is:

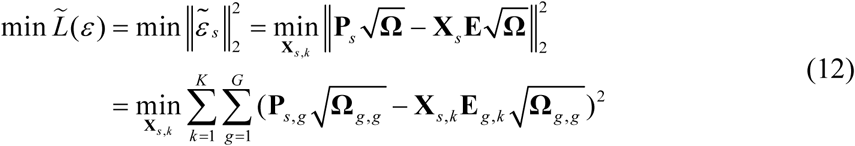

where each element in **X** *_s_* _,*i*_ is restrained to be non-negative. Each row of the cell-type composition spectrum **X** is normalized into [0, 1] by 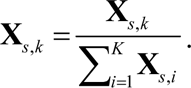

By minimizing the weighted loss function 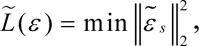 we can highlight the contribution of genes indicative of cell identities, and finally calculate the cell-type composition spectrum **X** in a more biological way.

Referring to SPOTlight, by selecting all cells in **B̃** from the same cell type *k,* and computing the median of each column of **E** for a consensus cell type-specific topic signature, we use **H** to further correct **E**.

Additionally, while UCASpatial is implemented in R, its deconvolution algorithm NMF is implemented with an efficient C++ code that is linked back to the main functions of UCASpatial through Rcpp, ensuring scalable computation.

Finally, considering that the number of cells in each spot is generally within a certain range, for example, 5-20 cells for 10x Visium, we believe that a proportion less than the threshold will be noise in the calculation process. By default, we use a less stringent filter of 0.01 to remove potential noise in the cell-type composition **X**.

### Optional other weighting strategies

In addition to an entropy-based weighting strategy, we implemented alternative weighting strategies to accommodate diverse user requirements and analytical contexts—including mutual information (MI) weighting and variance-based weighting—are provided and described as follows.

Variance-based Weighting: For each gene, mean expression levels were computed across predefined cellular subpopulations (clusters) using normalized scRNA-seq data from the selected assay and data slot. The variance of these cluster-wise mean expressions served as a specificity metric, reflecting the extent of differential expression between subpopulations. These variances were linearly normalized to the [0,1] range to obtain variance-based weights. To integrate gene expression magnitude and percentage, these weights were combined multiplicatively with average log fold change and cluster-wise expression frequency metrics. The final combined weight was bounded to a maximum value of 1.

Mutual Information-based Weighting: To capture nonlinear or complex dependencies between gene expression and subpopulation identity, gene expression was binarized (expressed vs. non-expressed) across cells. Mutual information between this binary expression state and cluster labels was computed using empirical entropy-based estimators, quantifying the reduction in uncertainty of cluster identity given gene expression. Mutual information scores were rescaled to [0,1] and combined with log fold change and expression percentage as above. The resulting weight was capped at 1.

Notably, the weighting strategy can be conveniently adjusted via the *weight.strategy* parameter interface provided in UCASpatial.

### Simulation strategy and evaluation metrics

#### Simulation strategy

We generated simulated spatial transcriptomic data sets from a reference CRC scRNA-seq data set (fig. S1, A to D, see in the *Human specimens (in-house cohort)* part). To simulate different situations of tumor microenvironment, we generated three different regions including tumor core (TC), tumor margin (TM), and tumor stroma (TS) ^17, 47^. Each region has five replicates with each replicate containing *N_spot_* = 50 spots, generated based on random seeds (19980205, 19990911, 19960330, 20000502, 20230509). The number of cells in each spot is restricted to less than 20. To mimic the true situation as much as possible, we also imposed restrictions on the types of cells present in different regions, so that spots from different regions have different cell type co-localization patterns. Detailed restrictions for regional simulation are listed in table S1. To better mimic the real capture, if the synthetic spot had over 25,000 UMI counts, it would be randomly down-sampled into 20,000 UMI counts.

We applied different methods (see in the *Compared methods* part) to deconvolute the above datasets. In each analysis, we provided identical simulated spatial data and reference scRNA-seq data as input for all methods. To ensure an equitable assessment of algorithm performance, we adopted the identical threshold of 0.05 (established on spots containing up to 20 cells) as UCASpatial for algorithms lacking default filtering thresholds, aiming to eliminate noise in accessing the ability to determine whether individual cell types exist at a spot (i.e., spots with non-zero cell abundance or not).

#### Construct complexity hierarchy

To evaluate UCASpatial in microenvironments with different complexity, we rank the complexity hierarchy of these simulated spots by calculating their cell-type composition entropy: 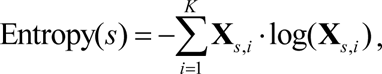 where **X**_*s*, *i*_ denotes the proportion of cell type *i* in spot *s*. We classify them into three groups (low, medium, and high), with *N_spot_* =250 in each group, where higher entropy indicates a higher level of complexity.

#### Evaluation metrics

In each simulation replicate, we calculated the true cell-subpopulation proportions on each spatial location as the number of cells in each cell subpopulation divided by the total number of cells in the location. *T_s_* _,*k*_ and *P_s_*_,*k*_ represent the true and predicted proportion of cell subpopulation *k*, respectively, in the *s*th spot through the number of total cell subpopulations *K*. To evaluate the performance of the methods comprehensively, the main benchmark metrics used were PCC, RMSE, and F1 score. The definitions of these metrics are provided below.

##### PCC

The cell subpopulation-level PCC value was calculated between the predicted proportion of the specific cell subpopulation *P_k_* and the ground truth proportion of the specific cell subpopulation *T_k_* using the following equation:

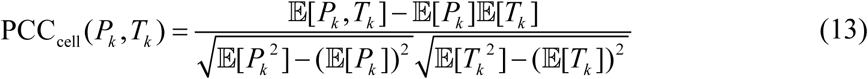

The spot level PCC value was calculated between the predicted spot composition of *P_s_* and the ground truth spot composition *T_s_* using the following equation:

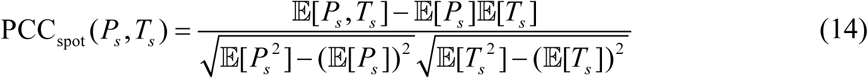

##### RMSE

The cell subpopulation-level RMSE was calculated between the predicted proportion of the specific cell subpopulation *P_k_* and the ground truth proportion of the specific cell subpopulation *T_k_* using the following equation:

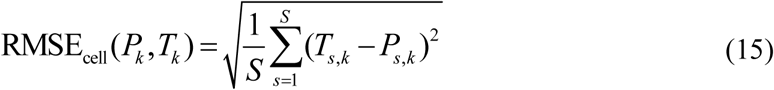

The spot level RMSE value was calculated between the predicted spot composition of *P_i_* and the ground truth spot composition *T_i_* using the following equation:

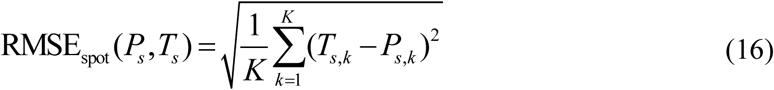

##### F1 score

F1 score was selected to evaluate the ability of alternative methods to determine the presence of individual cell types at specific locations, i.e., with non-zero cell abundances, based on the confusion matrix. A confusion matrix is a table used in the field of machine learning and statistics to assess the performance of a classification model. It summarizes the performance of a classification algorithm by showing the counts of true positive (TP), true negative (TN), false positive (FP), and false negative (FN) predictions for each class in a dataset.

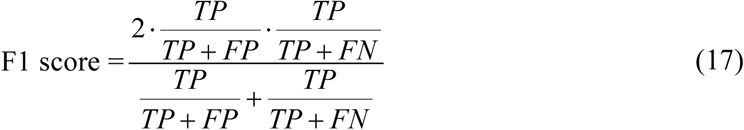

#### Compared methods

We compared UCASpatial with five deconvolution methods: (1) SPOTlight (version 0.1.7), (2) RCTD (version 1.2.0) (3) cell2location (version 0.8a0), (4) CARD (version 1.0) and (5) stereoscope (v 0.3.1). (6) Spotiphy (version 0.1.3). Results of the stereoscope were computed with parameters “-sce 25000 -n 5000 -ste 25000 -stb 500 -scb 500 –gp”. Results of the RCTD were use the ‘full model’. For other methods, we followed the tutorial on the corresponding GitHub pages and used the recommended default parameter settings for deconvolution analysis.

#### Computer platform for evaluating runtime and memory usage

We ran CPU tests of the 7 methods on a computer cluster with an identical CPU platform (2.1 GHz, 60.5 MB L3 cache, 44 CPU cores) and 384 GB memory (DDR4 2,666 MHz). The GPU tests for cell2location and stereoscope were performed on the same computer with an additional GPU (GeForce RTX 4090 with 24 GB memory). To assess the impact of various data attributes (including the number of cells in scRNA-seq data and the number of spots in spatial data) on the computing resources consumed by those 7 methods, we constructed simulated spatial datasets with: 1) varying number of spots (50, 500, 5000, and 50000), 12500 cells, and 20 cell types; 2) varying numbers of cells (approximately 1000, 2500, 5000, 10000, 15000, 25000, and 35000), 20 cell types, and 5000 spots. For the memory evaluation, we used the simulated dataset with 50000 spots ST and about 12500 cells, and 20 cell types. Then, we evaluated the impacts of these data attributes on computing resources consumed.

#### Analysis of human CRC 10x Visium HD data

To test UCASpatial’s ability on the 10x Visium HD platform, we applied UCASpatial to 10x Visium HD data from a human CRC sample ^33^. We used the recommended bin size of 8 μm × 8 μm for this analysis. With the scRNA-seq data shown in Fig. 3B as a reference, we spatially annotated 34 cell subpopulations using the hyper-parameter setting ‘meta.filter = F’ (fig. S2A). For comparison, we also performed RCTD using the ‘full’ model. Other aforementioned methods did not finish the task for the 8μm bin annotation, possibly due to high memory demands, long run times, or software version incompatibility with 10x Visium HD. The high-resolution images of the deconvolution results have been uploaded to GitHub (https://github.com/BIGHanLab/UCASpatial) for better visibility.

In our testing environment (2.1 GHz, 60.5 MB L3 cache, 44 CPU cores), UCASpatial completed the task in just 1.97 hours, while RCTD required 2.27 days for complete annotation. Furthermore, we evaluated UCASpatial on 10x Visium HD data with 2 μm × 2 μm bin size, which was processed in 2.14 hours, demonstrating its scalability to high-resolution data.

#### Analysis of mouse hippocampus Slide-seq V2 data

To test UCASpatial’s ability in Slide-seq V2 platform, we used UCASpatial (meta.resolution = 200, meta.purity = 0.98, cos.filter.threshold = 0.1, ent.filter.threshold = 0.2, logfc.threshold = 0.1, min.diff.pct = 0.05, meta.assay = ‘RNA’) to spatially annotate cell subpopulations in Slide-seq V2 data of the mouse hippocampus using published scRNA-seq datasets of mouse hippocampus^10, 104^.

It is noteworthy that for CARD, we utilized a downsampled scRNA-seq reference dataset consistent with the original CARD paper^12^ to faithfully reproduce its results, whereas other methods were run using the full single-cell reference. Similar to the strategy in the CARD paper, we didn’t apply cell2location on this dataset due to its heavy computational burden. Results of the stereoscope were computed with parameters “-sce 25000 -n 5000 -ste 25000 -stb 500 -scb 500 – gp”. Results of the RCTD were used with the ‘full model’. For other methods, we followed the tutorial on the corresponding GitHub pages and used the recommended default parameter settings for deconvolution analysis. The visualization results of each method can be found in fig. S2H.

### Simulated data generation and method benchmarking based on 10x Visium HD data

Public 10x Visium HD data from human colorectal cancer (CRC) samples were obtained^37^. Cell segmentation and spatial alignment were performed using Space Ranger software (version 4.0.1), and reads were aligned to the hg38 reference genome. Cells were annotated according to their transcriptomic profiles (fig. S2, J and K, and Fig. 3A). To approximate the capture area of a standard Visium spot (55 µm in diameter), we overlaid a grid of 49 µm × 49 µm square bins onto the coordinate system of each tissue section. For each square bin, we aggregated the transcriptomic counts (UMI counts) from all segmented cells whose centroids lay within its boundaries, thereby generating a composite gene expression profile representing a single “pseudo-spot” (fig. S2I).

To evaluate the performance of each deconvolution method on these pseudo-spots, we employed UCASpatial (with meta.filter = FALSE) to spatially annotate cell subpopulations within each pseudo-spot. We did not apply Cell2location to this dataset due to its substantial computational demands. Results from Stereoscope were obtained using the parameters “-sce 500 -n 5000 -ste 500 -stb 500 -scb 500 –gp.” For RCTD, we used the “full model.” For all other methods, we followed the tutorials available on the respective GitHub repositories and applied the recommended default parameter settings for deconvolution analysis.

Detailed deconvolution results of each method are publicly available in the Zenodo repository (see Data Availability).

### Experimental and Analytical Approach to Human CRC

#### Human specimens (in-house cohort)

All patients were enrolled and pathologically diagnosed with colon cancer at Peking University Cancer Hospital & Institute (table S2). Written informed consent was obtained from all patients. This study was approved by the Institutional Ethics Committee of Beijing Cancer Hospital (2023YJZ16). Clinical and pathological stages were determined according to the guidelines of the AJCC version 8. Detailed clinical information was summarized in table S2. Fresh tumor tissues and patient-matched normal adjacent colon tissues were collected from surgical resection specimens, with no prior exposure to chemotherapy or radiation therapy. The resected samples were stored in RPMI 1640 (Gibco) supplemented with HEPES (Thermo Fisher, 15630080), Penicillin-Streptomycin (Thermo Fisher, 15140122), Amphotericin B (Sigma, S1636), and Gentamicin Sulfate (Gibco, 15710-064) and transported at 4°C immediately. As for CRC1, CRC2 and CRC3, a small portion of each tissue was frozen and embedded in OCT (Sakura Tissue-Tek), while the remainder of the same tissue was dissociated into a single-cell suspension.

#### Single-cell RNA-seq for patient samples

Fresh human tumor specimens and patient-matched adjacent normal tissues were washed twice with DPBS (gibco) and then mechanically disrupted into approximately 1 mm^3^ pieces in the RPMI-1640 (gibco) and further enzymatically digested with Human Tumor Dissociation Kit (Miltenyi Biotec, 130-095-929) in C-tube in combination with gentleMACS™ Dissociator (Miltenyi Biotec, 130-093-235) presetting program h_tumor_01, h_tumor_02 and h_tumor_03 sequentially according to the manufacturer’s instruction. Between each program, samples were incubated for 30 minutes at 37℃ under continuous rotation using a rotator. The dissociated cells were subsequently passed through a 70-μm cell strainer (Biologix, 15-1070) and centrifuged at 300g for 7 min. After the supernatant was removed, the cell pellet was washed with DPBS. The pelleted cells were resuspended in a cell cryopreservation medium (FBS with 10% DMSO) at a concentration of 1×10^6^ cells/ml and further cryopreserved in liquid nitrogen for long-term storage.

On the day of scRNA-seq droplet capture, cells were thawed rapidly in a 37℃-water bath and washed off DMSO in DPBS. Palleted cells were resuspended in a sorting buffer (PBS with 2% FBS and 2 mM EDTA). Cells were stained with CD45-FITC (BioLegend Cat# 368508, RRID: AB_2566368) for 20 minutes at 4℃ in the dark. 7-AAD was freshly added prior to FACS sorting performed on the BD FACS Aria III instrument. Based on FACS analysis, 7-AAD^-^CD45^+^ cells, and 7-AAD^-^CD45^-^ cells were sorted into 1.5ml tubes (ExCell Bio, CS015-0041) and counted manually under the microscope (Leica), respectively. Libraries were generated using Chromium Next GEM Single Cell 5’ Library & Gel Bead Kit v1.1 (10x Genomics, PN-1000165) according to the manufacturer’s protocol. Purified libraries were sequenced by the Illumina NovaSeq 6000 platform with 150-bp paired-end reads.

#### 10x Visium spatial transcriptomics

CRC samples were frozen and embedded in OCT (Sakura Tissue-Tek). All tissue blocks were sectioned using the Leica CM1950 cryostat and were cut at 10 μm. Samples were selected on the basis of morphology, orientation (based on H&E staining), and RNA integrity number (RIN) that was obtained using Agilent 2100 Bioanalyzer. The RIN of selected samples ranged from 4.5 to 8.3. The sections were treated as recommended by 10x Genomics and the optimization procedure showed the optimal permeabilization time determined as 12 min for CRC samples using the Visium Spatial Tissue Optimization Kit (10x Genomics, PN-1000193). Spatial gene expression slides were equilibrated inside the cryostat at -20℃ before sectioning and all of the reagents used for spatial transcriptomics were from the Visium Spatial Gene Expression Kit (10x Genomics, PN-1000187). According to the manufacturer’s instruction, the tissue sections were fixed in chilled methanol at -20℃ for 30 min and stained with H&E. Brightfield histological images were performed on the Olympus Microscopes BX53 at ×20 magnification and then taken by cellSens Dimension (version 1.18) software. Images were exported as tiled tiffs for analysis. After imaging, tissue sections were permeabilized at 37℃ for optimal permeabilization time of release of mRNA as mentioned above. Following permeabilization, mRNA was reverse-transcribed and underwent second-strand synthesis, denaturation, cDNA amplification, and cleanup by SPRIselect beads sequentially (Visium, 10x Genomics, User Guide CG000239). cDNA amplification cycles were determined by quantitative PCR (qPCR). Library quality was evaluated using TapeStation (Agilent Technologies). Sequencing was performed on the Illumina HiSeq X Ten platform.

#### Single-cell RNA-seq data processing and annotation

In-house single-cell RNA-seq data from 10x Genomics were processed using Cell Ranger (version 6.0.1), aligning to the hg38 reference genome. Low-quality cells were filtered out if they expressed fewer than 500 or more than 7,000 genes. Cells with mitochondrial gene expression exceeding 20% were also excluded. Batch effects of in-house and public scRNA-seq datasets were corrected using the integration workflow in the Seurat R package (version 3.1.4) ^105^.

To identify and refine cell subpopulations, we employed a multi-step approach. Initially, we employed log-normalization with a scaling factor of 10,000 to adjust for variations in library size among cells. We then identified top 2000 highly variable genes utilizing the Seurat package and conducted Principal component analysis (PCA) on these genes, retaining the top 20 principal components that determined by the elbow method on a Scree plot. Next, we proceeded with an iterative clustering approach to delineate clusters within the single-cell data. Generally, this iterative process initially identified clusters using the Leiden algorithm, followed by a merging of clusters from cell types that appeared to be identical. In the context of certain cell types characterized by heightened heterogeneity, such as T&NK, myeloid, and epithelial cells, we performed cell-type-specific re-integration based on top 2000 variable features to capture more signature genes that can represent the heterogeneity of subpopulations. Then we performed sub-clustering to further identify subpopulations using Leiden clustering. This iterative clustering process allowed us to refine the initial clustering, enabling the splitting of initial clusters that contained multiple subpopulations into more specific lower-level subpopulations. One T cell and myeloid cluster with few specific markers but exhibiting high expression of cell cycle-related genes was removed, as these genes could not be fully regressed out after cell cycle score scaling. In total, we defined 34 cell types in CRC, and this CRC scRNA-seq reference was used throughout the manuscript to maintain consistency in presenting spatial mapping results (Fig. 3B and fig. S3, B to D).

To infer tumor clonal diversity from scRNA-seq data, CNVs were evaluated using the InferCNV v1.2.2 R package (https://github.com/broadinstitute/inferCNV). CNVs of epithelial cells were estimated using non-epithelial cells as a reference. We used a cutoff of 0.1 for the minimum average read counts per gene among reference cells, clustered each group of cells separately, and denoised our output (fig. S5L).

#### Spatial transcriptomics data processing

Raw sequencing reads of spatial transcriptomics were quality-checked and mapped by Space Ranger v1.3.0. The gene-spot matrices generated after ST data processing from ST and Visium samples were analyzed with the Seurat package in R. The processed data were used for downstream analysis of UCASpatial (meta.resolution = 300, meta.purity = 0.9, other parameters using default) and other methods (using default parameters). Using the threshold of 3% to filter the noise after decomposition by each method.

#### Gene signature score calculated in ST

Selected the top 20 genes of each cell subpopulation provided in table S4 or table S6 as the signature gene list for the murine wound healing ST dataset and human CRC ST dataset, respectively. Used the “AddModuleScore” method in the Seurat R package to calculate the signature score of each cell subpopulation in the ST dataset using the signature gene list.

#### Cellular distance evaluation in spatial transcriptomics

In order to quantify the distance relationships between different cell subpopulations in spatial transcriptomics data, we developed an algorithm based on the deconvolution results of UCASpatial. To calculate the spatial distance between cell subpopulation A and cell subpopulation B in spatial transcriptomics data, we first conduct a breadth-first search for each spot containing cell subpopulation A to identify the nearest spot containing cell subpopulation B, and record the distance as well as the proportions of subpopulation A and B within their respective spots. Then we calculate a weighted average on each shortest distance, taking into account the proportions of subpopulation A and subpopulation B within the spots simultaneously. This yields the distribution of spatial distances between the two subpopulations.

#### Multi-color immunohistochemistry (mIHC)

mIHC was performed manually using the Opal 7-Color Manual IHC Kit (AKOYA BIOSCIENCES, Cat# NEL81 1001KT), following the manufacturer’s protocol. Fresh human tumor specimens were excised and fixed in 10% formalin for 24 hours at 4°C. The fixed tissues were then embedded in paraffin and sectioned at 5 µm thickness for histological evaluation. Tissue sections were baked in the oven at 65°C for 1 hour, dewaxed with xylene, and rehydrated through a graded series of alcohol concentrations (100%, 95%, 70%).

Microwave treatment is used in the method to quench endogenous peroxidase activity, for antigen retrieval, and to remove antibodies after a target has been detected. The slide was boiled in the microwave oven for 45 seconds at 100% power and then 15 min at 20% power, followed by blocking for 10 min at room temperature. Primary antibodies against protein CD14 (Proteintech Cat# 60253-1-Ig, 1:200), C1Qc (Proteintech Cat# 16889-1-AP, 1:100), IGF1 (Proteintech Cat# 28530-1-AP, 1:100), SPP1 (Proteintech Cat# 22952-1-AP, 1:200), PDPN (Proteintech Cat# 11629-1-AP, 1:100), ANXA1 (Abcam Cat# ab214486, 1:4000), CXCL13 (Abcam Cat# ab246518, 1:1000), PD-1 (Abcam Cat# ab137132, 1:500), CD4 (Abcam Cat# ab133616, 1:100), FOXP3 (Abcam Cat# ab20034, 1:1000), IL-17A (Proteintech Cat# 26163-1-AP, 1:1000), HLA-DR (Abcam Cat# ab92511, 1:400), CD20 (Abcam Cat# ab78237, 1:250), CLEC9A (Proteintech Cat# 55451-1-AP, 1:200), CD1c (Abcam Cat# ab246520, 1:1000), DC-LAMP (Abcam Cat# ab271053, 1:500), GZMK (Abcam Cat# ab282703, 1:500), CD8 (Abcam Cat# ab17147, 1:100), CD103 (Abcam Cat# ab224202, 1:800), and CTHRC1 (Abcam Cat# ab256458, 1:400) were sequentially applied at 4°C overnight or at room temperature for 1 hour.

The secondary antibody Polymer HRP Ms + Rb (Cat# ARH1001EA) was applied and incubated at room temperature for 10 minutes. Signal amplification was performed using the Opal Fluorophore Working Solution on the slides at room temperature for 10 minutes. Finally, nuclei were stained with DAPI (Invitrogen, Cat# D1306, RRID: AB_2629482). Multi-color IHC imaging was collected using the Vectra Polaris Imaging System (PerkinElmer) ^106^ at 20x magnification and integrated using the InForm Advanced Image Analysis Software (PerkinElmer, version 2.4.1).

Cell segmentation and quantification was carried out using Qupath ^107^. In short, DAPI stains were used for cell detection with the following parameters: requestedPixelSize 1 μm, backgroundRadius 10 μm, sigma 0.5 μm, minArea 10 μm, maxArea 400 μm, threshold 0.1, cellExpansion 1.5μm. Cell subpopulations were classified based on the combinatorial expression of key protein markers (fig. S4C). The distances between CD8^+^ T cell subpopulations and the tumor region were calculated using the “Distance to annotation 2D” function in QuPath. For the pseudo-spot visualization, we first divide the mIHC data into densely distributed bins, each with a width and length of approximately 50 μm. Then, we calculate the proportion of each cell subpopulation in the pseudo-spots by dividing the number of specified cell subpopulations by the total cell count defined by DAPI.

### Tumor clonal inference and clonalTIME quantification using UCASpatial-clonalTIME

To infer tumor clonal diversity within each spatial transcriptomics slice, CNVs were evaluated by ordering genes according to their chromosomal location and applying a moving average to the relative expression values. Genes were arranged according to their genomic localization using the SpatialInferCNV R package ^39^. CNVs in high epithelial spots (overall epithelial proportion > 40%) were estimated using reference spots containing few or no malignant cells selected from UCASpatial annotated remaining spots (overall epithelial proportion < 40%). Considering the non-dispersed spatial distribution of epithelial cells, scattered high epithelial spots (neighboring same-type spots < 1, step = 1) were considered noise and defined as remaining regions. The spot-resolution CNV profiles for each spatial transcriptomics slice were then estimated using the SpatialInferCNV workflow (cutoff = 0.1, cluster_by_groups = FALSE, denoise = TRUE, HMM = FALSE), excluding chrM, chrX and chrY genes. To further define tumor regions and infer tumor clonal diversity based on CNV data, we constructed dendrogram trees to determine clonal groupings of each spot. Dendrogram nodes with at least one high-probability CNV region were defined as tumor clones and assigned a specific identity. Nodes without high-probability CNVs were assigned to normal epithelial or remaining regions based on their histological structure. The average cell subpopulation composition across all spots within a clonal region was defined as the cell subpopulation composition of that clonalTIME (Fig. 3H). To perform a comprehensive comparison of individual CNVs across clones, we initially identified the CNV locations and statuses of each clone using an HMM model in SpatialInferCNV (Fig. 4, A to C, and fig. S5A). CNV regions located on the same chromosome with similar variation patterns in the TCGA-COAD dataset were merged if their PCC (log_2_FC_CNV) > 0.8, where log_2_FC_CNV denotes the log2 fold change of the copy number of the tumor over the copy number of normal. These CNVs were then separated based on the location of the centromere (table S5).

### ClonalTIME phenotyping

To classify the clonalTIME phenotype of each tumor clone, the scaled proportion of the T cell subpopulation of each clone was clustered using the K-means function (centers = 2). ClonalTIMEs were classified as T cell-riched or T cell-excluded based on the proportion of T cell subpopulation within each clustering group (Fig. 3I and fig. S4, J and K). To determine whether a given cell subpopulation was enriched in a specific clonalTIME phenotype, we calculated the ratio of observed cell number to expected cell number (Ro/e) for T cell-rich clonalTIME, where the expected proportion of cell subpopulation was determined from the chi-squared test. We assumed that a subpopulation was enriched in T cell-rich clonalTIME if Ro/e > 1.2 and enriched in T cell-excluded clonalTIME if Ro/e < 0.8.

### Multi-cellular community analysis

To identify cell subpopulation co-localization patterns, we first adjusted the cellular composition of each spot by averaging the proportions of cell subpopulations within that spot and its immediate neighbors (step = 1). Subsequently, we employed the Pearson method to calculate the correlations among the adjusted cell subpopulations across all spots within the tumor region. To identify distinct multi-cellular communities, we applied the ward.D2 hierarchical clustering algorithm to the correlation matrix. This analysis yielded three multicellular communities in the human CRC part, and two multicellular communities in the murine wound healing part (Fig. 3J and Fig. 6D).

The functionality of each multicellular community was characterized based on the shared enriched pathways exhibited by its constituent cell subpopulations. Shared enriched pathways were defined as those that were statistically significantly enriched in at least half of the subpopulations within each community, with a minimum of one subpopulation per community contributing to this enrichment. To address the issue of redundancy in pathway databases, the identified shared enriched pathways were subjected to hierarchical clustering using the pairwise Pearson correlation distance calculated across the gene space. Here, each pathway was represented by its associated differentially expressed genes. Pathway clusters were named manually.

### Multi-cellular community score

The proportions of each cell subpopulation were first normalized by the proportion of non-epithelial cells, ensuring the sum of non-epithelial cell proportions of a spot equals one. To enable inter-cell type comparability, we scaled the proportion of each cell subpopulation using the 99.5th percentile across all tumor spots, resulting in a score for each cell subpopulation. The average score of all cell subpopulations within a community was then calculated as the community score.

### Spatial quantification of TE expression

We quantify the expression of transposable elements (TEs) using spatial transcriptomic data based on a previously reported algorithm ^53^. This algorithm allocates TE reads to TE metagenes based on the TE type-specific sequence (Fig. 4H). In brief, the genes/TEs and cell count matrix were generated according to default parameters and normalized using the NormalizeData function. Clonal-specific TE expression programs were identified by comparing each tumor clone region using the FindAllMarkers function. The list of CRC-specifically upregulated TEs was downloaded from previous literature ^54^. The *HERV-H* expression level was calculated as the average expression of previously reported *HERV-H* elements (*HERVH-int*, *LTR7*, *LTR7B*, and *LTR7Y*) (Fig. 4J).

### Differential expression analysis using spatial transcriptomics

Differentially expressed genes across each clone were analyzed using the Seurat R package (version 4.0.5). Space Ranger output files were imported, and the data were normalized and scaled using the default NormalizeData and ScaleData functions. Differential gene expression analyses were performed comparing groups using the FindMarkers function, with the following parameters: log_2_(fold change) ≥ 0.1 and adjusted P-value ≤ 0.05 (Fig. 4G). The list of CHIP-seq-based STAT1 target genes was downloaded from previous literature ^108^. For differentially expressed genes in the CRC4 slice, Gene Ontology (GO) enrichment analysis was performed by clusterProfiler (v4.6.0) (Fig. 4G) ^109^.

### TCGA-COAD data analysis

Gene expression and clinical data for Colon Adenocarcinoma (COAD) were downloaded from The Cancer Genome Atlas (TCGA). Average copy number data were downloaded from the UCSC Xena browser (https://xena.ucsc.edu). For CNV analysis, only the samples with CNV information were obtained. The average log_2_FC_CNV > 0.35 was defined as copy number gain, while the average log_2_FC_CNV < -0.35 was defined as copy number loss (Fig. 4F). The signatures of each cell subpopulation were obtained in CRC single-cell data by the FindAllMarkers function in the Seurat package with default parameters. The normalized expression of the top 10 signatures in each cell subpopulation was used to calculate the average signature score. The reads per million (RPM) of known human endogenous retroviruses (HERV) in TCGA-COAD were downloaded from a previous report ^110, 111^. For HERV analysis, samples without HERV expression information were excluded. Overall *HERV-H* expression was calculated as the average expression of known transcribed *HERV-H* loci (*HERV-H1-7*) (Fig. 4K) ^110^. For chromosomal instability assessment, samples were stratified into tertiles by aneuploidy score, followed by comparative analysis of T cell marker expression across groups. The pathway activity of each sample was calculated using GSVA with default parameter settings (fig. S5K) ^112^. Survival analyses were then performed with the survival R package (version 3.4-0) (Fig. 4L and fig. S5R).

### MSK-IMPACT data analysis

The genomic and clinical data of 71 CRC patients with Immune Checkpoint Blockade (ICB) therapy were downloaded from published studies ^57, 58^. The copy number status of each loci was determined using the GISTIC v2.0 function in the GenePattern platform ^113, 114^. The average log_2_FC_CNV > 0.35 was defined as copy number gain, while the average log_2_(copy number/2) < -0.35 was defined as copy number loss. Survival analyses were then performed with the survival R package (Fig. 4L and fig. S5, P to Q).

### Experimental and Analytical Approach to Mouse Ear

#### Ear wound creation and treatment

MRL/MpJ (MRL) mice (RRID: IMSR_JAX:000486) and C57BL/6 (B6) mice (RRID: MGI:2159769) were used for ear wound creation. All mice utilized in the wounding experiments were aged between 8-10 weeks. Mice were subjected to short-term anesthesia induced by isoflurane prior to wound generation. The dorsal and ventral aspects of mouse ears were sanitized with cotton balls saturated in 75% alcohol. Mouse ears were delicately flattened using forceps, and a tissue-penetrating ear puncher (2 mm) was used to create perforating injuries within the vascularized regions of the ear. Hemostatic powder was applied to the wounds using disposable cotton swabs. All instruments were sterilized under high pressure before use.

To block the differentiation of *Fmod*^+^ fibroblasts, mice were either locally injected with rosiglitazone (3 mg/kg) every other day post-injury or smeared with rosiglitazone dissolved in petrolatum every day. For the injection administration, the control group received an equivalent volume of DMSO. For the topical administration, the control group received petrolatum containing DMSO.

To block IL11-IL11RA signaling, mice were locally injected with anti-IL11RA antibody (2.5 µg/wound) at day 3 post-injury. The control group received an equivalent volume of anti-rabbit IgG. All mouse procedures were approved by the Institutional Animal Care and Use Committees of Beijing Institute of Genomics, Chinese Academy of Sciences (2019A010).

### Flow cytometry

Single-cell suspensions were obtained from the ear tissues of B6 and MRL mice at day 7 post-injury. Specifically, mouse ear tissues were mechanically minced and then incubated at 37°C for 5-10 minutes with 40 µL collagenase II (50 mg/mL) and 40 µL collagenase IV (50 mg/mL). Following a brief low-speed centrifugation to precipitate undigested tissue fragments, the supernatant was transferred into a stop solution composed of fetal bovine serum, EDTA, and D-HBSS. Additional digestion enzymes were added to the partially digested tissue fragments, and the digestion process was repeated until complete tissue digestion was achieved. The digested mixture was filtered through a 70-µm membrane and centrifuged at 4°C at 1200 rpm for 10 minutes. After discarding the supernatant, the cell pellet was washed in FACS buffer (PBS supplemented with 2% FBS and 1 mM EDTA), and then resuspended in FACS buffer, and incubated with Fc Block (clone 2.4G2; BioX Cell) for 10 minutes. Subsequently, the indicated antibodies were added, and staining was continued for 30 minutes on ice in the dark. Flow cytometry was performed using a BD Symphony A5 cytometer for analysis and a BD Fusion cell sorter for sorting experiments. The flow cytometry results were analyzed using FlowJo™ v10 Software (BD Life Sciences).

### Multi-color immunohistochemistry (mIHC)

At predetermined time points, murine ears were excised and fixed in 10% formalin for 48 hours at 4°C. The fixed tissues were then embedded in paraffin and sectioned at 7 µm thickness for histological evaluation. Tissue sections were then deparaffinized, rehydrated and stained using 7-color multispectral Opal reagents (AKOYA BIOSCIENCES, Cat# number NEL81 1001KT).

Heat-mediated antigen retrieval was performed using a microwave for rehydrated tissue sections. Primary antibodies against CD11b (Abcam Cat# ab133357, clone EPR1344, 1:500), IL-1β polyclonal antibody (Invitrogen, Cat# P420B, 1:200), CD163 (Abcam Cat# ab182422, clone EPR19518, 1:1000), CD68 (Proteintech Cat# 28058-1-AP, 1:2000), CD74 (Abcam Cat# ab289885, 1:2000), GPNMB (Proteintech Cat# 66926-1-Ig, 1:200), α-SMA(Abcam Cat# ab124964, 1:2000), PDGFR alpha (PDGFRα, Abcam Cat# ab96569, 1:500), CD34 (Invitrogen, Cat# MA5-29674, 1:2500), OGN (Proteintech Cat# 12755-1-AP, 1:200), SOX9 (CST Cat# 82630T, 1:500), Collagen Type II (Proteintech Cat# 28459-1-AP, 1:200), IGFBP5 (Proteintech Cat# 55205-1-AP, 1:200), PRG4 (Proteintech Cat# 25732-1-AP, 1:200), K14 (Santa Cruz Cat# sc-53253, 1:300), CD31 (Abcam Cat# ab281583, 1:2000), FcεRIα (Proteintech Cat#10980-1-AP, 1:50), Ly6G (Abclonal Cat# A22270, 1:200), IL11RA (Invitrogen, Cat# PA5-47667, 1:500) and Osteopontin (Proteintech Cat# 22952-1-AP, 1:500) were individually applied.

The secondary antibody Polymer HRP Ms + Rb (Cat# ARH1 001EA) was applied and incubated at room temperature for 10 minutes. Signal amplification was performed using the Opal Fluorophore Working Solution on the slides at room temperature for 10 minutes. Finally, nuclei were stained with DAPI (Invitrogen, Cat# D1306, RRID: AB_2629482). mIHC imaging was collected using the Vectra Polaris Imaging System (PerkinElmer) ^106^ at 20x magnification and analyzed using the InForm Advanced Image Analysis Software (PerkinElmer, version 2.4.1).

Cell segmentation and quantification were carried out using Qupath ^107^. In short, DAPI stains were used for cell detection with the following parameters: requestedPixelSize 0.5 μm, backgroundRadius 8 μm, sigma 0.5 μm, minArea 10 μm, maxArea 400 μm, threshold 0.5, cellExpansion 5 μm. Cell subpopulations were identified by training an object classifier based on the combination of the marker’s expression. In the epidermis, keratinocytes were measured, while in the dermis, a comprehensive identification and quantification of other cellular subpopulations were carried out. For the pseudo-spot visualization, we first divide the mIHC data into densely distributed bins, each with a width and length of approximately 50 μm. Then, we calculate the proportion of each cell subpopulation in the pseudo-spots by dividing the number of specified cell subpopulations by the total cell count defined by DAPI.

### Single-cell RNA-seq

Single-cell suspensions were obtained from B6 and MRL mouse ear tissue at day 7 post-wounding. Calcein and 7-AAD were freshly added before cell sorting. Then FITC^+^ 7-AAD^-^ single cells were sorted for library construction of scRNA-seq. The libraries were performed using Chromium Next GEM Single Cell 3’ GEM, Library & Gel Bead Kit V3 (PN-1000075) purchased from 10x Genomics according to the protocols provided by manufacturers. The libraries were sequenced on an Illumina Novaseq 6000 platform.

### Spatial gene expression assay

Fresh mouse ear tissues were embedded in the Optimal Cutting Temperature Compound (OCT, Sakura Tissue-TEK), and stored at -80°C. The 10-μm section was placed on the pre-chilled Optimization slides (Visium, 10x Genomics, PN-1000193). After collecting histological morphological information, the tissue was processed within the optimal permeabilization time of 18 minutes to digest and release mRNA from tissue slides. The released mRNA was subsequently captured by proximal oligonucleotides within the tissue, culminating in the generation of the respective libraries. The libraries were sequenced on an Illumina HiSeq X Ten platform.

### Single-cell RNA-seq data processing

10x Genomics scRNA-seq data were aligned and quantified using the Cell Ranger Single-Cell Software Suit (version 3.0.2) against the mm10 reference genome. Genes with fewer detected cell numbers (fewer than 3) and cells with fewer detected genes (fewer than 200) were removed. To further filter out low-quality data, the following criteria were applied to filter cells: (1) the number of expressed genes greater than 500 or less than 5,000; (2) mitochondrial proportion less than 20%. To cluster single cells by their expression, the unsupervised graph-based clustering algorithm implemented in the Seurat R package (version 3.1.4) was used. The top 2,000 variable genes identified by the vst method were used to perform principal component analysis. The RunUMAP and RunTSNE functions were employed to generate UMAP and t-SNE dimensionality reduction results, while the FindNeighbors function was used to construct a shared nearest neighbor graph. The DoubletFinder was utilized to predict potential doublets and filter them out.

### Spatial transcriptomics data processing

Using the Space Ranger software (https://support.10xgenomics.com/spatial-gene-expression/software/pipelines/latest/tutorials), the 10x Visium spatial transcriptomics data was aligned and quantified against the mm10 version of the mouse reference genome. Downstream analysis was conducted using the R package Seurat. Integration of unpaired scRNA-seq data with spatial transcriptomics sequencing data was performed for deconvolution analysis to obtain the proportion of each cell subtype within each spot of the spatial transcriptome. Parameters for UCASpatial: downsample_n = 1000, meta.resolution = 200, meta.filter = 0.95, cos.filter.threshold = 0.1, the others use the default. Parameters for other methods: Default. Using the threshold of 3% to filter the noise after decomposition by each method (fig. S6C).

### Spatial transcriptomics data visualization

The original data placed mouse ear tissues on two slides: tissues from day 0, day 7, and day 15 were on the first slide, and tissues from day 3 were on the second slide. To better visualize the spatial transcriptomics data of mouse ears, we merged all the tissues into one slide and aligned them based on the time point post-wounding. Specifically, we first extracted the coordinates of each spot in all 4 tissues. We then transformed the coordinates of spots from day 3, with each row replaced by the column, and each column replaced by the row. We next transformed the coordinates of each spot’s columns, abiding by the time points. ggplot2 was used for visualizing spatial transcriptomics data after coordinate transformation.

The murine ear tissues have been annotated into four groups of regions based on H&E staining: unwounded (day 0), wound bed (1 mm radius), wound proximal (0-1 mm from the wound bed), and wound distal (> 1 mm from the wound bed).

### Functional enrichment analysis in murine wound healing data

The signatures of macrophage subpopulations, *Fmod*^+^ fibroblast, and *Igfbp5*^+^ chondrocyte in the murine wound healing dataset were obtained in scRNA-seq data by the FindAllMarkers function in the Seurat package with min.pct = 0.1, min.diff.pct = 0.1, logfc.threshold = 0.25 and other default parameters. Metascape was used to perform functional enrichment analysis for *Fmod*^+^ fibroblast and *Igfbp5*^+^ chondrocyte (Fig. S7, L and M) ^115^. The enrichKEGG in the clusterProfiler package was used to perform functional enrichment analysis for macrophage subpopulations (fig. S6N) ^109^.

### Trajectory interference for chondrocyte subpopulations

Monocle3 was used to infer the trajectory of chondrocytes in scRNA-seq data from the murine wound healing model (fig. S7I) ^116-119^. The top 20 PCA defined in scRNA-seq of chondrocytes were used. The reduce_dimension function was used to reduce the dimensions of chondrocytes (Parameters: umap.metric = “cosine”, umap.min_dist = 0.05, umap.n_neighbors = 20). cluster_cells function with parameters cluster_method = “Louvain”, resolution = 0.001, random_seed = 123 was used to cluster the cells. The learn_graph function with default parameters was used to learn the principal graph from the reduced dimension space using reversed graph embedding.

### Ligand and receptor interaction analysis

To identify the significant and specific ligand-receptor pairs between *Igfbp5*^+^ chondrocyte, *Fmod*^+^ fibroblast, and *Cd36^+^ Gpnmb^+^ Il1b^-^* macrophage, the CellPhoneDB tool was used in scRNA-seq data from murine wound healing ^120, 121^. Cell expression was normalized by dividing the total counts. The interaction pairs with P-value < 0.01 or the mean attraction strength > 0 were retained.

## Data availability

All data are available in the main text or the supplementary materials. The sequencing data of both scRNA-seq and spatial transcriptomics of murine wound healing have been deposited in the Genome Sequence Archive (GSA) repository with the accession number CRA016052. The sequencing data of human CRC has been deposited in the Genome Sequence Archive for human (GSA-human) repository with the accession number HRA007218. The public datasets used in this study include the scRNA-seq dataset of human CRC (HRA000979), the 10x Visium spatial transcriptomic dataset of human CRC (HRA000979), and the mouse hippocampus Slide-seqV2 dataset (https://singlecell.broadinstitute.org/single_cell/study/SCP948/robust-decomposition-of-cell-type-mixtures-in-spatial-transcriptomics). The high-resolution deconvolution results of each method for human CRC, murine wound healing, and 10x HD have been uploaded to GitHub: https://github.com/BIGHanLab/UCASpatial/tree/main/Reproduction/High-res%20Figure. The deconvolution results of Visium HD-derived pseudo-spots and the data and code to reproduce the results in this study are available in the Zenodo repositories at https://zenodo.org/records/12634581?preview=1&token=eyJhbGciOiJIUzUxMiJ9.eyJpZCI6ImU0OGM4YjBjLTliYzQtNDc4ZC05Njg3LTczYTdhOTExNTliNiIsImRhdGEiOnt9LCJyYW5kb20iOiI0ZjYyYzU1OGYwNzc4M2E2NDQ5YWFiMjZkZmY2OGE4NCJ9.qcl1aa2yo6mmdkZXdHw2exPmfhyA7M5oPS1ZoKXbqVpBdQbSWkagYIGL2U3Dhj2ffSde0rhe5MFiuIRYuPC_Vw.

## Code availability

The UCASpatial is available in the GitHub repository https://github.com/BIGHanLab/UCASpatial. The data and code to reproduce the results in this study are available in the Zenodo repositories at https://zenodo.org/records/12634581?preview=1&token=eyJhbGciOiJIUzUxMiJ9.eyJpZCI6ImU0OGM4YjBjLTliYzQtNDc4ZC05Njg3LTczYTdhOTExNTliNiIsImRhdGEiOnt9LCJyYW5kb20iOiI0ZjYyYzU1OGYwNzc4M2E2NDQ5YWFiMjZkZmY2OGE4NCJ9.qcl1aa2yo6mmdkZXdHw2exPmfhyA7M5oPS1ZoKXbqVpBdQbSWkagYIGL2U3Dhj2ffSde0rhe5MFiuIRYuPC_Vw.

## Supplementary Information

Figs. S1 to S7 for multiple supplementary figures.

Tables S1 to S6 for multiple supplementary tables.

## Funding

National Key R&D Program of China (2024YFC3405901 to D.H.)

Strategic Priority Research Program of the Chinese Academy of Science (XDB0570101 to D.H.)

National Natural Science Foundation of China (NSFC) (T2495272, 32121001, 22293052)

Beijing Natural Science Foundation (L244023, Z200023)

National Key R&D Program of China (2024YFA1802102)

Next-Generation Bioinformatics Algorithms (XDA0460302)

National Funded Postdoctoral Researcher program (Y.Z.)

CAS Youth Interdisciplinary Team

## Author contributions

Conceptualization: D.H., M.M.X., Yin XU

Methodology: Yin XU

Software: Yin XU, Z.H.

Formal analysis: Yin XU, Z.H., Y.Z., F.Z.

Investigation: Yin XU, Z.H., Y.Z., M.G., P.G., F.Z., J.Y., R.C., G.L., L.D., Yu XIA

Resources: Z.W., L.S., J.L.

Visualization: Yin XU, Z.H., Y.Z.

Project administration: D.H., M.M.X.

Supervision: D.H., M.M.X., Yin XU, Z.H., Y.Z.

Writing – original draft: D.H., M.M.X., Yin XU, Z.H., Y.Z.

Writing – review & editing: H.N., W.G., B.M., Y.G., Z.L.

Funding acquisition: D.H., M.M.X., Y.Z.

## References

1. Brbic, M. et al. Annotation of spatially resolved single-cell data with STELLAR. Nat Methods 19, 1411–1418 (2022).

2. Vandereyken, K., Sifrim, A., Thienpont, B. & Voet, T. Methods and applications for single-cell and spatial multi-omics. Nat Rev Genet 24, 494–515 (2023).

3. Stahl, P.L. et al. Visualization and analysis of gene expression in tissue sections by spatial transcriptomics. Science 353, 78–82 (2016).

4. Junker, J.P. et al. Genome-wide RNA Tomography in the zebrafish embryo. Cell 159, 662–675 (2014).

5. Rodriques, S.G. et al. Slide-seq: A scalable technology for measuring genome-wide expression at high spatial resolution. Science 363, 1463–1467 (2019).

6. Vickovic, S. et al. High-definition spatial transcriptomics for in situ tissue profiling. Nat Methods 16, 987–990 (2019).

7. Liao, J., Lu, X., Shao, X., Zhu, L. & Fan, X. Uncovering an Organ’s Molecular Architecture at Single-Cell Resolution by Spatially Resolved Transcriptomics. Trends Biotechnol 39, 43–58 (2021).

8. Rao, A., Barkley, D., Franca, G.S. & Yanai, I. Exploring tissue architecture using spatial transcriptomics. Nature 596, 211–220 (2021).

9. Elosua-Bayes, M., Nieto, P., Mereu, E., Gut, I. & Heyn, H. SPOTlight: seeded NMF regression to deconvolute spatial transcriptomics spots with single-cell transcriptomes. Nucleic Acids Res 49, e50 (2021).

10. Cable, D.M. et al. Robust decomposition of cell type mixtures in spatial transcriptomics. Nat Biotechnol 40, 517–526 (2022).

11. Kleshchevnikov, V. et al. Cell2location maps fine-grained cell types in spatial transcriptomics. Nat Biotechnol 40, 661–671 (2022).

12. Ma, Y. & Zhou, X. Spatially informed cell-type deconvolution for spatial transcriptomics. Nat Biotechnol 40, 1349–1359 (2022).

13. Andersson, A. et al. Single-cell and spatial transcriptomics enables probabilistic inference of cell type topography. Commun Biol 3, 565 (2020).

14. Yang, J. et al. Spotiphy enables single-cell spatial whole transcriptomics across an entire section. Nat Methods 22, 724–736 (2025).

15. Li, B. et al. Benchmarking spatial and single-cell transcriptomics integration methods for transcript distribution prediction and cell type deconvolution. Nat Methods 19, 662–670 (2022).

16. Li, H. et al. A comprehensive benchmarking with practical guidelines for cellular deconvolution of spatial transcriptomics. Nat Commun 14, 1548 (2023).

17. Sun, D., Liu, Z., Li, T., Wu, Q. & Wang, C. STRIDE: accurately decomposing and integrating spatial transcriptomics using single-cell RNA sequencing. Nucleic Acids Res 50, e42 (2022).

18. Kuppe, C. et al. Spatial multi-omic map of human myocardial infarction. Nature 608, 766–777 (2022).

19. Ackerman, J.E. et al. Defining the spatial-molecular map of fibrotic tendon healing and the drivers of Scleraxis-lineage cell fate and function. Cell Rep 41, 111706 (2022).

20. Anderson, A.C., et al. Spatial transcriptomics. Cancer Cell 40, 895–900 (2022).

21. Foster, D.S. et al. Multiomic analysis reveals conservation of cancer-associated fibroblast phenotypes across species and tissue of origin. Cancer Cell 40, 1392–1406 e1397 (2022).

22. Heiser, C.N. et al. Molecular cartography uncovers evolutionary and microenvironmental dynamics in sporadic colorectal tumors. Cell 186, 5620–5637 e5616 (2023).

23. Matusiak, M., et al. Spatially Segregated Macrophage Populations Predict Distinct Outcomes In Colon Cancer. Cancer Discov (2024).

24. Kersten, K. et al. Spatiotemporal co-dependency between macrophages and exhausted CD8(+) T cells in cancer. Cancer Cell 40, 624–638 e629 (2022).

25. van der Leun, A.M., Thommen, D.S. & Schumacher, T.N. CD8(+) T cell states in human cancer: insights from single-cell analysis. Nat Rev Cancer 20, 218–232 (2020).

26. Kirschenbaum, D. et al. Time-resolved single-cell transcriptomics defines immune trajectories in glioblastoma. Cell 187, 149–165 e123 (2024).

27. Benotmane, J.K. et al. High-sensitive spatially resolved T cell receptor sequencing with SPTCR-seq. Nat Commun 14, 7432 (2023).

28. Liu, S. et al. Spatial maps of T cell receptors and transcriptomes reveal distinct immune niches and interactions in the adaptive immune response. Immunity 55, 1940–1952 e1945 (2022).

29. Delfini, M., Stakenborg, N., Viola, M.F. & Boeckxstaens, G. Macrophages in the gut: Masters in multitasking. Immunity 55, 1530–1548 (2022).

30. Xun, Z. et al. Reconstruction of the tumor spatial microenvironment along the malignant-boundary-nonmalignant axis. Nat Commun 14, 933 (2023).

31. Zhang, D. et al. Inferring super-resolution tissue architecture by integrating spatial transcriptomics with histology. Nat Biotechnol (2024).

32. Fu, T. et al. Spatial architecture of the immune microenvironment orchestrates tumor immunity and therapeutic response. J Hematol Oncol 14, 98 (2021).

33. Michelli F. Oliveira, J.P.R., Meii Chung, Stephen Williams, Andrew D. Gottscho, Anushka Gupta, Susan E. Pilipauskas, Syrus Mohabbat, Nandhini Raman, David Sukovich, David Patterson, Visium HD Development Team, Sarah E. B. Taylor Characterization of immune cell populations in the tumor microenvironment of colorectal cancer using high definition spatial profiling. biorxiv 06.04.597233 (2024).

34. Del Bigio, M.R. Ependymal cells: biology and pathology. Acta Neuropathol 119, 55–73 (2010).

35. Ramachandran, V.S. (2016).

36. Meyer, G. Building a human cortex: the evolutionary differentiation of Cajal-Retzius cells and the cortical hem. J Anat 217, 334–343 (2010).

37. Oliveira, M.F. et al. High-definition spatial transcriptomic profiling of immune cell populations in colorectal cancer. Nat Genet 57, 1512–1523 (2025).

38. Seferbekova, Z., Lomakin, A., Yates, L.R. & Gerstung, M. Spatial biology of cancer evolution. Nat Rev Genet 24, 295–313 (2023).

39. Erickson, A. et al. Spatially resolved clonal copy number alterations in benign and malignant tissue. Nature 608, 360–367 (2022).

40. Lomakin, A. et al. Spatial genomics maps the structure, nature and evolution of cancer clones. Nature 611, 594–602 (2022).

41. Qi, J. et al. Single-cell and spatial analysis reveal interaction of FAP(+) fibroblasts and SPP1(+) macrophages in colorectal cancer. Nat Commun 13, 1742 (2022).

42. Trimm, E. & Red-Horse, K. Vascular endothelial cell development and diversity. Nat Rev Cardiol 20, 197–210 (2023).

43. Li, G., Gao, J., Ding, P. & Gao, Y. The role of endothelial cell-pericyte interactions in vascularization and diseases. J Adv Res (2024).

44. Fridman, W.H. et al. B cells and tertiary lymphoid structures as determinants of tumour immune contexture and clinical outcome. Nat Rev Clin Oncol 19, 441–457 (2022).

45. Munoz-Erazo, L., Rhodes, J.L., Marion, V.C. & Kemp, R.A. Tertiary lymphoid structures in cancer - considerations for patient prognosis. Cell Mol Immunol 17, 570–575 (2020).

46. Danenberg, E. et al. Breast tumor microenvironment structures are associated with genomic features and clinical outcome. Nat Genet 54, 660–669 (2022).

47. Dhainaut, M. et al. Spatial CRISPR genomics identifies regulators of the tumor microenvironment. Cell 185, 1223–1239 e1220 (2022).

48. Ramirez, C.F.A. et al. Cancer cell genetics shaping of the tumor microenvironment reveals myeloid cell-centric exploitable vulnerabilities in hepatocellular carcinoma. Nat Commun 15, 2581 (2024).

49. Chen, X., Agustinus, A.S., Li, J., DiBona, M. & Bakhoum, S.F. Chromosomal instability as a driver of cancer progression. Nat Rev Genet 26, 31–46 (2025).

50. Honda, K. et al. IRF-7 is the master regulator of type-I interferon-dependent immune responses. Nature 434, 772–777 (2005).

51. Liu, S.Y., Sanchez, D.J., Aliyari, R., Lu, S. & Cheng, G. Systematic identification of type I and type II interferon-induced antiviral factors. Proc Natl Acad Sci U S A 109, 4239–4244 (2012).

52. Chiappinelli, K.B. et al. Inhibiting DNA Methylation Causes an Interferon Response in Cancer via dsRNA Including Endogenous Retroviruses. Cell 162, 974–986 (2015).

53. He, J. et al. Identifying transposable element expression dynamics and heterogeneity during development at the single-cell level with a processing pipeline scTE. Nat Commun 12, 1456 (2021).

54. Kong, Y. et al. Transposable element expression in tumors is associated with immune infiltration and increased antigenicity. Nat Commun 10, 5228 (2019).

55. Sexton, C.E., Tillett, R.L. & Han, M.V. The essential but enigmatic regulatory role of HERVH in pluripotency. Trends Genet 38, 12–21 (2022).

56. Tumeh, P.C. et al. PD-1 blockade induces responses by inhibiting adaptive immune resistance. Nature 515, 568–571 (2014).

57. Samstein, R.M. et al. Tumor mutational load predicts survival after immunotherapy across multiple cancer types. Nat Genet 51, 202–206 (2019).

58. Zehir, A. et al. Mutational landscape of metastatic cancer revealed from prospective clinical sequencing of 10,000 patients. Nat Med 23, 703–713 (2017).

59. Eming, S.A., Martin, P. & Tomic-Canic, M. Wound repair and regeneration: mechanisms, signaling, and translation. Sci Transl Med 6, 265sr266 (2014).

60. Hu, K.H. et al. Transcriptional space-time mapping identifies concerted immune and stromal cell patterns and gene programs in wound healing and cancer. Cell Stem Cell 30, 885–903 e810 (2023).

61. Mascharak, S. et al. Multi-omic analysis reveals divergent molecular events in scarring and regenerative wound healing. Cell Stem Cell 29, 315–327 e316 (2022).

62. McBrearty, B.A., Clark, L.D., Zhang, X.M., Blankenhorn, E.P. & Heber-Katz, E. Genetic analysis of a mammalian wound-healing trait. Proc Natl Acad Sci U S A 95, 11792–11797 (1998).

63. Clark, L.D., Clark, R.K. & Heber-Katz, E. A new murine model for mammalian wound repair and regeneration. Clin Immunol Immunopathol 88, 35–45 (1998).

64. Heydemann, A. The super super-healing MRL mouse strain. Front Biol (Beijing) 7, 522–538 (2012).

65. Wynn, T.A. & Vannella, K.M. Macrophages in Tissue Repair, Regeneration, and Fibrosis. Immunity 44, 450–462 (2016).

66. Kim, M.H. et al. Dynamics of neutrophil infiltration during cutaneous wound healing and infection using fluorescence imaging. J Invest Dermatol 128, 1812–1820 (2008).

67. Karasuyama, H., Miyake, K., Yoshikawa, S. & Yamanishi, Y. Multifaceted roles of basophils in health and disease. J Allergy Clin Immunol 142, 370–380 (2018).

68. Phillipson, M. & Kubes, P. The Healing Power of Neutrophils. Trends Immunol 40, 635–647 (2019).

69. Raziyeva, K. et al. Immunology of Acute and Chronic Wound Healing. Biomolecules 11 (2021).

70. Hill, D.A. et al. Distinct macrophage populations direct inflammatory versus physiological changes in adipose tissue. Proc Natl Acad Sci U S A 115, E5096–E5105 (2018).

71. Jaitin, D.A. et al. Lipid-Associated Macrophages Control Metabolic Homeostasis in a Trem2-Dependent Manner. Cell 178, 686–698 e614 (2019).

72. Henlon, Y. et al. Single-cell analysis identifies distinct macrophage phenotypes associated with prodisease and proresolving functions in the endometriotic niche. Proc Natl Acad Sci U S A 121, e2405474121 (2024).

73. Franzen, L. et al. Mapping spatially resolved transcriptomes in human and mouse pulmonary fibrosis. Nat Genet 56, 1725–1736 (2024).

74. Abedini, A. et al. Single-cell multi-omic and spatial profiling of human kidneys implicates the fibrotic microenvironment in kidney disease progression. Nat Genet 56, 1712–1724 (2024).

75. Buechler, M.B. et al. Cross-tissue organization of the fibroblast lineage. Nature 593, 575–579 (2021).

76. Konieczny, P. et al. Interleukin-17 governs hypoxic adaptation of injured epithelium. Science 377, eabg9302 (2022).

77. Gawriluk, T.R. et al. Comparative analysis of ear-hole closure identifies epimorphic regeneration as a discrete trait in mammals. Nat Commun 7, 11164 (2016).

78. Pastar, I. et al. Epithelialization in Wound Healing: A Comprehensive Review. Adv Wound Care (New Rochelle) 3, 445–464 (2014).

79. Rousselle, P., Braye, F. & Dayan, G. Re-epithelialization of adult skin wounds: Cellular mechanisms and therapeutic strategies. Adv Drug Deliv Rev 146, 344–365 (2019).

80. Pena, O.A. & Martin, P. Cellular and molecular mechanisms of skin wound healing. Nat Rev Mol Cell Biol 25, 599–616 (2024).

81. Qu, Y. et al. A comprehensive analysis of single-cell RNA transcriptome reveals unique SPP1+ chondrocytes in human osteoarthritis. Comput Biol Med 160, 106926 (2023).

82. Chau, M. et al. The synovial microenvironment suppresses chondrocyte hypertrophy and promotes articular chondrocyte differentiation. NPJ Regen Med 7, 51 (2022).

83. Ignatyeva, N., Gavrilov, N., Timashev, P.S. & Medvedeva, E.V. Prg4-Expressing Chondroprogenitor Cells in the Superficial Zone of Articular Cartilage. Int J Mol Sci 25 (2024).

84. Massengale, M., et al. Adult Prg4+ progenitors repair long-term articular cartilage wounds in vivo. JCI Insight 8 (2023).

85. Diekman, B.O. et al. Cartilage tissue engineering using differentiated and purified induced pluripotent stem cells. Proc Natl Acad Sci U S A 109, 19172–19177 (2012).

86. Ono, N., Ono, W., Nagasawa, T. & Kronenberg, H.M. A subset of chondrogenic cells provides early mesenchymal progenitors in growing bones. Nat Cell Biol 16, 1157–1167 (2014).

87. Yue, B. Biology of the extracellular matrix: an overview. J Glaucoma 23, S20–23 (2014).

88. Burgess, H.A. et al. PPARgamma agonists inhibit TGF-beta induced pulmonary myofibroblast differentiation and collagen production: implications for therapy of lung fibrosis. Am J Physiol Lung Cell Mol Physiol 288, L1146–1153 (2005).

89. Hua, Q. et al. PPARgamma mediates the anti-pulmonary fibrosis effect of icaritin. Toxicol Lett 350, 81–90 (2021).

90. Kokeny, G., Calvier, L. & Hansmann, G. PPARgamma and TGFbeta-Major Regulators of Metabolism, Inflammation, and Fibrosis in the Lungs and Kidneys. Int J Mol Sci 22 (2021).

91. Zhou, X. et al. Circuit Design Features of a Stable Two-Cell System. Cell 172, 744–757 e717 (2018).

92. Takawale, A. et al. Tissue Inhibitor of Matrix Metalloproteinase-1 Promotes Myocardial Fibrosis by Mediating CD63-Integrin beta1 Interaction. Hypertension 69, 1092–1103 (2017).

93. Dong, J. & Ma, Q. TIMP1 promotes multi-walled carbon nanotube-induced lung fibrosis by stimulating fibroblast activation and proliferation. Nanotoxicology 11, 41–51 (2017).

94. Zhang, Y. et al. An NFAT1-C3a-C3aR Positive Feedback Loop in Tumor-Associated Macrophages Promotes a Glioma Stem Cell Malignant Phenotype. Cancer Immunol Res 12, 363–376 (2024).

95. Sehgal, A., Irvine, K.M. & Hume, D.A. Functions of macrophage colony-stimulating factor (CSF1) in development, homeostasis, and tissue repair. Semin Immunol 54, 101509 (2021).

96. Cook, S.A. Understanding interleukin 11 as a disease gene and therapeutic target. Biochem J 480, 1987–2008 (2023).

97. Sedger, L.M. & McDermott, M.F. TNF and TNF-receptors: From mediators of cell death and inflammation to therapeutic giants - past, present and future. Cytokine Growth Factor Rev 25, 453–472 (2014).

98. Cook, S.A. & Schafer, S. Hiding in Plain Sight: Interleukin-11 Emerges as a Master Regulator of Fibrosis, Tissue Integrity, and Stromal Inflammation. Annu Rev Med 71, 263–276 (2020).

99. Widjaja, A.A. et al. Redefining IL11 as a regeneration-limiting hepatotoxin and therapeutic target in acetaminophen-induced liver injury. Sci Transl Med 13 (2021).

100. Widjaja, A.A. et al. Targeting endogenous kidney regeneration using anti-IL11 therapy in acute and chronic models of kidney disease. Nat Commun 13, 7497 (2022).

101. Zhang, B., Yao, K., Zhou, E., Zhang, L. & Cheng, C. Chr20q Amplification Defines a Distinct Molecular Subtype of Microsatellite Stable Colorectal Cancer. Cancer Res 81, 1977–1987 (2021).

102. Dai, M., Pei, X. & Wang, X.J. Accurate and fast cell marker gene identification with COSG. Brief Bioinform 23 (2022).

103. Kotliar, D. et al. Identifying gene expression programs of cell-type identity and cellular activity with single-cell RNA-Seq. Elife 8 (2019).

104. Saunders, A. et al. Molecular Diversity and Specializations among the Cells of the Adult Mouse Brain. Cell 174, 1015–1030 e1016 (2018).

105. Butler, A., Hoffman, P., Smibert, P., Papalexi, E. & Satija, R. Integrating single-cell transcriptomic data across different conditions, technologies, and species. Nat Biotechnol 36, 411–420 (2018).

106. Lee, C.W. et al. Multiplex immunofluorescence staining and image analysis assay for diffuse large B cell lymphoma. J Immunol Methods 478, 112714 (2020).

107. Bankhead, P. et al. QuPath: Open source software for digital pathology image analysis. Sci Rep 7, 16878 (2017).

108. Satoh, J. & Tabunoki, H. A Comprehensive Profile of ChIP-Seq-Based STAT1 Target Genes Suggests the Complexity of STAT1-Mediated Gene Regulatory Mechanisms. Gene Regul Syst Bio 7, 41–56 (2013).

109. Yu, G., Wang, L.G., Han, Y. & He, Q.Y. clusterProfiler: an R package for comparing biological themes among gene clusters. OMICS 16, 284–287 (2012).

110. Mayer, J., Blomberg, J. & Seal, R.L. A revised nomenclature for transcribed human endogenous retroviral loci. Mob DNA 2, 7 (2011).

111. Rooney, M.S., Shukla, S.A., Wu, C.J., Getz, G. & Hacohen, N. Molecular and genetic properties of tumors associated with local immune cytolytic activity. Cell 160, 48–61 (2015).

112. Hanzelmann, S., Castelo, R. & Guinney, J. GSVA: gene set variation analysis for microarray and RNA-seq data. BMC Bioinformatics 14, 7 (2013).

113. Mermel, C.H. et al. GISTIC2.0 facilitates sensitive and confident localization of the targets of focal somatic copy-number alteration in human cancers. Genome Biol 12, R41 (2011).

114. Reich, M. et al. GenePattern 2.0. Nat Genet 38, 500–501 (2006).

115. Zhou, Y. et al. Metascape provides a biologist-oriented resource for the analysis of systems-level datasets. Nat Commun 10, 1523 (2019).

116. Trapnell, C. et al. The dynamics and regulators of cell fate decisions are revealed by pseudotemporal ordering of single cells. Nat Biotechnol 32, 381–386 (2014).

117. Qiu, X. et al. Reversed graph embedding resolves complex single-cell trajectories. Nat Methods 14, 979–982 (2017).

118. Cao, J. et al. The single-cell transcriptional landscape of mammalian organogenesis. Nature 566, 496–502 (2019).

119. McInnes, L., Healy, J., Saul, N., Lukas & Großberger UMAP: Uniform Manifold Approximation and Projection. Journal of Open Source Software 3, 861 (2018).

120. Vento-Tormo, R. et al. Single-cell reconstruction of the early maternal-fetal interface in humans. Nature 563, 347–353 (2018).

121. Efremova, M., Vento-Tormo, M., Teichmann, S.A. & Vento-Tormo, R. CellPhoneDB: inferring cell-cell communication from combined expression of multi-subunit ligand-receptor complexes. Nat Protoc 15, 1484–1506 (2020).

