## Supplementary Figures 1-7 and Tables 1-6 for "Ultra-precision deconvolution of spatial transcriptomics decodes immune heterogeneity and fate-defining programs in tissues"

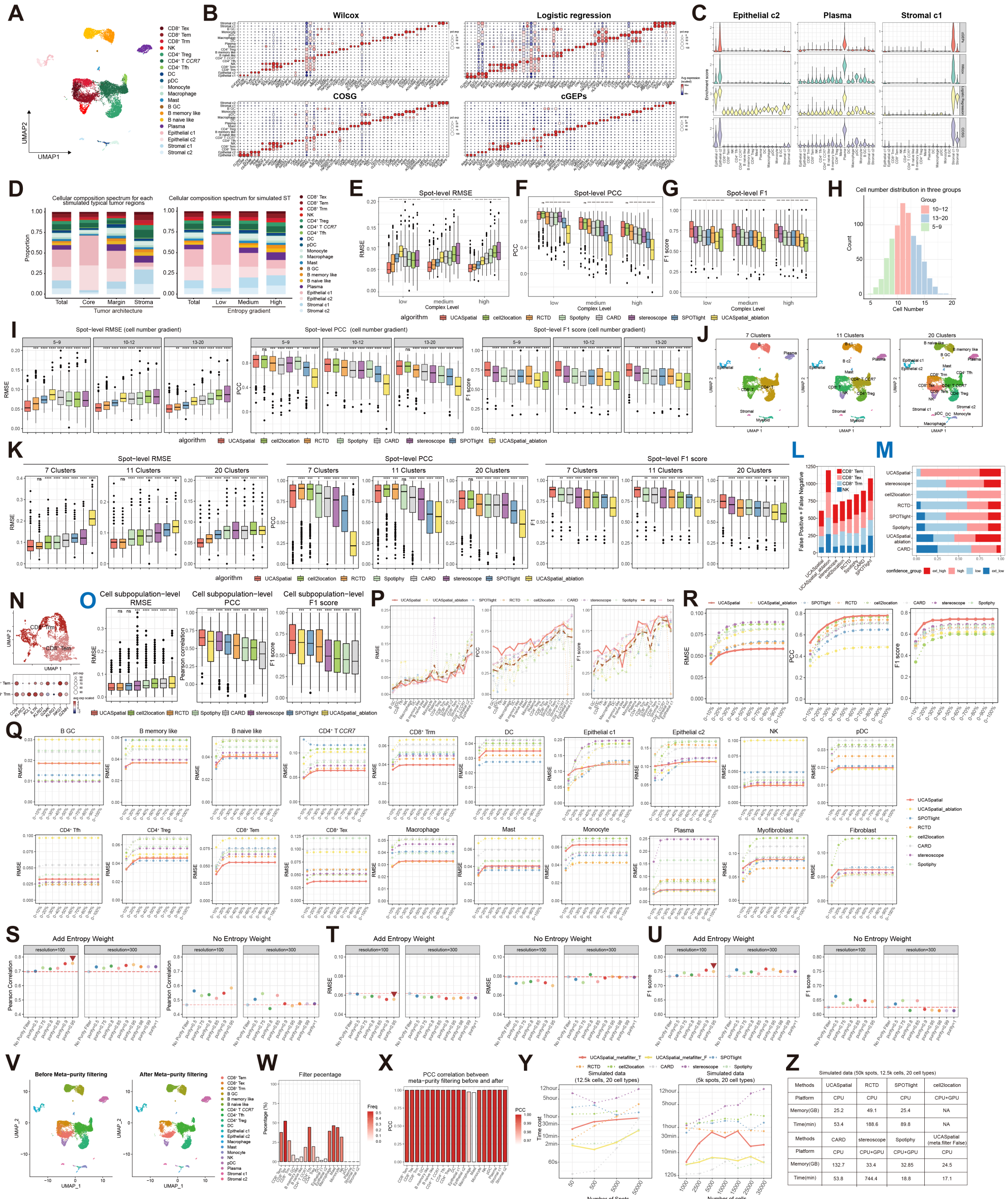


**Fig. S1. Benchmarking the performance of UCASpatial in the simulated ST dataset.** (**A**) UMAP plot displaying the reference scRNA-seq data for simulation comprising 20 cell populations in human CRC. (**B**) Bubble heatmap showing the expression of the top 3 representative genes for each subpopulation identified by different methods (including Wilcoxon-test, Logistic regression, COSG, and cGEPs). Dot size indicates the fraction of cells per subpopulation that express a given marker gene. Colors show the average expression of each gene in each subpopulation. (**C**) Violin plots representing the enrichment scores of the top 10 marker genes identified by cGEPs, Wilcoxon-test, Logistic regression, and COSG for three representative cell types: epithelial c2, plasma cells, and stromal c1. (**D**) Left, stack barplot showing the cellular composition for total spots and spots of three simulated typical tumor regions. Right, stack barplot showing the cellular composition for total spots and spots of three entropy groups. Entropy was calculated based on the cellular composition of each spot. Colors represent different cell subpopulations. (**E**-**G**) Boxplots showing the performances (**E**) RMSE, (**F**) PCC, and (**G**) F1 score of UCASpatial and other methods, including UCASpatial without entropy weighting and meta-purity filtering (UCASpatial_ablation), at spot-level. Colors represent different methods. (**H**) Barplot showing the number of cells each spot contains in three spot-cell-number levels. (**I**) Boxplot showing spot-level RMSE (left), PCC (middle), and F1 score (right) for each method including UCASpatial without entropy weighting and meta-purity filtering (UCASpatial_ablation), across three spot-cell-number levels. (**J**) UMAP plots displaying three different numbers of reference subpopulations (n=7, 11, 20, respectively). (**K**) Boxplot showing spot-level RMSE (left), PCC (middle), and F1 score (right) for each method including UCASpatial without entropy weighting and meta-purity filtering (UCASpatial_ablation), across three reference resolutions. (**L**) Stacked bar plot showing the accumulated count of false predictions of these cell subpopulations for each method, including UCASpatial without entropy weighting and meta-purity filtering (UCASpatial_ablation). Colors represent different cell subpopulations. FP, false positive; FN, false negative; FP + FN, total error frequency. (**M**) The cell-subpopulation-level PCC values for each method were binned into four confidence groups: 0-0.25 (low), 0.25-0.5 (medium-low), 0.5-0.75 (medium-high), and 0.75-1 (high). Stacked bar plot showing the ratio of four confidence groups for each method, including UCASpatial without entropy weighting and meta-purity filtering (UCASpatial_ablation). (**N**) Upper: UMAP plot showing $CD8^+$ Trm and $CD8^+$ Tem cells from scRNA-seq. Bottom: Bubble heatmap showing the expression of representative genes for $CD8^+$ Trm and $CD8^+$ Tem cells. Dot size indicates the fraction of cells per subpopulation that express a given marker gene. Colors showed the average expression of each gene in each subpopulation. (**O**) Boxplot showing the capacity for distinguishing diverse cell subpopulations of UCASpatial, UCASpatial_ablation, and other methods. RMSE (left), PCC (middle), and F1 score (right) between the predicted proportions and the ground truth were calculated for each cell subpopulation. (**P**) Line charts showing cell subpopulation-level RMSE (left), PCC (middle), and F1 score (right) for each method across 20 cell subpopulations. 'avg' represents the average RMSE, PCC or F1 score, and 'best' represents the minimum RMSE, or the maximum PCC or F1 score, respectively, of the six methods excluding UCASpatial and UCASpatial_ablation. (**Q**) Line charts showing the RMSE for each cell subpopulation across different proportions of UCASpatial and other methods, including UCASpatial without entropy weighting and meta-purity filtering (UCASpatial_ablation). (**R**) Line charts representing the average RMSE (left), PCC (middle), and F1 score (right) of 20 cell subpopulations across various proportions for each method. (**S**-**U**) Dot plots displaying the performance (**S**) PCC, (**T**) RMSE, and (**U**) F1 score with different hyperparameter values testing within the simulated ST dataset. Left panel, the performance of UCASpatial with entropy-based weight. Right panel, the performance of UCASpatial without entropy-based weight. Columns in each panel denote performance when varying selected hyperparameters (*resolution* and *purity* of meta-purity filtering process) of UCASpatial. Red arrows denote the default hyperparameter values used in the benchmark part (resolution = 100, purity = 0.95). The red dashed line represents the performance (F1 score, PCC, and RMSE) of UCASpatial without the meta-purity filter. (**V**) UMAP plot displaying 20 cell subpopulations before and after applying meta-purity filtering from the in-silico scRNA-seq reference. (**W**) Bar plots showing the percentage of cells filtered out from each cell subpopulation during the meta-purity filtering process corresponding to the left panel. (**X**) Bar plots showing the PCC correlation between the deconvolution results before and after meta-purity filtering in the in-silico scRNA-seq dataset. (**Y**) Left, line charts showing the time cost of each method under different numbers of spatial spots: 50, 500, 5000, and 50000, using a referenced scRNA-seq with about 12500 cells. Right, Line charts showing the time cost of each method under different numbers of referenced scRNA-seq cells: 1000, 2500, 5000, 10000, 15000, 25000, and 35000, using simulated spatial transcriptomic data with 5000 spots. (**Z**) The runtime and memory usage of different methods for processing a simulated dataset, which contains 50000 spots in spatial transcriptomics data and about 12500 cells in the referenced scRNA-seq data with 20 cell types. (**AA**) Left top, hematoxylin and eosin (H&E) staining slice of colorectal cancer (CRC) sample of 10X Visium HD platform, accompanied by specific pathological annotations including circular muscle, submucosa, mucosa, tumor region, tertiary lymphoid structure (TLS), and small vessels. Red boxes labeled an example TLS. The others, spatial plots displaying the distribution of exampled immune cell subpopulations ($CD8^+$ Tex, $CD8^+$ Tem, $CD4^+$ Tn, $CD4^+$ Tfh, and B), stromal cell subpopulations (*FAP*$^+$ myofib, *CD34*$^+$ iFib, and smooth muscle), and epithelial cell subpopulations (Epithelial c1, Epithelial c2, and Epithelial c3), at spatial locations obtained from UCASpatial in human CRC data based on 10X Visium HD platform with a bin size of 8μm × 8μm. Colors indicate cell subpopulation proportions. (**AB**) Heatmap displaying the cell-subpopulation-level PCC for UCASpatial and RCTD in the 10X Visium HD dataset with a bin size of 8μm × 8μm. (**AC**) Boxplot showing the cell subpopulation-level PCC for UCASpatial and RCTD across 34 cell subpopulations in the 10X Visium HD dataset with a bin size of 8μm × 8μm. The red dots represent the average PCC of each method. (**AD**) Barplot showing the runtime of UCASpatial and RCTD across 34 cell subpopulations in the 10X Visium HD dataset with a bin size of 8μm × 8μm. (**AE**) The graphic model of the mouse hippocampus, with the labels of CA1, CA3, and DG regions. CA, cornu ammonis. DG, dentate gyrus. (**AF-AG**) Spatial plot displaying the distributions of cell subpopulations at spatial locations obtained from UCASpatial in mouse hippocampus data based on Slide-seq V2 platform, including (**AF**) CA1, CA3, and DG (Dentate); (**AG**) ependymal, choroid, and Cajal-Retzius. Colors indicate cell subpopulation proportions. Boxplot elements in Figures (**E-G**, **I**, **K**, **O**, and **AC**) are defined as: center line, median; box limits, upper and lower quartiles; whiskers, 1.5× interquartile range; points, outliers. Statistical analysis in Figures (**E-G**, **I**, **K**, **O**, and **AC**) was performed using a two-tailed paired t-test. ns, non-significant, $p > 0.05$; $*0.01 \le p < 0.05$; $**p < 0.01$; $***p < 0.001$; $****p < 0.0001$.

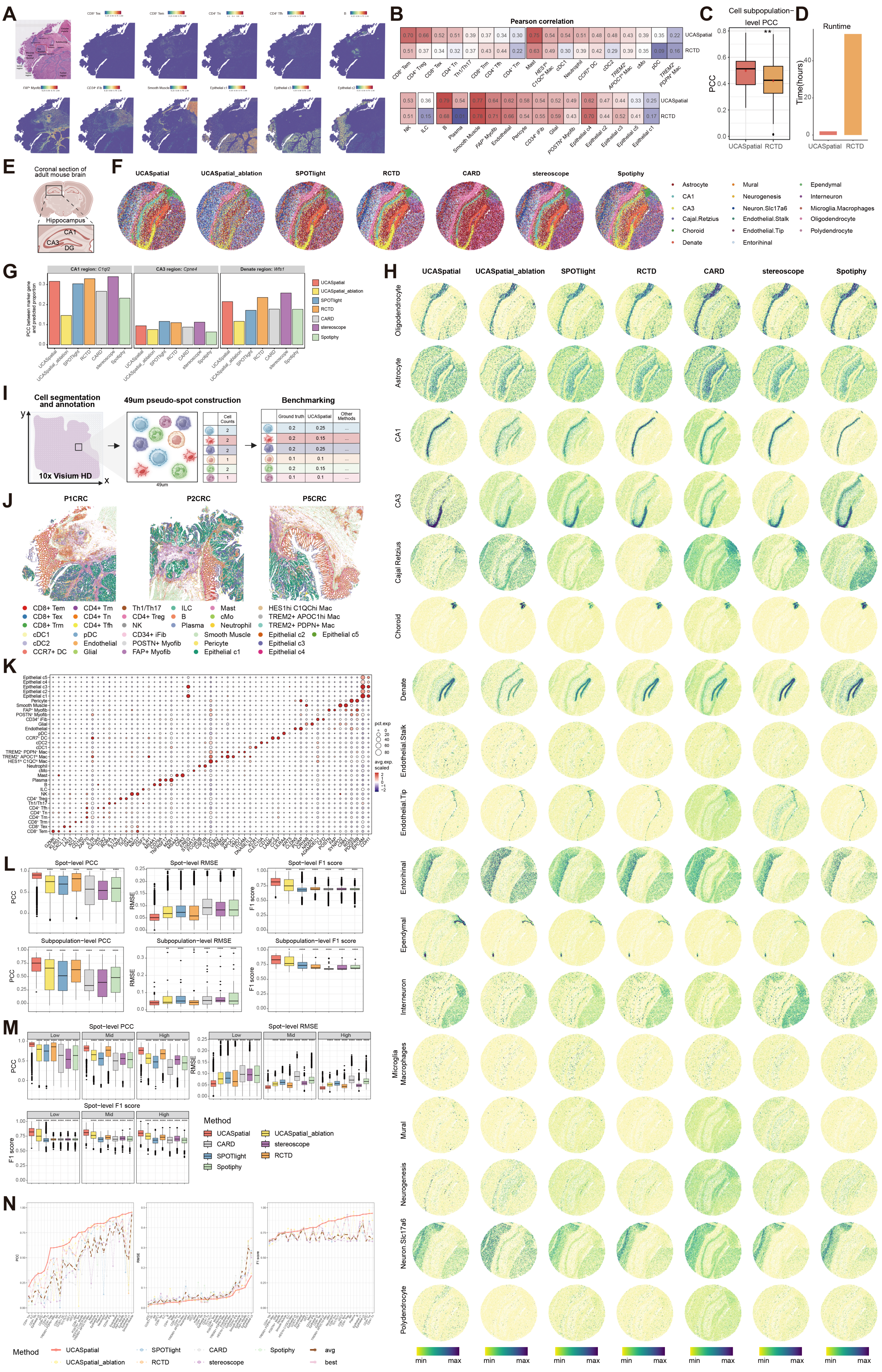

**Fig. S2. Benchmarking the performance of UCASpatial in 10x HD CRC, Slide-seq V2 mouse hippocampus, and Visium HD-derived pseudo-spot datasets.** (**A**) Left top, hematoxylin and eosin (H&E) staining slice of colorectal cancer (CRC) sample of 10X Visium HD platform, accompanied by specific pathological annotations including circular muscle, submucosa, mucosa, tumor region, tertiary lymphoid structure (TLS), and small vessels. Red boxes labeled an example TLS. The others, spatial plots displaying the distribution of exampled immune cell subpopulations ($CD8^+$ Tex, $CD8^+$ Tem, $CD4^+$ Tn, $CD4^+$ Tfh, and B), stromal cell subpopulations (*FAP*$^+$ myofib, *CD34*$^+$ iFib, and smooth muscle), and epithelial cell subpopulations (Epithelial c1, Epithelial c2, and Epithelial c3), at spatial locations obtained from UCASpatial in human CRC data based on 10X Visium HD platform with a bin size of 8μm × 8μm. Colors indicate cell subpopulation proportions. (**B**) Heatmap displaying the cell-subpopulation-level PCC for UCASpatial and RCTD in the 10X Visium HD dataset with a bin size of 8μm × 8μm. (**C**) Boxplot showing the cell subpopulation-level PCC for UCASpatial and RCTD across 34 cell subpopulations in the 10X Visium HD dataset with a bin size of 8μm × 8μm. The red dots represent the average PCC of each method. (**D**) Barplot showing the runtime of UCASpatial and RCTD across 34 cell subpopulations in the 10X Visium HD dataset with a bin size of 8μm × 8μm. (**E**) The graphic model of the mouse hippocampus, with the labels of CA1, CA3, and DG regions. CA, cornu ammonis. DG, dentate gyrus. (**F**) The top 1 cell type in each location inferred from seven different deconvolution methods. Colors represent different cell types. (**G**) Bar plots showing the Pearson correlation coefficient between the marker gene expression and the predicted cell type proportion region of each method. Wfs1, Cpne4, and C1ql2 represent CA1, CA3, and dentate gyrus, respectively. (**H**) Scatter plots showing the detailed deconvolution results of 17 cell types across spatial locations for seven methods. Color was shown to represent the minimal to maximal range of cell type proportions correspondingly. (**I**) Graphic model showing the generation of stimulated spatial transcriptomic data with ground truth. (**J**) Spatial mapping of CRC samples using single-cell-resolution bins, colored by cell subpopulation annotation. (**K**) Bubble heatmap showing the expression of marker genes of 34 cell subpopulations in three Visium HD sections (downsampled 10000 cells for each section). Dot size indicates the fraction of cells per subpopulation that express a given marker gene. Colors show the average expression of each gene in each subpopulation. (**L**) Boxplots showing the performances of PCC, RMSE, and F1 score of UCASpatial and other methods, including UCASpatial without entropy weighting and meta-purity filtering (UCASpatial_ablation), at the spot-level and the cell subpopulation-level. Colors represent different methods. (**M**) Boxplots showing the performances of PCC, RMSE, and F1 score for each method across three complexity levels. (**N**) Line charts showing cell subpopulation-level PCC (left), RMSE (middle), and F1 score (right) for each method across 34 cell subpopulations. 'avg' represents the average RMSE, PCC, or F1 score, and 'best' represents the minimum RMSE, or the maximum PCC or F1 score, respectively, of the five methods excluding UCASpatial and UCASpatial_ablation.

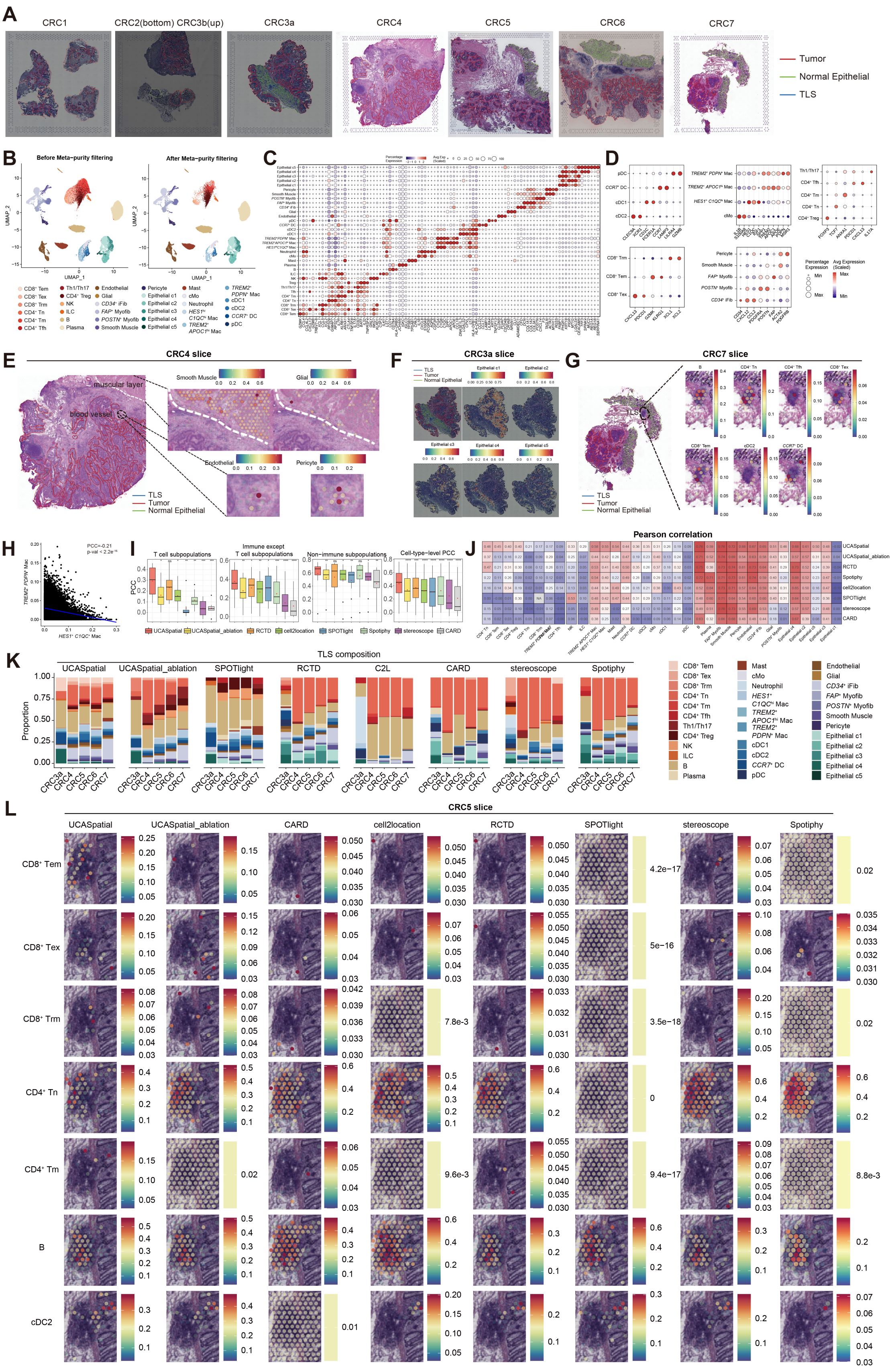

**Fig. S3. UCASpatial accurately annotates cell subpopulations in the human CRC TIMEs.** (A) Hematoxylin and eosin (H&E) staining slices of colorectal cancer (CRC), accompanied by specific pathological annotations. The tumor area is demarcated by a red line, the normal epithelial region is outlined in green, and the Tertiary Lymphoid Structures (TLS) region is indicated by a blue line. (**B**) The UMAP plot showing 34 cell subpopulations before and after applying meta-purity filtering from the scRNA-seq reference of human colorectal cancer. (**C**) Bubble heatmap showing the expression of top 3 marker genes of 34 cell subpopulations in CRC. Dot size indicates the fraction of cells per subpopulation that express a given marker gene. Colors show the average expression of each gene in each subpopulation. (**D**) Bubble heatmap showing the expression of marker genes of indicative cell subpopulations in CRC. Dot size indicates the fraction of cells per subpopulation that express a given marker gene. Colors show the average expression of each gene in each subpopulation. (**E**) Left, H&E staining of CRC4 is presented, accompanied by specific pathological annotations highlighting the exampled blood vessel (dashed black circle) and muscular layer (dashed white circle). Right, Spatial plot displaying the distribution of smooth muscle and glial cells in the muscular layer (upper), and endothelial cells and pericytes in blood vessels (bottom). Colors indicate cell subpopulation proportions. (**F**) Upper left, H&E staining of CRC3a is presented, accompanied by specific pathological annotations highlighting the tumor (red) and healthy epithelial region (green). The others, spatial plots displaying the distribution of epithelial cell subpopulations obtained from UCASpatial. Colors indicate cell subpopulation proportions. (**G**) Left, H&E staining of CRC7 is presented, accompanied by specific pathological annotations highlighting the TLS region (blue). Right, spatial plot displaying the distribution of indicating immune cell subpopulations obtained from UCASpatial. Colors indicate cell subpopulation proportions. (**H**) Scatterplot showing the PCC between the proportion of *HES1*$^{hi}$ *C1QC*$^{hi}$ macrophage and *TREM2*$^{+}$ *PDPN*$^{+}$ macrophage across all spots from 8 CRC slices. (**I**) Boxplot showing cell-subpopulation-level PCC for 8 methods across T cell subpopulations, immune cell subpopulations excluding T cells, non-immune subpopulations, and all cell subpopulations. Statistical analysis was performed using a two-tailed unpaired *t*-test. ns, non-significant, $p > 0.05$; $*0.01 \leq p < 0.05$; $**p < 0.01$; $***p < 0.001$; $****p < 0.0001$. Boxplot elements are defined as: center line, median; box limits, upper and lower quartiles; whiskers, 1.5× interquartile range; points, outliers. (**J**) Heatmap showing cell-subpopulation-level PCC for 8 methods between estimated proportions and marker gene scores. UCASpatial_ablation represents UCASpatial without the entropy-based weight and meta-purity filter. (**K**) Stacked bar plots showing the composition of cell subpopulations in the TLS regions annotated by 8 methods. Only slices containing TLS regions, as defined by pathological annotations shown in (A), are displayed in the stacked bar plots. (**L**) Spatial plot displaying the distribution of cell subpopulations within the TLS of the CRC5 slice, as annotated by 8 methods. UCASpatial_ablation: UCASpatial without entropy-based weight and meta-purity filter.

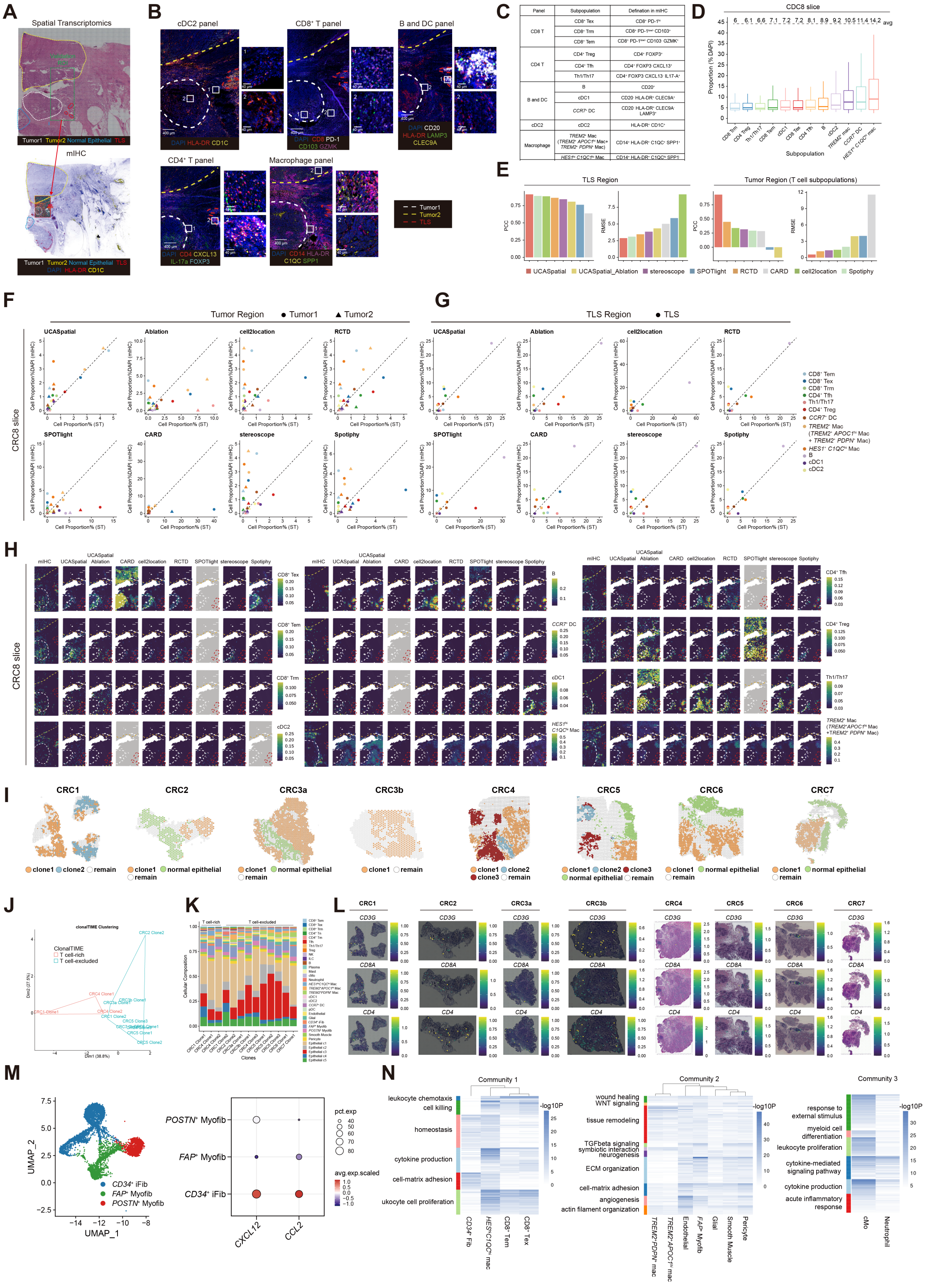


**Fig. S4. Characterization of functional multicellular communities associated with T cell-rich and T cell-excluded TIMEs at a clonal resolution.** (**A**) Pathological annotations of the CRC8 slices within the validation cohort. Upper: The H&E staining of the slice used for spatial transcriptomics sequencing. The dashed green box indicates the validation region of interest (ROI) for mIHC. Dashed lines represent the contours of the regions, as indicated by the text, using the same color. Bottom: The mIHC staining of slices from the same tumor sample, with the validation ROI and pathological annotations highlighted using dashed lines. Regions with conserved tissue morphology across different slices were defined as validation ROIs, and only these validation ROIs were utilized for subsequent statistical analyses. The validation ROIs were divided into three regions: tumor1, tumor2, and TLS, based on pathological annotations. (**B**) Representative immunofluorescence imaging of CRC8 tissues. White, yellow, and red dashed lines indicate tumor1, tumor2, and TLS regions, respectively. The color of each channel is shown at the bottom of each panel. Scale bar, 400 or 40 μm. (**C**) mIHC staining panels designed for cell subpopulation validations in human CRC. The depicted marker combinations were used as surrogates to approximate specific cell states. (**D**) Boxplot showing the proportion of 12 immune cell subpopulations if they existed in pseudo-spots evaluated by mIHC. Numbers above the boxes represent the average proportion of each subpopulation. *TREM2*$^{+}$ mac represents both *TREM2*$^{+}$ *APOC*$^{hi}$ macrophages and *TREM2*$^{+}$ *PDPN*$^{+}$ macrophages. Boxplot elements are defined as follows: center line, median; box limits, upper and lower quartiles; whiskers, 1.5× interquartile range. (**E**) Left, bar plots displaying PCC and RMSE for each method in TLS regions for 12 immune cell subpopulations. Right, bar plots displaying PCC and RMSE for each method in tumor regions for 6 T cell subpopulations. (**F**) Scatter plots showing the proportion of cell subpopulations evaluated by mIHC and each deconvolution method in tumor regions. The dashed line represents y = x. The proportions of cell subpopulations in tumor1 and tumor2 were calculated separately. (**G**) Scatter plots showing the proportion of cell subpopulations evaluated by mIHC and each deconvolution method in TLS regions. The dashed line represents y = x. (**H**) The proportion of cell subpopulations in the mIHC ROI and spatial transcriptomics evaluated by each deconvolution method. The mIHC data was separated into densely 50μm bins, and then the proportion of each subpopulation was quantified (number of subpopulation cells / number of DAPI-identified cells). Completely gray sections represent that the proportion of indicated cell subpopulations in any spot is < 0.03. White, yellow, and red dashed lines indicate tumor1, tumor2, and TLS regions, respectively. (**I**) Spatial plots displaying the distribution of tumor clones and the normal epithelial region at spatial locations obtained from UCASpatial-clonalTIME (Figure 3H). Colors indicate different regions, in which green represents the normal epithelium, and the others represent different tumor clones. (**J**) Scatter plot showing the K-means clustering result of tumor clones based on the proportion of diverse T cell subpopulations. Colors indicate different clonalTIME phenotypes. (**K**) Stacked bar plots showing the subpopulation composition of each tumor clone. Colors indicate different cell subpopulations. The clonalTIME phenotypes are labeled at the top. (**L**) Expression levels of selected T-cell-related genes (*CD3G*, *CD4*, and *CD8A*) in 8 human CRC slices. (**M**) Left, UMAP plot showing fibroblast subpopulations from the scRNA-seq data used in human CRC. Right, bubble heatmap showing the expression of representative genes for indicating cell subpopulations. Dot size indicates the fraction of cells per group that express a given marker gene. Colors showed the average expression of each gene in each subpopulation. (**N**) Heatmap displaying shared pathways within each community. Enriched pathways (hypergeometric test, adj.p <

0.001, adjusted by Benjamini-Hochberg method) in up-regulated genes within each cell subpopulation, shared between at least half of subpopulations within the community (Methods). Colors represent the $-\log_{10}$ p-value calculated by the hypergeometric test. Pathways clustered by shared genes.

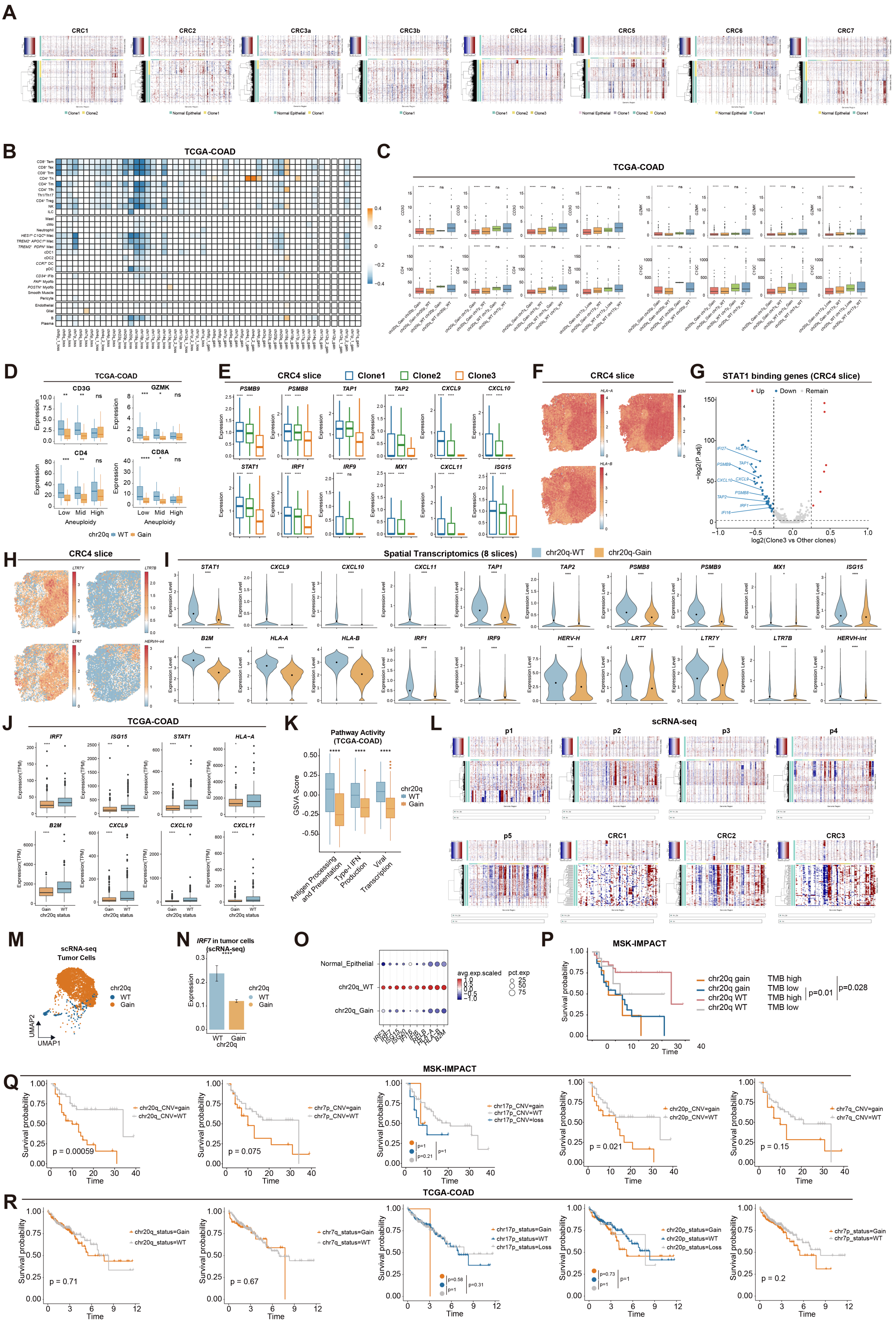


**Fig. S5. Tumor genetic chr20q-gain is associated with the formation of T cell-excluded clonalTIME in CRC.** (**A**) Heatmap showing the CNV profiles inferred from the spatial transcriptomics data of 7 slices: CRC1, CRC2, CRC3a, CRC3b, CRC4, CRC5, CRC6, and CRC7. The CNV profile of the CRC4 slice is shown in (Fig. 4B).

Clonal groupings of spots were determined by hierarchical clustering. (**B**) Heatmap showing the Pearson correlation coefficient (PCC) between the copy number of indicated CNV status and signature scores of each cell subpopulation in the TCGA-COAD dataset. The normalized expression of the top 10 signature genes in each cell subpopulation was used to calculate the average signature score. Colors represent the PCC values. Blank squares indicate non-significant associations (PCC < 0.2). (**C**) Boxplots showing selected gene expression in samples with indicated CNV status in the TCGA-COAD dataset. (**D**) Boxplots showing selected gene expression in samples with indicated aneuploidy and CNV status in the TCGA-COAD dataset. (**E**) Boxplots showing the expression level of selected genes in each clone of the CRC4 slice, colors represent different tumor clones. (**F**) Spatial visualization of the expression levels of selected antigen processing and presenting related genes in the CRC4 slice. (**G**) Volcano plot displaying differentially expressed STAT1-binding genes in clone3 versus other clones in the CRC4 slice. STAT1-binding genes used here were obtained from previous literature (Methods). P-values were calculated using the Wilcoxon rank-sum test and adjusted by the Bonferroni correction method. (**H**) Spatial visualization of the expression levels of each *HERV-H* associated element in the CRC4 slice. (**I**) Violin plots representing the expression of genes associated with interferon response and elements of *HERV-H* between the chr20q-WT, and chr20q-gain clones in 8 slices. (**J**) Boxplots showing the expression of selected interferon-related genes in samples with different CNV statuses in the TCGA-COAD dataset. (**K**) Boxplot displaying the distribution of GSVA activity of the indicated pathways (antigen processing and presentation, type-I IFN production, viral transcription) between chr20q-WT and chr20q-gain tumor samples in the TCGA-COAD dataset. (**L**) Heatmap showing the CNV profiles inferred from 5 public and 3 in-house scRNA-seq data (Methods). (**M**) UMAP plot showing the chr20q status of tumor cells from the 8 scRNA-seq mentioned in (**L**). The chr20q statuses of tumor cells were defined based on the CNV profiles of (**L**). (**N**) Bar plot displaying the average expression of *IRF7* in tumor cells with different chr20q status from scRNA-seq data. Error bars indicate the standard error. (**O**) Bubble heatmap showing the expression of representative interferon-related genes for indicating cell groups. Dot size indicates the fraction of cells per group that express a given marker gene. Colors show the average expression of each gene in each group. (**P**) Overall survival (Kaplan-Meier curves) of CRC patients post ICB in the MSK-IMPACT cohort stratified by the levels of indicating CNV status. (**Q**) Overall survival (Kaplan-Meier curves) of CRC patients without ICB treatment in the TCGA-COAD cohort stratified by the levels of indicating CNV status. (**R**) Overall survival (Kaplan-Meier curves) of CRC patients post-ICB in the MSK-IMPACT cohort stratified by the levels of TMB and chr20q CNV status. Statistical analysis in Figures (**C**, **D**, **E**, **I**, **J**, **K**, and **N**) was performed using a two-tailed unpaired *t*-test. Statistical analysis in Figures (**P**-**R**) was performed using the log-rank test. ns, non-significant, $p > 0.05$; $*0.01 \leq p < 0.05$; $**p < 0.01$; $***p < 0.001$; $****p < 0.0001$. Boxplot elements in Figures (**C**, **D**, **E**, **J**, and **K**) are defined as: center line, median; box limits, upper and lower quartiles; whiskers, 1.5× interquartile range; points, outliers.

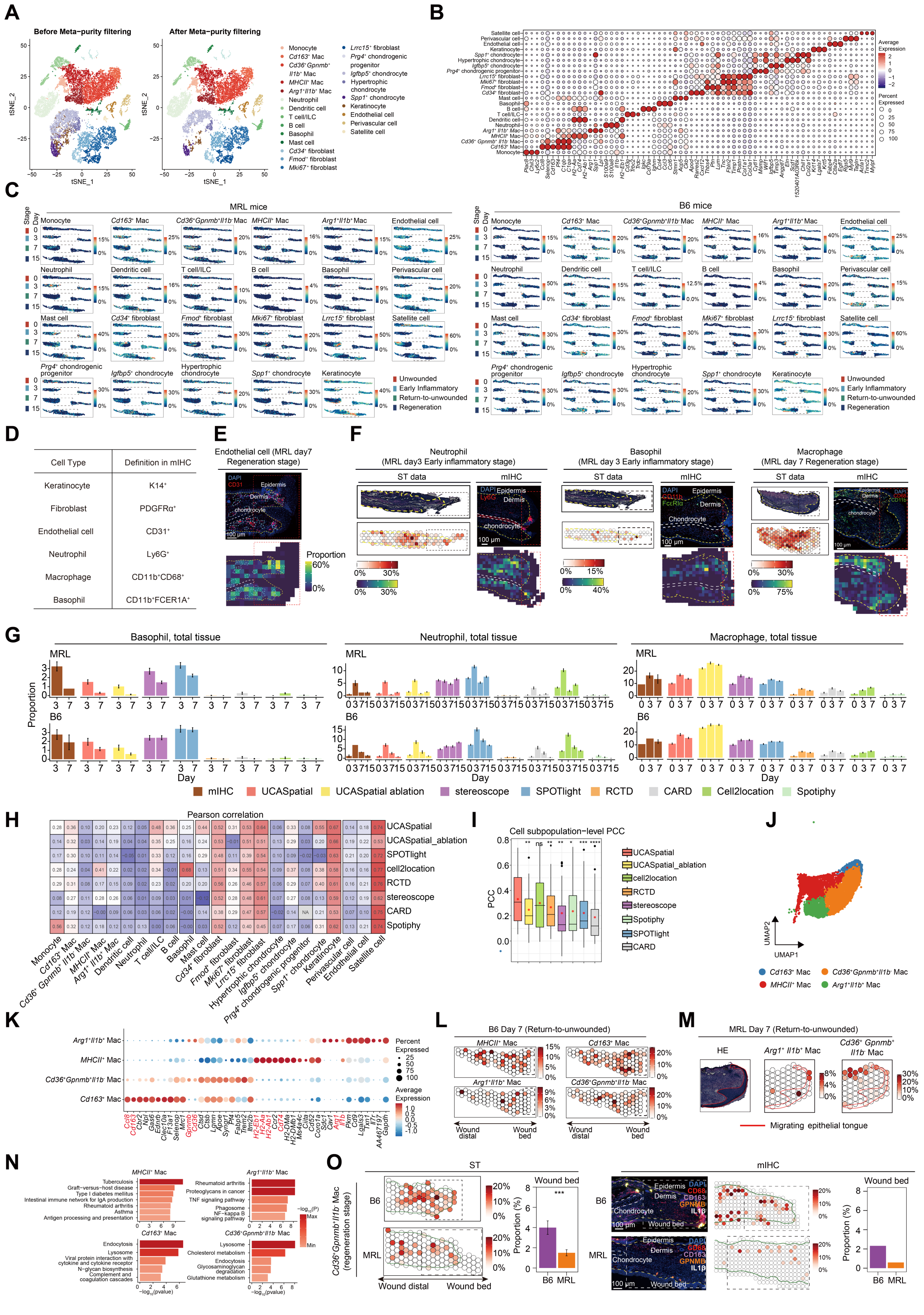

**Fig. S6. UCASpatial accurately annotates cell subpopulations with spatiotemporal dynamics in the murine wound healing ST dataset.** (**A**) The t-distributed Stochastic Neighbor Embedding (tSNE) plot showing 23 cell subpopulations before and after applying meta-purity filtering from the scRNA-seq reference of mouse wound healing. (**B**) Bubble heatmap showing the expression of representative genes for 23 cell subpopulations in mouse ears. Dot size indicates the fraction of cells per subpopulation that express a given marker gene. Colors show the average expression of each gene in each subpopulation. (**C**) Spatial plot displaying the distribution of 23 cell subpopulation proportions at spatial locations obtained from UCASpatial in murine wound healing data. The plot shows MRL mice on the left and B6 mice on the right. Colors indicate cell subpopulation proportions, with a max cutoff of 99.5 quantiles for better visualization. The dashed black rectangles indicate the wound bed. (**D**) The mIHC staining panels for major cell-type validation in murine wound healing. The depicted marker combinations were used as surrogates to approximate specific cell states. (**E**) Upper, a representative immunofluorescence imaging of endothelial cells using ear tissues from MRL mice at the return-to-unwounded stage (day 7 post-injury). Bottom, a quantitative evaluation of the distributions of endothelial cells based on the mIHC, with a 50μm-bin resolution. The staining images indicate DAPI (blue) and CD31 (red). The dashed blue lines indicate the blood vessels. The dashed white rectangles indicate the wound bed. The dashed yellow line indicates the interface of the epidermis-dermis. Scale bar, 100 μm. (**F**) Spatial distributions from ST data and representative immunofluorescence images of neutrophil (left), basophil (middle), and macrophage (right). The dashed rectangles indicate the wound bed. The dashed yellow lines indicate the interface of the epidermis-dermis. The white dashed lines indicate the chondrocyte. Scale bar, 100 μm. (**G**) Left, bar plots showing the proportion of basophils in mIHC and ST data. For mIHC data, B6 day 3, n= 5; MRL day 3, n = 4; B6 day 7, n = 2; MRL day7, n = 1. Middle, bar plots showing the proportion of neutrophils in mIHC and ST data. For mIHC data, B6 day 0 and MRL day 0, n = 3; B6 day 3, n= 1; MRL day 3, n = 2; B6 day 7, n = 2; MRL day7, n = 1; B6 and MRL day 14, n = 3. Right, bar plots showing the proportion of macrophages in mIHC and ST data. For mIHC data, B6 day 0 and MRL day 0, n = 1; B6 day 3, n= 1; MRL day 3, n = 2; B6 day 7, n = 3; MRL day7, n = 2. (**H**) Heatmap displaying cell-subpopulation-level Pearson correlation coefficient (PCC) between gene signature scores and estimated proportions from 8 methods. UCASpatial ablation represents the same framework as UCASpatial but with the core functions of entropy-based weighting and meta-purity filtering process disabled. (**I**) Boxplots showing cell-subpopulation-level PCC between gene signature scores and estimated proportions from 8 methods across 23 cell subpopulations. UCASpatial ablation represents the same framework as UCASpatial but with the core functions of entropy-based weighting and meta-purity filtering process disabled. The red dots represent the average PCC or rho of each method. Boxplot elements are defined as: center line, median; box limits, upper and lower quartiles; whiskers, 1.5× interquartile range; points, outliers. Statistical analysis was performed using a two-tailed unpaired t-test. ns, non-significant, $p > 0.05$; *$0.01 \le p < 0.05$; **$p < 0.01$; ***$p < 0.001$; ****$p < 0.0001$. (**J**) UMAP plot showing the distribution of four macrophage subpopulations. (**K**) Bubble heatmap showing the expression of representative genes for four macrophage subpopulations in murine scRNA-seq data. Dot size indicates the fraction of cells per subpopulation that express a given marker gene. Colors show the average expression of each gene in each subpopulation. (**L**) Spatial visualization of the distribution of indicated macrophage subpopulations at the wound bed on day 7 in B6 mice based on UCASpatial. Colors represent the proportion of each macrophage subpopulation. The dashed rectangles represent the wound bed. (**M**) H&E staining and spatial plot displaying the distribution of *Arg1*$^{+}$ *Il1b*$^{+}$ macrophage and *Cd163*$^{+}$ macrophage proportions in the wound bed at the return-to-unwounded stage of MRL mice. The red lines represent the migrating epithelial tongue. (**N**) Functional enrichment analysis using GO terms showing the functions of 4 macrophage subpopulations. Markers of macrophage subpopulations were used (cutoff: $\log_2$ [fold change] > 0.25, min.pct = 0.1, and min.diff.pct = 0.1). (**O**) The left panel, spatial distribution of *Cd36*$^{+}$ *Gpnmb*$^{+}$ *Il1b*$^{-}$ macrophage at the regeneration stage (day 15 post-injury) in B6 (upper) and MRL mice (bottom), and its proportion at the wound bed based on UCASpatial's deconvolution results. The green line indicates the interface of the epidermis-dermis. The dashed rectangle indicates the wound bed. Statistical analysis was performed using a two-tailed unpaired t-test. ***$p < 0.001$. The right panel, a representative immunofluorescence imaging and a quantitative evaluation of the distribution (50μm-bin resolution) of *Cd36*$^{+}$ *Gpnmb*$^{+}$ *Il1b*$^{-}$ macrophage at the regeneration stage (day 15 post-injury) in B6 (upper) and MRL mice (bottom). Bar plot showing its proportion at the wound bed based on mIHC. The staining images indicate DAPI (blue), CD68 (red), GPNMB (orange), and IL-1β (white). The dashed yellow line and the green line indicate the interface of the epidermis-dermis. The dashed white line indicates the chondrocyte. The dashed rectangles indicate the wound bed. Scale bar, 100 μm.

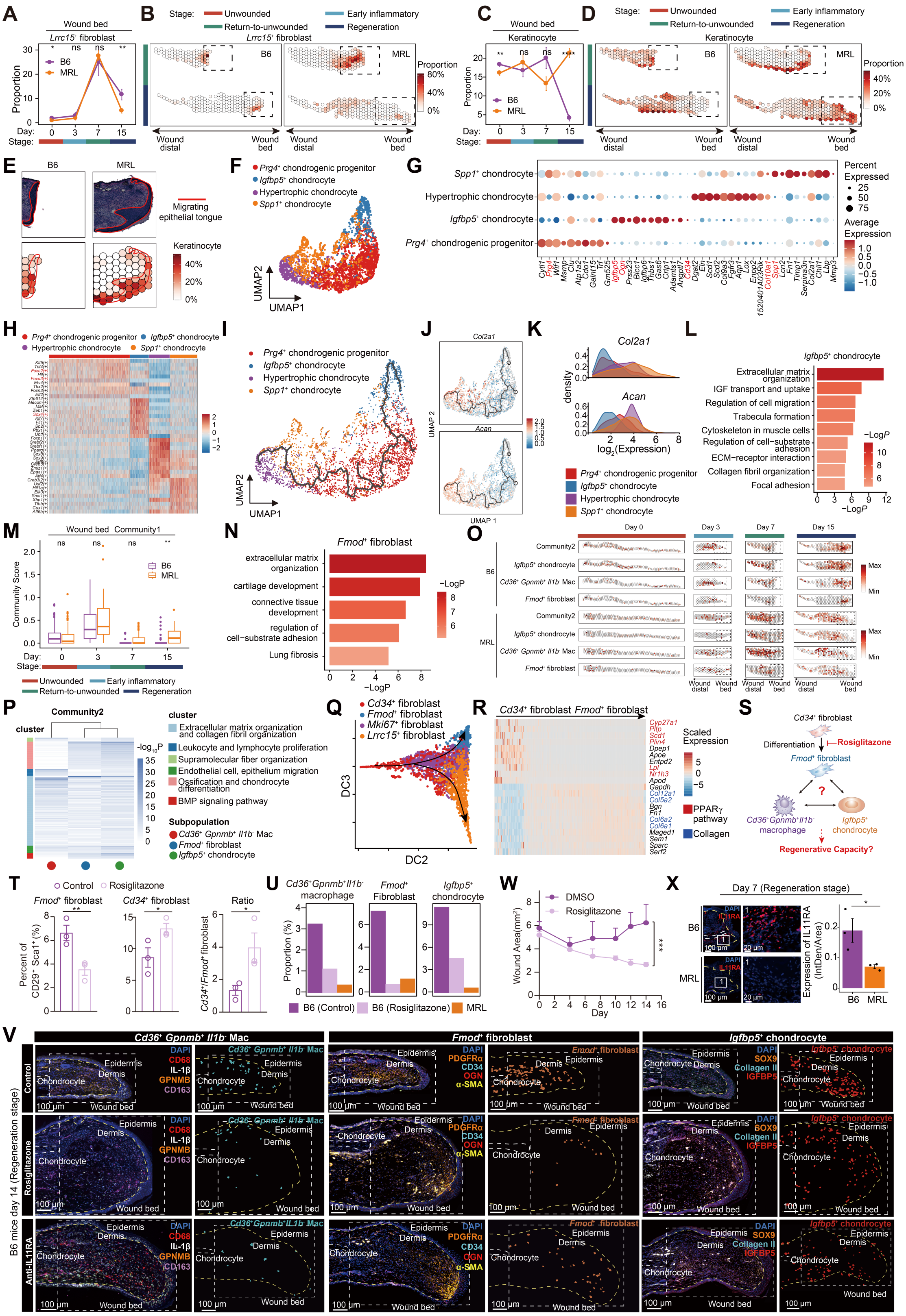


**Fig. S7. UCASpatial reconstructs the spatiotemporal dynamics of diverse macrophage subpopulations during murine wound healing.** (**A**) Lineplot showing the temporal dynamics of the proportion of *Lrrc15*$^{+}$ fibroblasts at the wound bed between MRL and B6 mice. (**B**) Spatial visualization of the distribution of *Lrrc15*$^{+}$ fibroblasts at the wound bed on day 7 and day 15 in MRL and B6 mice. Colors represent the proportion. The dashed rectangles represent the wound bed. (**C**) Lineplot showing the temporal dynamics of the proportion of keratinocytes at the wound bed between MRL and B6 mice. (**D**) Spatial visualization of the distribution of keratinocytes at the wound bed on day 7 and day 15 in MRL and B6 mice. Colors represent the proportion. The dashed rectangles represent the wound bed. (**E**) H&E staining and spatial plot displaying the distribution of keratinocyte proportions in the wound bed at the return-to-unwounded stage of B6 and MRL mice. The red lines represent the migrating epithelial tongue. (**F**) UMAP plot showing four chondrocyte subpopulations in scRNA-seq. (**G**) Bubble heatmap showing the expression of representative genes for four chondrocyte subpopulations in murine scRNA-seq data. Dot size indicates the fraction of cells per subpopulation that express a given marker gene. Colors show the average expression of each gene in each subpopulation. (**H**) Heatmap showing the expression of the top 10 transcription factor regulons for four chondrocyte subpopulations. (**I**) UMAP plot showing the trajectory of four chondrocyte subpopulations inferred by Monocle3. (**J**) UMAP plots showing the expression of *Col2a1* and *Acan* across four chondrocyte subpopulations along the trajectory. (**K**) Density plots showing the expression level of *Col2a1* and *Acan* across four chondrocyte subpopulations. **(L)** Functional enrichment analysis showing the functions of *Igfbp5*$^{+}$ chondrocyte using its signature genes (cutoff: $\log_2$ [fold change] > 0.1, min.pct = 0.1, and min.diff.pct = 0.25). (**M**) Boxplots showing the score of community1 at wound bed in MRL and B6 mice at different stages. Statistical analysis was performed using a two-tailed unpaired *t*-test. ns, non-significant, $^{**}p < 0.01$. Boxplot elements are defined as: center line, median; box limits, upper and lower quartiles; whiskers, 1.5× interquartile range; points, outliers. (**N**) Functional enrichment analysis using GO terms showing the functions of *Fmod*$^{+}$ fibroblast using its signature genes (cutoff: $\log_2$ [fold change] > 0.25, min.pct = 0.25, and min.diff.pct = 0.25). (**O**) Spatial visualization of the distribution of the pro-fibrotic community, including *Igfbp5*$^{+}$ chondrocyte, *Fmod*$^{+}$ fibroblast, and *Cd36*$^{+}$*Gpnmb*$^{+}$*Il1b*$^{-}$ macrophage subpopulations at the wound bed on day 0, day 3, day 7, and day 15 in B6 mice (upper) and MRL mice (down) based on UCASpatial. Colors represent the proportion of each cell subpopulation. The dashed rectangles represent the wound bed. (**P**) Heatmap displaying shared functional pathways within community2. Enriched pathways (hypergeometric test, adj.p < 0.001, adjusted by Benjamini-Hochberg method) in up-regulated genes within each cell subpopulation, shared between at least half of the subpopulations within the community (Methods). Colors represent the -log10 p-value calculated by the hypergeometric test. Pathways clustered by shared genes. (**Q**) Inferred differentiation trajectory among four fibroblast subpopulations from scRNA-seq by diffusion map algorithm. (**R**) Heatmap showing the transcriptional trends of selective genes along the diffusion components during the transition from *Cd34*$^{+}$ fibroblasts to *Fmod*$^{+}$ fibroblasts. The color scale represents Z-score-normalized expression levels. (**S**) The schematic diagram for intervening in the differentiation of *Fmod*$^{+}$ fibroblasts. (**T**) B6 mice were subjected to a 2.6 mm diameter ear punch injury followed by daily treatment with petrolatum containing either DMSO (control, n = 3) or rosiglitazone (n = 3). Flow cytometry analysis of *Fmod*$^{+}$ fibroblasts (left) (defined as $CD34^{-}CD55^{+}CD90.2^{+}$) and *Cd34*$^{+}$ fibroblasts (middle) (defined as $CD34^{+}CD55^{-}$

CD90.2$^{+}$) on day 7 post-injury. Right, the ratio of *Cd34*$^{+}$ fibroblasts and *Fmod*$^{+}$ fibroblasts. (**U**) Representative immunofluorescence imaging of cell subpopulations within community2 using ear tissues in control (upper), rosiglitazone treatment (middle), and anti-IL11RA treatment (bottom) group in B6 mice at the regeneration stage (day 14 post-injury), including *Cd36*$^{+}$ *Gpnmb*$^{+}$ *Il1b*$^{-}$ macrophages (CD68$^{+}$GPNMB$^{+}$IL1β$^{-}$CD163$^{-}$) (left), *Fmod*$^{+}$ fibroblasts (PDGFRα$^{+}$OGN$^{+}$CD34$^{-}$α-SMA$^{-}$) (middle), and *Igfbp5*$^{+}$ chondrocytes (SOX9$^{+}$IGFBP5$^{+}$Collagen II$^{-}$) (right). The control group was treated with DMSO. The dashed white lines indicate chondrocytes. The dashed white rectangle indicates the wound bed. The dashed yellow line indicates the interface of the epidermis-dermis. Scale bar, 100 μm. (**V**) Bar plots showing the proportion of *Cd36*$^{+}$ *Gpnmb*$^{+}$ *Il1b*$^{-}$ macrophages (left), *Fmod*$^{+}$ fibroblasts (middle), and *Igfbp5*$^{+}$ chondrocytes (right) in wound bed regions at the regeneration stage in mIHC data. The B6 control group was administered with DMSO. (**W**) The wound curves of B6 ear wounds in the control (n = 6) and the treatment group (n = 6) demonstrate wound healing over time. The control group was treated with petrolatum containing DMSO, and the treatment group was treated with petrolatum containing rosiglitazone. Wound area was recorded and calculated as $\pi \times r1 \times r2$, $r1$ and $r2$ represent radius aligned with or orthogonal to the body axis, respectively.. (**X**) Left, representative immunofluorescence imaging of ear tissues in B6 and MRL mice at the return-to-unwounded stage (day 7 post-injury). The staining images indicate DAPI (blue) and IL11RA (red). The dashed white lines indicate chondrocytes. The dashed yellow line indicates the interface of epidermis-dermis. Scale bar, 100 μm. Right, bar plot showing the expression of IL11RA in B6 (n = 3) and MRL mice (n = 3). IntDen/Area was used to quantify the expression of IL11RA, calculated by dividing the integrated density (IntDen) by the area of the tissue region. Statistical analysis in Figures (**A, C, M, and W**) was performed using a two-tailed unpaired t-test. Statistical analysis in Figures **(T and X)** was performed using a one-tailed unpaired t-test. *$0.01 \le p < 0.05$; **$p < 0.01$; ***$p < 0.001$.

### Supplementary Tables

**Table S1. Cellular composition of simulated spots reflecting typical tumor architectures within the TME, related to Figures 2 and S1.**

| **Cell types** | **Relative proportion in Tumor core (TC)** | **Relative proportion in Tumor margin (TM)** | **Relative proportion in Tumor stroma (TS)** |
|---|---|---|---|
| Epithelial | 50% - 80% | 30% - 60% | 0% - 20% |
| Myeloid | 0% - 20% | 10% - 30% | 10% - 20% |
| Stromal | 0% - 10% | 10% - 30% | 20% - 50% |
| CD4 T | 0% - 20% | 0% - 20% | 10% - 30% |
| CD8 T | 0% - 20% | 0% - 20% | 10% - 30% |
| B | 0 | 0% - 20% | 0% - 20% |
| Plasma | 0 | 0% - 20% | 0% - 20% |
| Total | 50% - 150% | 50% - 200% | 50% - 190% |

Detailed proportions for cell subpopulations within each cell type were referred from scRNA-seq data.

**Table S2. Clinical characteristics and sample information for patients of human CRC, related to Figures 3, 4, and S2-S4.**

| sample ID | Gender | Age | Tumor Location | Histologic type of tumor | Pathologic Staging (pTNM) |
|---|---|---|---|---|---|
| CRC1 | Female | 57 | Right Colon | Adenocarcinoma | pT3(tumor invades into pericolic tissues),pN0(no regional lymph node metastasis),M0(No distant metastasis) |
| CRC2 | Female | 52 | Left Colon | Adenocarcinoma | pT4b(tumor directly invades or is adherent to other),N2b(≥7 regional lymph node metastases),M0(no distant metastasis) |
| CRC3a/ CRC3b | Male | 67 | Left Colon | Adenocarcinoma | pT3(tumor invades into pericolic tissues),N1a(one regional lymph node metastasis),pM(distant Metastasis) |
| CRC8 | Male | 67 | Right Colon | Adenocarcinoma | cT4(tumor directly invades or is adherent to other),N2(4 or more regional nodes metastases),M0(no distant metastasis) |
| CRC9 | Female | 71 | Right Colon | Adenocarcinoma | pT3(tumor invades into pericolic tissues)N0(no regional lymph node metastasis), M0(no distant metastasis) |
| CRC10 | Male | 51 | Right Colon | Adenocarcinoma | pT3(tumor invades into pericolic tissues)N2(4 or more regional nodes metastases), cM(liver metastasis) |

**Table S3. Cell subpopulations and their abbreviations in the human CRC dataset, related to Figures 3, 4, and S2-S4.**

| Cell subpopulation | Abbreviation |
|---|---|
| $CD8^+$ effector memory T cell | $CD8^+$ Tem |
| $CD8^+$ exhausted T cell | $CD8^+$ Tex |
| $CD8^+$ resident memory T cell | $CD8^+$ Trm |
| $CD4^+$ naive T cell | $CD4^+$ Tn |
| $CD4^+$ memory T cell | $CD4^+$ Tm |
| $CD4^+$ T follicular helper cell | $CD4^+$ Tfh |
| T helper 1/17 cell | Th1/Th17 |
| $CD4^+$ regulatory T cell | $CD4^+$ Treg |
| Natural killer cell | NK |
| Innate lymphoid cell | ILC |
| B cell | B |
| Plasma cell | Plasma |
| Mast cell | Mast |
| Classical Monocyte | cMo |
| Neutrophil | Neutrophil |
| $HES1^{hi}$ $C1QC^{hi}$ macrophage | $HES1^{hi}$ $C1QC^{hi}$ Mac |
| $TREM2^+$ $APOC1^{hi}$ macrophage | $TREM2^+$ $APOC1^{hi}$ Mac |
| $TREM2^+$ $PDPN^+$ macrophage | $TREM2^+$ $PDPN^+$ Mac |
| Type 1 conventional dendritic cell | cDC1 |
| Type 2 conventional dendritic cell | cDC2 |
| $CCR7^+$ dendritic cell | $CCR7^+$ DC |
| Plasmacytoid dendritic cell | pDC |
| $CD34^+$ inflammatory fibroblast | $CD34^+$ iFib |
| $FAP^+$ myofibroblast | $FAP^+$ Myofib |
| $POSTN^+$ myofibroblast | $POSTN^+$ Myofib |
| Smooth muscle cell | Smooth Muscle |
| Pericyte | Pericyte |
| Endothelial cell | Endothelial |
| Glial cell | Glial |
| Epithelial cell type 1 | Epithelial c1 |
| Epithelial cell type 2 | Epithelial c2 |
| Epithelial cell type 3 | Epithelial c3 |
| Epithelial cell type 4 | Epithelial c4 |
| Epithelial cell type 5 | Epithelial c5 |

**Table S4. Gene list for calculating ModuleScore in CRC, related to Figure S2.**

| Cell subpopulation | Gene |
|---|---|
| $CD8^+$ Tem | *CTSW, CST7, XCL2, TRAT1, PRF1, TRGC2, DTHD1, CCL5, GZMM, SH2D1A, GZMA, NKG7, CD8A, CD8B, CRTAM, GZMH, KLRG1, IFNG, SAMD3, GZMK* |
| $CD8^+$ Tex | *NKG7, KLRC2, SLA2, GZMA, CXCR6, VCAM1, TTN, LAG3, CRTAM, IFNG, GZMH, CTSW, CD8B, CD8A, KLRC4, LRRN3, KRT86, NELL2, CXCL13, TRBV7-6* |
| $CD8^+$ Trm | *GZMM, NKG7, GPR171, CCL5, HOPX, IFNG, CRTAM, GPR18, KLRK1, SLA2, LINC01871, CD8A, XCL2, KLRD1, XCL1, CD8B, KLRC2, TRGC2, LINC02446, CD160* |
| $CD4^+$ Tn | *CD2, TNFSF8, GIMAP4, OXNAD1, AP3M2, CD3D, CD3G, BCL11B, CCR7, IPCEF1, CHRM3-AS2, GIMAP7, TRAC, TRAT1, LINC01550, TCF7, LEF1, TXK, CD40LG, PASK* |
| $CD4^+$ Tm | *RORA, IL7R, TRAC, GIMAP7, CD6, CD3G, ICOS, CD3E, CD2, CD3D, CD28, LINC01871, ITK, SPOCK2, GPR171, KLRB1, CCR6, TRAT1, CD40LG, MYBL1* |
| $CD4^+$ Tfh | *SH2D1A, NMB, ZNF831, MAGEH1, CCR4, CD40LG, CH25H, TIGIT, PDCD1, TOX2, SEPTIN9, PLAAT4, CTLA4, PTPN13, SEPTIN6, CD28, CXCL13, TNFSF8, FAAH2, FBLN7* |
| Th1/Th17 | *IKZF3, SPOCK2, CAMK4, ICOS, CD3G, LCK, RASGRP1, GZMA, CD5, GPR171, ADAM19, TRAT1, TMIGD2, LINC01871, TNFRSF25, CCR6, CD6, CXCR6, CD40LG, IL17A* |
| $CD4^+$ Treg | *ZC3H12D, TBC1D4, CRADD, CD27, CD28, TNFRSF9, BATF, TNFRSF18, TNFRSF4, LAYN, CTLA4, TIGIT, SLAMF1, IL2RA, IKZF2, LINC01943, LAIR2, RTKN2, FOXP3, CCR8* |
| NK | *IL2RB, CLIC3, TBX21, CTSW, NKG7, XCL1, XCL2, KLRC2, GZMH, MATK, SAMD3, KLRD1, PRF1, S1PR5, KLRC1, SH2D1B, GNLY, FGFBP2, TRDC, KLRF1* |
| ILC | *XCL2, ALDOC, TEC, IL4I1, TXK, XCL1, SPINK2, TNFRSF25, IL23R, KRT86, PCDH9, TMIGD2, KIT, SLC4A10, LINC00299, KRT81, C20orf204, MPV17L, TNFSF11, CSF2* |
| Mast | *VWA5A, RGS13, MAOB, GATA2, IL1RL1, AL662860.1, TPSB2, RHEX, HPGDS, TPSD1, SLC18A2, HDC, TPSAB1, CPA3, AL157895.1, ADCYAP1, CTSG, MS4A2, GCSAML, CMA1* |
| cMo | *OLR1, IL1A, NLRP3, IL1RN, AC005280.2, S100A8, FCAR, GPR84, FPR2, RETN, DNAAF1, LUCAT1, MCEMP1, AC245128.3, MIR3945HG, CD300E, FCN1, APOBEC3A, S100A12, SERPINB2* |
| Neutrophil | *S100A8, SMIM25, FPR1, TNFAIP6, CLEC4E, AQP9, IL1RAP, MCEMP1, AC245128.3, CSF3R, FPR2, HCAR2, PI3, HCAR3, AL034397.3, ADGRG3, FFAR2, CXCR2, CMTM2, FCGR3B* |

| | |
|---|---|
| *HES1*$^{hi}$ *C1QC*$^{hi}$ Mac | *RGL1, SLC40A1, P2RY13, CMKLR1, SIGLEC1, MERTK, IGF1, LILRB5, GAL3ST4, MMP12, STAB1, GPR34, GATM, LINC00996, SIGLEC12, FOLR2, F13A1, CD209, ADORA3, CD163L1* |
| *TREM2*$^{+}$ *APOC1*$^{hi}$ Mac | *FBP1, MMP12, LPL, CYP27A1, BHLHE41, OSCAR, LINC02345, MARCO, CCL18, NR1H3, TREM2, RETN, DNAJC5B, DCSTAMP, OTOA, HAMP, MMP2-AS1, SLC2A5, CHI3L1, APOC2* |
| *TREM2*$^{+}$ *PDPN*$^{+}$ Mac | *VMO1, MERTK, CD209, OLFML2B, ITGAM, CCL13, OSCAR, CMKLR1, FCGR1A, PLA2G15, MSR1, TMEM37, TREM2, FAM20A, LINC01094, CCL18, MARCO, FOLR2, FCGR1B, LILRB5* |
| cDC1 | *DNASE1L3, HDAC9, BATF3, PPM1J, IDO1, FLT3, ASB2, ENPP1, SERPINF2, HLA-DOB, ZNF366, TLR10, PLCD1, RAB7B, LINC02206, WDFY4, CLNK, SLC24A4, CLEC9A, XCR1* |
| cDC2 | *PLD4, P2RY13, CLEC4A, PKIB, DNASE1L3, PID1, CALHM6, IL1R2, FCGR2B, GAPT, AL138899.1, CFP, PSTPIP2, CEACAM4, RGS18, CD1D, CD1E, FCER1A, CLEC10A, CD1C* |
| *CCR7*$^{+}$ DC | *UBD, GPR157, ARHGAP22, TNNT2, TBC1D8, ANKRD33B, IDO2, INSM1, TREML1, CRLF2, CCL19, BX255923.2, EBI3, HMSD, SLCO5A1, CCL17, TVP23A, LAMP3, CCL22, NCCRP1* |
| pDC | *FUZ, CXCR3, ENTPD7, PLAAT3, HIST2H2AA4, PLD4, SCT, ZFAT, TNFSF13, NIBAN3, PTPRS, MYBL2, IGLON5, PTCRA, ANKRD53, DRD4, CLEC4C, KRT5, LILRA4, SLC32A1* |
| *CD34*$^{+}$ iFib | *CCL11, FABP4, ABCA8, CILP, CCL8, DPT, CLEC3B, SVEP1, SFRP1, ADH1B, SCARA5, C16orf89, ABCA6, TNXB, HAPLN1, OGN, ABCA10, C7, SFTA1P, CP* |
| *FAP*$^{+}$ Myofib | *COL10A1, HTRA3, ITGA11, CXCL6, LSAMP, GREM1, MMP1, MFAP2, PODNL1, TWIST1, CTHRC1, COL7A1, CPXM1, FAP, SFRP4, THBS2, WISP1, COL11A1, MMP3, WNT2* |
| *POSTN*$^{+}$ Myofib | *CNTFR, C1QL1, SOX6, HSD17B2, BMP5, DHRS7C, PTGDR2, NSG1, TRPA1, TSLP, PCSK6, ALKAL2, SCUBE2, COL4A5, SCUBE1, NPY, ENHO, COL4A6, GLP2R, VSTM2A* |
| Smooth Muscle | *BCHE, LMOD1, PCDH7, DIO2, SYT10, HHIP-AS1, MYH11, CNN1, CES1, ACTG2, NPNT, MFAP5, SEMA3D, HHIP, DES, KCNJ3, OSR1, MYOCD, SOSTDC1, HSD17B6* |
| Pericyte | *EFHD1, FAM162B, SLC7A2, CDH6, GJA4, MAP3K7CL, AVPR1A, KCNJ8, GJC1, NOTCH3, ABCC9, HEYL, ITGA7, FHL5, ENPEP, FOXS1, LINGO1, HIGD1B, COX4I2, RERGL* |
| Endothelium | *KANK3, MADCAM1, VWF, CXorf36, ACKR1, KDR, CCM2L, CDH5, CLDN5, SHANK3, APLNR, BCL6B, PCAT19, TM4SF18, EXOC3L2, ROBO4, ECSCR, SOX17, MYCT1, FAM110D* |
| Glial | *MPZ, SORCS1, S100A1, CMTM5, GFRA3, L1CAM, TTR, PLP1, COL28A1, FOXD3, CADM2, XKR4, PCSK2, SOX2, NRXN1, CDH19, MYOT, PTPRZ1, FOXD3-AS1, SOX10* |

| | |
|---|---|
| B | *ARHGAP24, BACH2, LINC01857, CD79B, CD22, P2RX5, CXCR5, LINC02397, LY9, FCRLA, TNFRSF13B, CD79A, LINC00926, LINC01781, BANK1, TNFRSF13C, BLK, VPREB3, CD19, MS4A1* |
| Plasma | *FAM30A, ABCB9, MEI1, CD79A, LINC01480, C16orf74, AMPD1, DERL3, TXNDC5, SPINK2, SPAG4, MZB1, POU2AF1, CCR10, LINC02362, FCRL5, IGLL5, TNFRSF17, IGLV3-1, IGLV6-57* |
| Epithelium c1 | *CKB, C19orf33, KRT20, GPX2, FXYD3, KRT19, AC103702.2, ASCL2, PIGR, KRT8, KRT18, PRAC1, EPCAM, LGALS4, C10orf99, PHGR1, PRSS3, CD24, FABP1, DPEP1* |
| Epithelium c2 | *RHOV, TMEM238L, SEMA4G, SULT1B1, CES3, HOXB13, MT1H, RHBDL2, HLA-G, CFTR, OLFM4, CA1, RNF186, C4orf19, SLC28A2, GSTA1, UGT2A3, LEFTY1, CHP2, ADH1C* |
| Epithelium c3 | *HOXB9, NOX1, CLDN1, ETV4, AZGP1, TRIM31, RNF43, MET, PARD6B, SLC26A3, FABP6, CEACAM7, AXIN2, PPP1R14C, GAL, PRAP1, DPEP1, LY6G6F-LY6G6D, CEL, UCA1* |
| Epithelium c4 | *MB, CLCA1, FABP2, SPDEF, NR3C2, TPSG1, AC005833.1, L1TD1, ATOH1, SCGB2A1, ANXA13, GALNT8, RAP1GAP, REP15, NEURL1, ANO7, RETNLB, BEST2, ITLN1, ZG16* |
| Epithelium c5 | *RBP4, CCL26, SDR16C5, DMBT1, NPW, KLK15, CST6, KLK11, PLEKHS1, FAIM2, C4BPB, FAM3B, KLK12, TMPRSS3, TFF2, PRSS2, KREMEN2, TCN1, CLDN18, KLK8* |

| **Table S5. CNV annotations in human colorectal cancer, related to Figures 3, 4, S3, and S4.** | |
|---|---|
| **CNV** | **Location** |
| chr1q | chr1:180972712-212361853 |
| chr1p_1 | chr1:1311585-9198906 |
| chr1p_2 | chr1:15571699-25832942 |
| chr12q_1 | chr12:49184795-57959148 |
| chr12p | chr12:6650301-16037381 |
| chr12q_2 | chr12:109477402-132829078 |
| chr13q | chr13:19633659-114305817 |
| chr14q | chr14:96501433-105856218 |
| chr15q | chr15:34341719-59640239 |
| chr17q | chr17:36591879-42388568 |
| chr17p | chr17:508668-8156506 |
| chr18p | chr18:158383-18500000 |
| chr18q | chr18:18500001-80050651 |
| chr20q | chr20:28100001-64242253 |
| chr20p | chr20:270863-28100000 |
| chr22q | chr22:17137511-42070842 |
| chr3q | chr3:195274745-197888436 |
| chr4p | chr4:42408373-50000000 |
| chr4q_2 | chr4:118722823-186726722 |
| chr4q_1 | chr4:50000001-87459455 |
| chr5p | chr5:196868-36184319 |
| chr7q | chr7:60100001-158704804 |
| chr9q | chr9:128947699-137618906 |
| chr7p | chr7:192571-60100000 |
| chr8p | chr8:38988808-45200000 |
| chr8q | chr8:45200001-144950888 |
| chr8p_1 | chr8:406428-37844896 |

| **Table S6. Gene list for calculating ModuleScore in murine wound healing, related to Figure S5.** | |
|---|---|
| **Cell subpopulation** | **Genes** |
| Monocyte | *Plbd1, Ifi209, Sirpb1c, Ltb4r1, Mcemp1, Zbp1, Ms4a4c, Plac8, Slfn1, Mmp8, F10, Gpr141, Adgre4, Ms4a8a, Hp, Klra2, Ifitm6, Chil3, Ly6c2, Cd177* |
| *Cd163*$^{+}$ macrophage | *Tslp, Hpgds, Gpr34, Stard8, Ccdc152, Abca9, Arhgap19, Cbr2, Folr2, Npl, Hal, Tmem8, Ccl8, Ednrb, Clec10a, Nxpe5, Fcrls, Ccl12, Ccl24, Cd163* |
| *Cd36*$^{+}$ *Gpnmb*$^{+}$ *Il1b*$^{-}$ macrophage | *Tmem37, Tlr7, C1qa, Ms4a7, Nxpe5, C1qb, Hpse, Mrc1, Hal, C1qc, Fcrls, Stab1, Folr2, Arhgap19, Gpr34, Gpnmb, Hpgds, Syngr1, Abcc3, Gdf15* |
| *MHCII*$^{+}$ macrophage | *Tnip3, Clec4a3, Aif1, Plbd1, Fcgr1, Clec12a, Hpgd, Sirpb1c, H2-Aa, H2-Ab1, Dpep2, Ccr2, H2-DMa, Ms4a4c, H2-Eb1, Cd72, Ciita, H2-DMb1, Clec4b1, Cx3cr1* |
| *Arg1*$^{+}$ *Il1b*$^{+}$ macrophage | *Il1a, Acp5, Adam8, Dusp4, Clec5a, Slc37a2, Msr1, Il7r, Cd36, Slc7a11, Gpnmb, Kcnn4, Rnf128, Il1rn, Cxcl3, AA467197, F10, F7, Arg1, Mmp12* |
| Neutrophil | *Csf3r, S100a9, Rdh12, Mirt1, Fpr2, Siglece, Trem3, H2-Q10, Cxcr2, Ptgs2os2, Trim30b, Trem1, Fpr1, Asprv1, Il1f9, Stfa2l1, Gm5483, Wfdc21, Retnlg, AC110211.1* |
| Dendritic cell | *Cd40, Gpr171, Adam23, Clec4b1, Mgl2, Ciita, Tspan33, Ramp3, Ccl5, Batf3, Ffar2, Dpp4, Klrk1, Cd209d, Klrd1, Klrb1b, Il4i1, Cd209a, Flt3, Ccl22* |
| T cell/ILC | *Trdc, Itgae, Trac, Trbc2, Cd5, Cxcr6, Cd3d, Cd247, Icos, Lck, Ctla4, Trbc1, Il2rb, Foxp3, Zap70, Cd6, Cd3g, Cd163l1, Tcrg-C1, Cd3e* |
| B cell | *Fcrl1, Igkc, Fcrla, Tnfrsf13c, Cxcr5, Cd79a, Bank1, Iglc1, Fcmr, Mzb1, Iglc2, Iglc3, Pou2af1, Ighd, Fcer2a, Cd19, Gm31243, Ms4a1, Vpreb3, Pax5* |
| Basophil | *Ptger3, Aqp9, Cd7, Padi2, Hgf, Il6, Rab44, Grm6, Gata2, Sytl3, Wdr95, Cd200r3, Cyp11a1, Slc18a2, Il4, Ms4a2, Cpa3, Alox15, Mcpt8, Fcer1a* |
| Mast cell | *Pclaf, Kcnn4, Cdca3, Hist1h1b, Mcm5, Mki67, Hist1h2ae, Gmnn, Clec12a, Birc5, Cdca8, Acp5, Top2a, Ube2c, Tk1, Mmp9, Cyp2s1, Ocstamp, Atp6v0d2, Slc9b2* |
| *Cd34*$^{+}$ fibroblast | *Opcml, Gpc3, Cyp7b1, Tnmd, Syt13, Col14a1, Svep1, Cadm3, Entpd2, Adam33, Adgrd1, Tnxb, Clec3b, Cyp1b1, Ccl11, Adm, Efemp1, Adamts15, Nov, Dpep1* |
| *Fmod*$^{+}$ fibroblast | *P4ha3, AW551984, Sfrp2, Capn6, Il1rl1, Ltbp1, Camk4, Kcnj15, Vegfd, Tnfsf11, Ltbp2, Igfbp2, Cxcl5, Scx, Thbs4, Ptx3, Cda, Mme, Lnx1, Egfl6* |
| *Mki67*$^{+}$ fibroblast | *Tacc3, Cdca3, Scx, Cenpe, Hmga2, Hmmr, Rrm2, Tpx2,* |

| | |
|---|---|
| | *Tnfsf11, Il1rl1, Cdkn3, Ccna2, Pthlh, Cdc20, Ccnb1, Mxd3, Gjb3, Anln, F2rl1, Il11* |
| *Lrrc15*$^+$ fibroblast | *Wisp1, Ptk7, Aldh1l2, Twist2, Ddah1, Hhipl1, Wnt5a, Stra6, Grem2, Tnn, Lgr5, Fhod3, Lrrc15, Kif26b, Spon1, Epha3, Col7a1, Tubb3, Crabp1, Prss35* |
| *Prg4*$^+$ chondrogenic progenitor | *Chad, Galnt15, Ppp1r1b, Cadm1, Pla2g5, Bmp2, Gdf5, Etv5, Omd, Corin, Cdo1, Atp1a2, Gm19935, Tmem213, 5033426O07Rik, Atp6v0a4, Notum, Tspan8, Msmp, Gm525* |
| *Igfbp5*$^+$ chondrocyte | *Galnt15, Fam124a, Penk, Myoc, Clu, Smoc1, Hrct1, Col8a2, Syn3, Adamtsl3, Ucma, Gdf5, Fbln7, Cyp26a1, Uts2r, Cpxm2, Wfdc2, Angptl7, Prss12, Cilp2* |
| Hypertrophic chondrocyte | *Pcsk6, Timp4, Kcna2, Frzb, Krt8, Lbhd2, Cmtm5, Proser2, 1190005I06Rik, Corin, Ppp1r3c, Extl1, Zfp385c, Dgat2, Wnk4, Loxl4, Col10a1, Cidea, Sox8, Adamts14* |
| *Spp1*$^+$ chondrocyte | *Hapln1, Ncmap, Lbhd2, 1520401A03Rik, Col9a2, Col11a2, Col10a1, Atp6v0a4, D630045J12Rik, Scrg1, Lcn2, Meltf, Col2a1, Serpina3n, Ucma, Notum, Trim46, Matn3, Kcna6, Tmem213* |
| Keratinocyte | *Col17a1, Krt79, Krt2, Serpinb5, Tfap2b, Clca3a2, Capns2, Mt4, S100a14, Krt15, Trim29, Gm16136, Ckmt1, Dsp, Fgfbp1, Fcgbp, Esrp1, 4631405K08Rik, Dsg1a, Lypd3* |
| Endothelial cell | *Cdh5, Adgrl4, Tie1, Sox18, C1qtnf9, Ccl21a, Mmrn2, Cldn5, Emcn, Ptprb, Grrp1, Flt4, Tmem252, Aplnr, Cyyr1, Mmrn1, Ushbp1, Podxl, Gpihbp1, Myct1* |
| Perivascular cell | *Rasl12, Ppp1r14a, Emid1, Rgs4, Tbx2, Kcnj8, Lmod1, Gucy1b1, Foxs1, Tusc5, Gucy1a1, Egflam, Abcc9, Gm13889, Vtn, Trpc6, Rgs5, Myh11, Higd1b, Olfr558* |
| Satellite cell | *Plp1, Chodl, Tnnt3, Ank1, Ckm, Acta1, Atp2a1, Myoz2, Myl1, Tcap, Cacng1, Apobec2, Tnnc2, Sln, Mb, Fitm1, Cox8b, Myf5, Smpx, Cdh15* |
